# Key molecular and cellular events of table olive fruit abscission zone formation during natural maturation and after ethephon treatment

**DOI:** 10.1101/2025.11.10.687704

**Authors:** Minmin Wang, Emily Santos, Shaina Eagle, Alisa Chernikova, Shuxiao Zhang, Phuong Tran, Giulia Marino, Judy Jernstedt, Franz Niederholzer, Becky Wheeler-Dykes, Thomas Wilkop, Louise Ferguson, Georgia Drakakaki

## Abstract

Abscission zones are specialized cell layers that enable organ detachment, yet their morphology and regulation vary among species. Olive (*Olea europaea*), a non-climacteric fruit, requires high concentrations of ethylene-releasing compounds to reduce fruit removal force (FRF) for mechanical harvest. To elucidate mechanisms of fruit abscission zone (FAZ) development in olive, we integrated physiological, transcriptomic, and cellular analyses during natural maturation and after ethephon treatment. Olive fruits emitted low ethylene at color transition, coinciding with declining FRF, and application of 1500 ppm ethephon reduced FRF after one week. Transcriptome analyses of FAZ tissues, using a mesocarp-subtraction strategy to isolate FAZ-specific responses, identified 733 FAZ-specific genes shared between natural maturation and ethephon treatments, including genes of β-1,3-glucanases, pectate lyases, and the phenylpropanoid pathway. Microscopy revealed lignification and alkalization of FAZ, accompanied by reduced low-methylesterified homogalacturonan and non-fucosylated xyloglucan. Increased glucanase activity and reduction in plasmodesmata callose likely facilitate cellular communication during abscission. Further, this study provides the first evidence of FAZ alkalization and implicates transporter upregulation in pH regulation preceding abscission. Our findings advance understanding of abscission biology in non-climacteric fruits and delineate conserved features of FAZ development.

**Summary Statement:** Integrated physiological, transcriptomic, and cellular analyses reveal that fruit abscission zone development in olive involves lignification, alkalization, and coordinated cell wall remodeling, identifying β-1,3-glucanases, pH-regulating transporters, and pectate lyases that facilitate cell separation in a non-climacteric fruit.

## 1. Introduction

Abscission zones are predetermined positions where plant organ detachment takes place (Roberts, Elliott and Gonzalez-Carranza, 2002). The abscission zone typically consists of one or several cell layers that are not only morphologically distinct from surrounding tissues, but also respond differently to phytohormone signals from neighboring cells (Osborne and Morgan, 1989; Roberts, Elliott and Gonzalez-Carranza, 2002). The Arabidopsis flower abscission zone has been the most extensively characterized for morphology, genetics, and phytohormone regulation, serving as the primary model for plant organ abscission (Patharkar and Walker, 2018). However, the morphology and location of abscission zones vary greatly among plant species and organs (Pautot, Crick and Hepworth, 2025). The fruit abscission zone (FAZ), though of high economic importance for crop harvesting and postharvest fruit quality, has been studied to a lesser extent, primarily in selected crop species (e.g., tomato, citrus, pepper (Merelo *et al*., 2017; Hill *et al*., 2023; Pautot, Crick and Hepworth, 2025)), with no single model system fully representing the structural and developmental diversity of fruit abscission zones across various crop species. The development of the FAZ is also tightly regulated by phytohormones and coordinated with fruit maturation (Roberts, Elliott and Gonzalez-Carranza, 2002; Xie *et al*., 2013; Pautot, Crick and Hepworth, 2025).

Fruits are categorized as climacteric or non-climacteric based on their CO□ and ethylene release patterns during maturation and ripening (Kou *et al*., 2021). With ethylene being a key regulator of fruit abscission (Taylor and Whitelaw, 2001), differences in ethylene physiology influence FAZ development in these two fruit categories (Ayub, Pessenti and Pereira, 2020; Pujol and Garcia-Mas, 2023). With the exception of citrus, the characterization of FAZs in non-climacteric fruits remains very limited. Ethylene treatment induces lignification of the fruit abscission zone and upregulates genes in the phenylpropanoid pathway. It also alters pectin composition through the upregulation of polygalacturonase (PG) and pectate lyase (PL) (Merelo *et al*., 2017).

The reduction of fruit removal force is of high economic value in the table olive industry, as there is a negative correlation with fruit removal force and mechanical harvest efficiency (Zipori *et al*., 2014), overall benefiting tree health. Olive is a non-climacteric fruit, with low ethylene evolution and no distinct rise in CO_2_ levels at the mature or ripening stage (Ben-Tal and Lavee, 1976; Lavee and Martin, 1981). The developmental decrease in fruit removal force (FRF) during the harvest season has been consistently observed in olives across the literature, with premature olives showing a FRF over 400 g and mature olives near harvest showing a FRF around 200 g (Blanco-Roldan *et al*., 2009; Zipori *et al*., 2014; Alowaiesh, Singh and Kailis, 2016). Olive requires high concentrations of exogenous ethylene-releasing compounds to reduce the fruit removal force at the pedicel. The effective treatment concentrations lie between 400 and 2000 ppm ethylene equivalents (Ben-Tal and Lavee, 1976; Rugini, Bongi and Fontanazza, 1982; Burns *et al*., 2008; Goldental-Cohen *et al*., 2017). Among the various olive fruit loosening agents tested over many years, the ethylene-releasing agents, 1-aminocyclopropane-1- carboxylic acid (ACC) and ethephon, remain the only effective chemicals in reducing FRF (Burns *et al*., 2008; Castro-Garcia and Ferguson, 2017).

Cellular and molecular studies of the olive FAZ have revealed several features associated with fruit abscission. Lignin deposition has been linked to FAZ development in the ‘Manzanillo’ cultivar (Reed and Hartmann, 1976). Comparative analyses of callose, pectin, hemicellulose, and extensin levels in pre-abscission and post-abscission FAZ tissues have further highlighted the dynamic remodeling of the cell wall during separation (Parra and Gomez-Jimenez, 2020; Parra *et al*., 2020). Transcriptome analysis of mature versus ripe olive FAZ (154 vs 217 day-post-anthesis) identified a β-1,3-glucanase gene and multiple phenylpropanoid pathway genes among the most upregulated genes (Gil-Amado and Gomez-Jimenez, 2013). Additionally, transcriptomic profiling of FAZ and leaf abscission zones under ethephon treatment revealed increased expression of genes encoding expansins and polygalacturonases (PGs) (Goldental-Cohen *et al*., 2017). In non-climacteric fruits such as citrus or pepper, lignification is a prominent feature of cell wall modification during FAZ development (Merelo *et al*., 2017; Hill *et al*., 2023). However, other cellular and molecular changes of major cell wall polymers, such as pectins and hemicelluloses, remain insufficiently characterized as defining events in olive FAZ development.

A decline in FRF occurs during both natural maturation and after ethephon treatment, reflecting the development of the FAZ. In order to understand the cellular, molecular, and anatomical changes underlying this process, we comprehensively characterized the olive FAZ during natural maturation and in response to ethylene stimulation. We compare immature and pre-abscission mature olive FAZ, based on growing degree days (GDD) and FRF values, in the table olive cultivar “Manzanillo”, and examine the common changes at the FAZ induced by both ethephon treatment and natural maturation. Cell wall modifications were examined at both cellular and molecular levels through immunohistochemistry and transcriptome profiling. By identifying structural features and candidate genes associated with FAZ differentiation that are common to both natural and ethephon-induced abscission, this study provides molecular and cellular markers useful for evaluating fruit loosening agents, guiding cultivar improvement, and understanding environmental modulation of fruit abscission.

## 2. Material and Methods

### 2.1 Plant material and growth conditions

Table olive trees (*Olea europaea* L.), cultivar ’Manzanillo’, were located at Nickels Soil Laboratory (Arbuckle, CA, USA). The trees were planted in 2001, in a matrix of 13 rows by 30 trees, with a spacing of 3.66 m x 5.49 m. The 7^th^ row, consisting of the cultivar ‘Sevillano’, serves as pollinizer. Trees were drip irrigated several times per week to match evapotranspiration. Adequate levels of tree mineral nutrition were confirmed with annual leaf analyses taken in July of each year in the study.

Day 0 for growing degree day calculations was marked by the 50% bloom time on April 25, 2022. Weather data was obtained from the on-site weather station and accessed via the website <u>ipm.ucanr.edu</u>.

Growing degree days (GDDs, °C) were calculated using the formula, 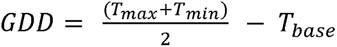, with the T_base_ being 15°C for table olive fruit growth (Pérez-López *et al*., 2008). Mechanical harvest was performed with a trunk-shaker on September 22, 2022, at 1320 GDD, a time near commercial harvest.

### 2.2 Fruit removal force measurement and ethephon treatment

The fruit removal force (FRF) was measured over a period of five dates during pre-harvest. The force required for detaching the olive fruit from the pedicel was recorded by a force tester (Model DS2-11, IMADA, Inc., Northbrook, IL). Ethephon treatment was performed by submerging branches of 24 unharvested trees on October 17, 2022. The selection criterion for the branches was the presence of more than 12 olives per branch, out of which a minimum of four olives showed an FRF over 300 g. Two of the aforementioned branches from each tree were then either dipped into 1500 ppm ethephon (Ethrel^TM^, Bayer Crop Science, St Louise, MO) or water as a control for 30 sec. Both treatments contained 0.025% surfactant (Activator 90, Loveland Products, Inc., Loveland, CO). Ethrel contains 21.7% ethephon in acidic form and reacts with water as pH increases, generating ethylene. The FRFs were measured one week after treatment (October 24, 2022), and samples were collected for transcriptome analysis. Repetition of the ethephon treatment and the same FRF measurement regime were performed in 2023, starting on October 16^th^, 2023.

### 2.3 Fruit ethylene emission and respiration rate measurements

Ten olive fruits were placed in a 460-mL airtight glass jar with a rubber-sealed lid and resealable headspace gas sampling tubing, representing one biological replicate. Four biological replicates were measured for ethylene and carbon dioxide (CO□) emission on five dates during the pre-harvest period. After a 1-hour equilibration period at room temperature, a 1-mL headspace gas sample was drawn and analyzed for ethylene concentration using gas chromatography with a flame ionization detector (Model Carle AGC-211, EG&G Chandler Engineering, Tulsa, OK), as described (Villalobos-Acuña *et al*., 2010). For respiration rate measurements quantifying carbon dioxide concentrations, a 10-mL headspace gas sample was injected into an infrared gas analyzer (Model PIR-2000R, Horiba Instruments Inc., Irvine, CA). Ethylene and CO□ concentrations were determined by comparing sample responses to authentic gas standards (Airgas, Radnor, PA).

### 2.4 Transcriptome analysis

Olives were collected based on the FRF of four fruits per branch. If 4 out of 12 olives showed the desired FRF as described below, the remaining 8 olives on the same branch were collected as one biological replicate for transcriptome analysis. The samples were separated into three categories: 1) Green immature samples (abbreviated as G) collected from branches where the adjacent fruit exhibited an FRF higher than 300 g (September 12, 2022). 2) Horticulturally mature samples (abbreviated as M) collected from branches where adjacent fruit showed a FRF lower than 200 g (October 24, 2022), and 3) ethephon-treated samples (abbreviated as E) from branches where adjacent fruit exhibited a FRF lower than 200 g (when?). Abscission zone tissue samples (abbreviated as AZ) were collected from the region connecting pedicel and fruit, mesocarp tissue samples (abbreviated as C) were collected from the adjacent regions. All samples, having a 1 mm^3^ volume, were snap-frozen with liquid nitrogen. The six collected sample categories are: green immature abscission zone (GAZ), horticulturally mature abscission zone (MAZ), ethephon-treated abscission zone (EAZ), green immature mesocarp (GC), horticulturally mature mesocarp (MC), ethephon-treated mesocarp (EC). Four biological replicates of the six groups (GAZ, MAZ, EAZ, GC, MC, EC) were used for RNA extraction.

RNA of the above samples was extracted using the MagMAX Plant RNA Isolation Kit (Applied Biosystems, Waltham, MA) and sent for library construction, transcriptome sequencing analysis, and data analysis (Novogene Corporation Inc., Sacramento, CA). During the analysis, reads were mapped to the OLEA9 genome assembly of *Olea europaea* subsp*. europaea* (common olive) (Cruz *et al*., 2016) by HISAT2 (Kim *et al*., 2019). The abundance of transcripts in each sample was quantified by featureCounts (Liao, Smyth and Shi, 2014) and normalized by fragments per kilobase of transcript sequence per million mapped reads (FPKM). Differential gene expression (DGE) analysis of GAZ vs MAZ, GAZ vs EAZ, GC vs MC, and GC vs EC was performed by DESeq2 (Love, Huber and Anders, 2014). The negative binomial distribution was used for the *p*-value calculation, and the screening threshold for differentially expressed genes (DEG) was set to |log_2_(FoldChange)| ≥ 1 with an adjusted *p*-value (padj) of ≤ 0.05. Candidate gene lists derived from intersections of two DGE analysis sets were generated using the INDEX and MATCH functions in Microsoft Excel. Up- and down-regulation in DGE analysis were processed through subset operations separately and then combined.

Real-time PCR was performed to validate DEGs selected from the transcriptome analysis. cDNA was synthesized using the High-Capacity cDNA Reverse Transcription kit (ThermoFisher Scientific, Waltham, MA), using the RNA mentioned above as template. The primers targeting these genes are listed in **Supplementary Figure 1**. Quantitative analysis was performed using the PowerSYBR Green PCR Master Mix (ThermoFisher Scientific) with the ABI7000 cycler (ThermoFisher Scientific) using default settings. The abundance of transcripts was determined by the relative quantification method (Schmittgen and Livak, 2008). In the relative quantification, *EIF4A* (OLEA9_A107450) was used as a reference (Tang *et al*., 2019), since it has a low coefficient of variance of 0.063 in this transcriptome analysis.

### 2.5 Glucanase activity assay

FAZ tissue of control and ethephon-treated samples was collected before the treatment (day 0), day 2, and day 7 after treatment, and snap-frozen in liquid nitrogen. Approximately 1 g of frozen FAZ tissue was homogenized in liquid nitrogen using mortars and pestles, and resuspended in 50 mM HEPES buffer (pH 7.5) containing 1 mM PMSF in a 1:1 w/v ratio. The homogenates were then filtered through Miracloth (MilliporeSigma, Burlington, MA) and subsequently cleared by 1000*g* centrifugation at 4°C for 2 min. The clear upper phase of the lysate was desalted by size exclusion chromatography using a PD-10 prepacked column (Cytiva, Marlborough, MA) with assay buffer (50 mM TrisHCl, pH 8.5) as eluent. The protein content was measured by the Bradford protein assay (Bradford, 1976), and extracted proteins were used for the glucanase assay. Background sugar levels are estimated via the boiled protein reaction.

Glucanase activity assays were performed using the dinitrosalicylic acid (DNS) method (Miller, 1959), in which the DNS reagent is composed of 1% 3,5-dinitrosalicylic acid, 0.05% sodium sulfite, and 1% sodium hydroxide in water. A 100 μL assay mixture contained 50 μL desalted crude extract, 20 μL of DI water, and 30 μL laminarin (TCI America, Portland, OR), yielding a final substrate concentration of 15 g/L. Stock laminarin had a concentration of 50 g/L. Assays were performed at 50°C for 45 min, terminated with 900 μL DNS reagent at 85°C for 10 min, then 170 μL 40% potassium sodium tartrate was added to stabilize the color before measuring the absorbance at 575 nm on a spectrophotometer (UV-1700, Shimadzu, Kyoto, Japan). The standard curve was built by dissolving 3.125 to 100 μg of glucose in the assay buffer. Background levels of reduced sugars in the assay were determined using boiled protein extracts as references.

### 2.6 Histochemical staining

Collected GAZ, MAZ, and EAZ tissue samples, of 1 mm^3^ volume, were sectioned at a thickness of 50 μm with a vibratome (VT1000S, Leica Biosystems, Deer Park, IL). For live staining and imaging within 24 hours, the sections were kept in PBS (137 mM sodium chloride, 2.7 mM potassium chloride, 10 mM disodium phosphate, and 1.8 mM monopotassium phosphate). Samples for histochemical analysis were preserved in 4% paraformaldehyde (PFA) with 0.1% Tween-20 in PBS solution.

Live staining of callose was performed by aniline blue fluorochrome (100-1, Biosupplies, Australia) using a concentration of 80 μg/mL in PBS for 5 min, and then washing with PBS before imaging. Live staining of 2’,7’-Bis-(2-Carboxyethyl)-5-(and-6)-Carboxyfluorescein, Acetoxymethyl Ester (BCECF, AM, Invitrogen, Carlsbad, CA) was performed using a concentration of 10 μM in PBS for 15 min in the dark at room temperature, before imaging. Stock solution of BCECF, AM was prepared in anhydrous DMSO at 10 mM according to the manufacturer’s instructions.

Fuchsin staining was performed according to a previously published protocol (Ursache *et al*., 2018) with sections prepared in ClearSee solution (Kurihara *et al*., 2015). Cellulose was stained by Calcofluor White Stain at a concentration of 0.22 g/L (Sigma-Aldrich, St Louis, MO) for identification of the cell profiles (Ursache *et al*., 2018). The immunostaining procedures generally followed the manufacturers’ instructions for the Alexa Fluor™ 488 Tyramide SuperBoost™ Kit (Thermo Fisher Scientific) with adjustments according to established protocols (Gerttula and Groover, 2017). Sections fixed in PFA solution were washed three times in PBT buffer (PBS plus 0.2% Tween-20) and incubated with primary antibody diluted in goat serum overnight. For callose immunostaining, anti β-1,3-glucan monoclonal mousez antibody (BS-400-2, Biosupplies, Australia) was diluted at a 1:1000 ratio in blocking buffer (10% goat serum, Tyramide SuperBoost™ Kit). For CCRC series mouse monoclonal antibodies (M1 and M101, (Pattathil *et al*., 2010)), a 1:5 dilution was used. Primary antibodies were detected with goat anti-mouse IgG secondary antibody using the Alexa Fluor™ 488 Tyramide SuperBoost™ Kit. High and low-methylesterified homogalacturonans were targeted by JIM7 and JIM5 primary antibodies (Knox *et al*., 1990), using a 1:10 dilution, and visualized with Goat anti-Rat IgG Secondary Antibody, Alexa Fluor™ 488 (A-11006, ThermoFisher Scientific) at a 1:50 dilution.

### 2.7 Microscopy

All images were acquired on a ZEISS LSM 980 confocal microscope with Airyscan 2 (Carl Zeiss Microscopy, Jena, Germany). Live staining of BCECF and aniline blue fluorochrome was recorded in confocal mode with a 1 AU pinhole, and other images were taken in SR-8Y Airyscan mode. All images were analyzed in ZEN Lite (Version 3.7) using the built-in “Measure” functions. The display of each channel in each figure was adjusted to the same white point value for comparison before exporting to TIFF files. Figures were assembled and cataloged in Microsoft PowerPoint.

Samples stained for BCECF analysis were imaged with a 10X 0.45 numerical aperture (NA) objective in three channels, with the chlorophyll autofluorescence channel using 639 nm excitation at 5% laser power and emission collection over 640-692 nm; the BCECF channel used excitation at 488 nm at 5% laser power and emission collection over 492-658 nm. Transmitted light images were collected with the forward signal from the 639 nm at 2% laser power and the transmitted light detector. 23 µm Z-stacks with 12 slices were taken, and a maximum intensity projection was generated prior to exporting the image and performing data analysis. Immature and ethephon-treated samples were further examined at higher resolution using a 40X 1.2 NA water immersion objective and a scan zoom of 4. Aniline blue fluorochrome-stained samples were imaged in two channels, with the aniline blue fluorochrome channel using 405 nm excitation at 1% laser power and emission collection over 413-605 nm the transmitted light channel used the forward signal from the 639 nm laser at 2% power.

Airyscan mode imaging was used for Fuchsin staining and all immunofluorescence staining. Fuchsin staining was imaged with a 20X 0.8 NA objective, using 561 nm excitation at 5% laser power. Alexa Fluor™ 488 conjugated secondary antibodies (anti-mouse for CCRC-M1/M101 and BS-400-2; anti-rat for JIM5/JIM7) were imaged with a 40X 1.2 NA water immersion objective, using 488 nm excitation at 5% laser power. The counterstaining Calcofluor White was imaged at a second channel using 405 nm excitation at 5% laser power. Calcofluor White channel was set to display a signal intensity range of 0- 3000 arbitrary units in ZEN Lite before images were exported to TIFF format, unless otherwise specified in the figure legends.

## 3. Results

### 3.1 Physiological changes during the pre-harvesting period

Previous studies measuring olive fruit ethylene emission at the mature stage suggest that olive is a non-climacteric fruit (Ben-Tal and Lavee, 1976; Lavee and Martin, 1981). Non-climacteric fruits lack the autocatalytic increase in ethylene production in response to exogenous ethylene during maturation (Kubo, 2015). Consequently, efforts to reduce FRF and facilitate mechanical harvest have relied on high concentrations of ethylene-releasing compounds (400–2000 ppm). To further validate the olive fruit ethylene physiology, we monitored ethylene emission and respiration during the pre-harvest period. Very low ethylene emission (<0.1 μL/kg/h) and no respiration burst were observed (**Figure 1A, B**), when fruit color changed from green to pale yellow (**Figure 1**), reaching horticultural maturity at around 1300 GDD (150 dpa). The ethylene emissions and respiration rates observed in this trial are consistent with previous reports (Ben-Tal and Lavee, 1976; Lavee and Martin, 1981), displaying a non-climacteric fruit pattern.

**Figure 1.**
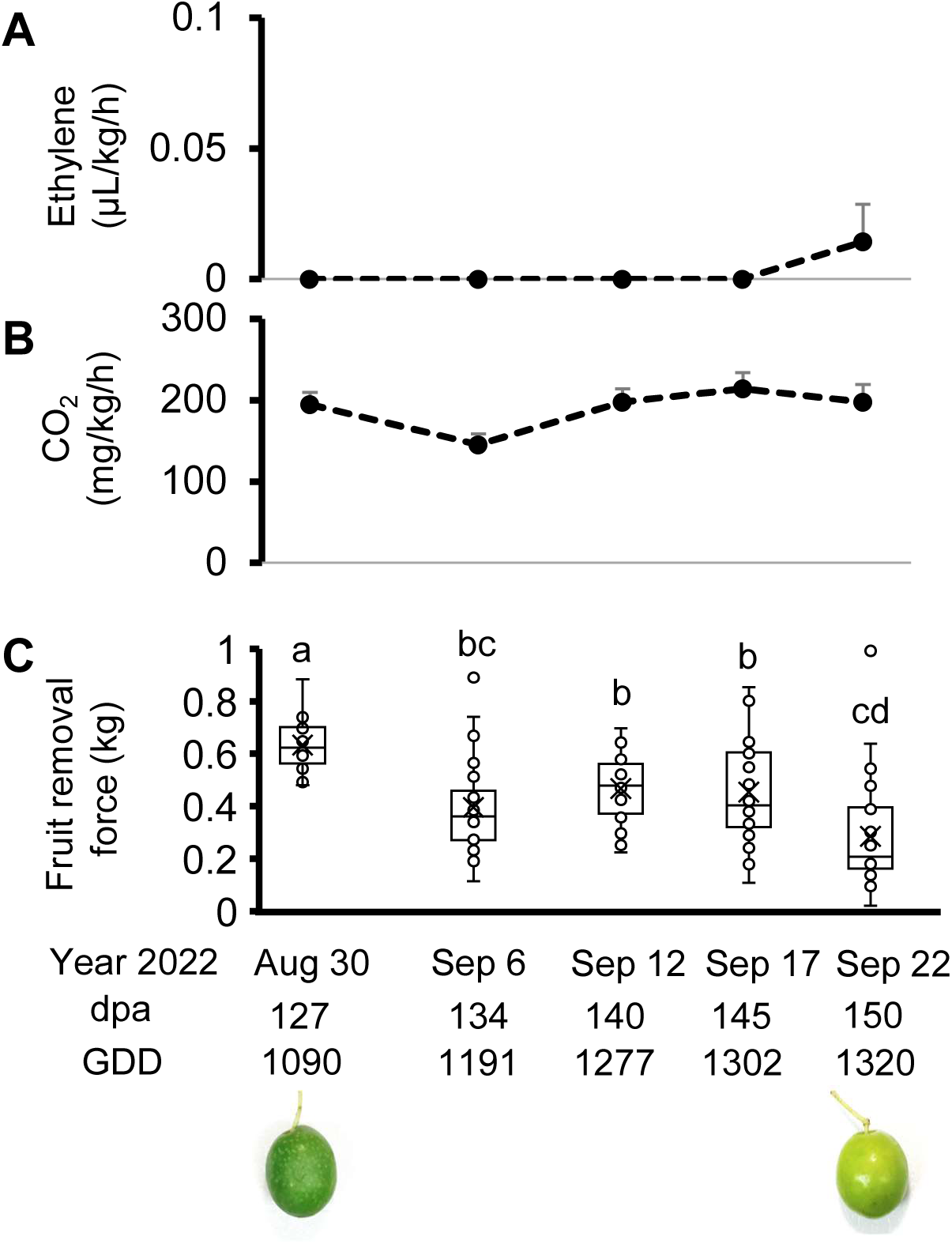
Ethylene emission, respiration rate, and fruit removal force of table olive fruits during the pre-harvest season. **(A)** Ethylene emission. Each biological replicate contains ten fruits of one branch (n = 4 trees). Error bars represent standard error. **(B)** Respiration rate as carbon dioxide emission is shown. Each biological replicate contains ten fruits of one branch per tree (n = 4 trees). Error bars stand for standard error. **(C)** Fruit removal force (n = 60 fruits from six trees). Different letters indicate significant difference between groups (one-way ANOVA, Tukey HSD, p < 0.05). ‘dpa’ stands for day-post-anthesis. Lines of box plots, from top to bottom, show the maximum, the third quartile, the median, the first quartile, and the minimum, with x marking the mean. Box plots in all figures are presented employing this format.

FRF showed a decreasing trend through the pre-harvest period (**Figure 1C**), which started in early September (∼1200 GDD) until late September (∼1300 GDD). The start of detectable ethylene emission in olive fruits (**Figure 1C**) coincides with the second significant decrease of FRF in late September.

### 3.2 Identification of the fruit abscission zone

We investigated several possible detachment regions in table olive, including the fruit abscission zone (FAZ), the pedicel abscission zone (PAZ), the leaf abscission zone (LAZ), and the secondary abscission zone (SAZ) (**Figure 2A**) as the site of fruit separation. The FAZ was the dominant detachment site, observed in over 85% of the fruit, harvested with a trunk shaker (**Figure 2B, C**). **Figure 2D–G** shows a stepwise magnification of the FAZ. The site of pedicel detachment from the fruit at abscission is indicated by a black dashed line (**Figure 2F),** while the region used in the cellular and transcriptome analysis throughout the manuscript is marked by a red square. The cell profile of the FAZ underneath the calyx, stained with Calcofluor White, is shown in **Figure 2G**. Notably, the cell size in the FAZ is increasing in the proximal-to-distal direction.

**Figure 2.**
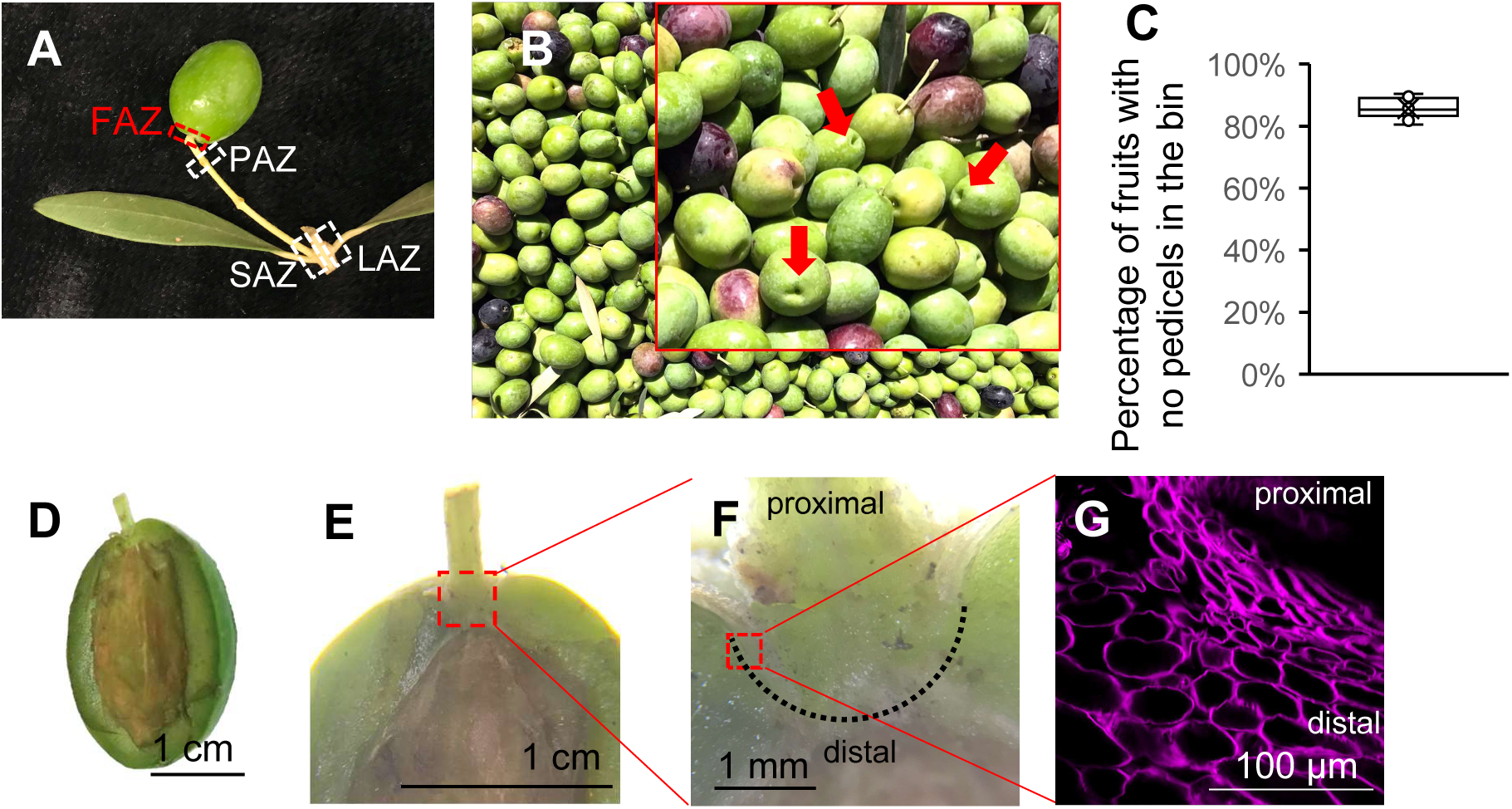
Identification of major abscission zones in table olive mechanical harvest. **(A)** Locations of abscission zones for fruit and leaf detachment. FAZ, fruit abscission zone. PAZ, pedicel abscission zone. SAZ, secondary abscission zone. LAZ, leaf abscission zone. **(B)** Representative image from a harvester collection container. Red arrows indicate fruit detachment at the fruit abscission zone (FAZ). **(C)** Harvester containers were imaged, and fruits with no pedicles were manually counted (n = 12 bins). (**D - G**) Zoom-in illustration highlighting the region of interest used to examine cellular changes at the FAZ. (**G**) Calcofluor white staining showing FAZ cell profiles.

### 3.3 Ethephon treatment induced a reduction in the fruit removal force

Over two consecutive growing seasons, we examined the relationship between the FRF and the application of 1500 ppm ethephon. A consistent, significant reduction of fruit removal force was observed after one week of treatment (n = 120, *p* < 0.05, t-test). Controls showed an FRF over 300 g, and treated fruits showed an FRF of around 200 g (**Figure 3**). Given the observed decrease in the FRF, we hypothesize that ethephon application induces significant anatomical, compositional, and molecular changes in the FAZ that contribute to this reduction.

**Figure 3.**
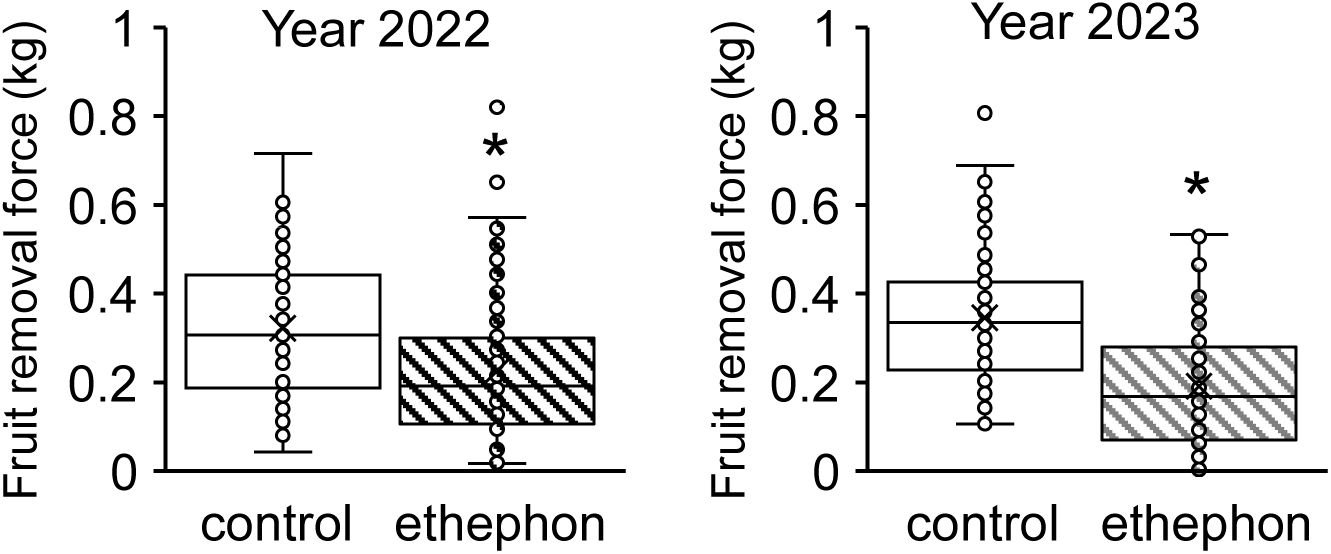
Ethephon induced FRF reduction. FRF of ethephon-treated fruits in 2022 (**A**) and 2023 (**B**) growing seasons. Branches containing four middle fruits of over 300 g FRF were submerged in 1500 ppm ethephon solution or water for 30 sec, and FRF was measured after one week (n = 120 fruits from 12 trees). The ethephon immersion experiment was repeated in 2023. Asterisk indicates significant difference from the control (t-test, p < 0.05, n = 120).

### 3.4 Transcriptome analysis reveals key genes driving olive FAZ development

Although ethylene has been used as a fruit loosening agent (Burns *et al*., 2008), leading to a reduction of the FRF, the mechanisms by which the application affects the development of the FAZ are not understood. Towards establishing specific gene expression changes in the FAZ during natural maturation and after ethephon treatment, we performed transcriptome analysis, providing cues on mechanisms altering the FAZ. Separate mesocarp samples were included for background elimination, in order to prevent the mesocarp cell wall remodeling events dominating the transcriptome data (**Figure 4A**). Samples were harvested based on GDD for developmental stages and further selected by FRF (**Figure 4B**). Over 80% unique reads were mapped to the OLEA9 genome assembly of *Olea europaea* subsp*. europaea* (common olive) (Cruz *et al*., 2016). Out of the total 82505 olive genes, 54335 genes were detected in this transcriptome analysis. The transcriptome dataset is accessible through the NCBI Sequence Read Archive (BioProject accession number: PRJNA1335555).

**Figure 4.**
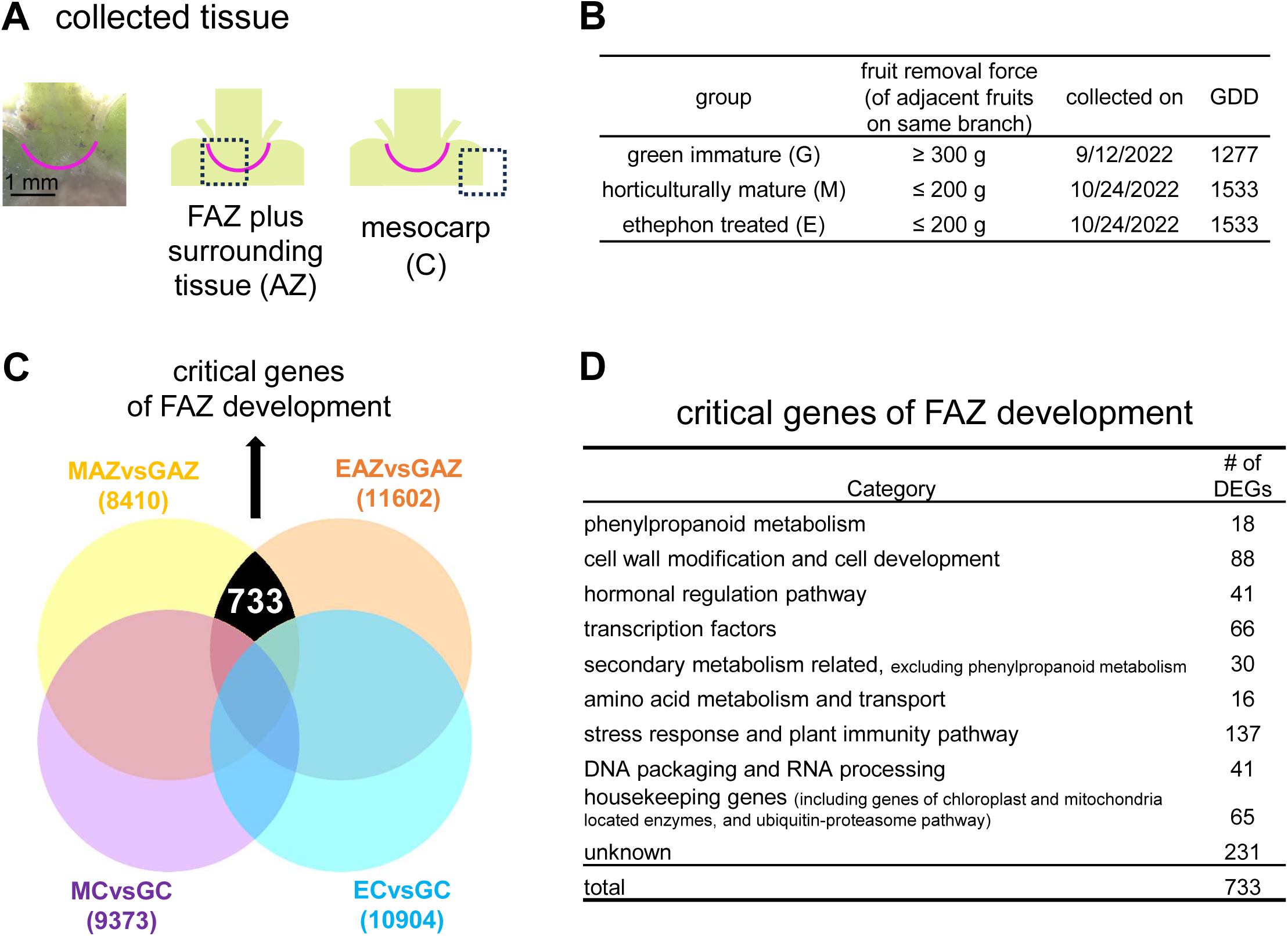
Transcriptome analysis of the olive fruit abscission zone. A combination of two types of tissue collected (**A**) and three different developmental stage/treatment groups (**B**) were used. (**A**) A 1 mm^3^ region flanking the fruit abscission zone was collected as the abscission zone tissue [AZ]; mesocarp [C] tissue was collected from the edge of the AZ region. The magenta line indicates the fruit abscission zone. (**B**) Green immature [G], horticulturally mature [M], and ethephon-treated [E] fruit samples with designated fruit removal force were collected on the indicated GDD. Ethephon-treated fruits were treated by ethephon on 10/17/22, with adjacent olives of FRF ≥300 g. (**C**) Identification of critical gene candidates of fruit abscission zone cell wall remodeling via a subset subtracting strategy. Abbreviation of treatment group and tissue: G = Green Immature; M = Horticulturally Mature; E = Ethephon treated; AZ = Fruit Abscission Zone; C = Mesocarp. The six combinations of treatment/development group and tissue are GAZ, MAZ, EAZ, GC, MC, and EC. Each DGE analysis (denoted as “vs”) uses a *p*-value threshold of 0.05 by comparison of the FPKM value. The number in parentheses indicates the number of DEGs identified in each DGE analysis. The solid black region represents the 733 DEGs (**C_int_**) identified after applying the formula “{(MAZvsGAZ) – [(MAZvsGAZ)∩(MCvsGC)]}∩{(EAZvsGAZ) – [(EAZvsGAZ)∩(ECvsGC)]}”. (**D**) Categorization of the 733 candidate genes identified from (**C**), based on Uniprot annotated biological function.

In order to find critical genes in abscission zone development, common to horticulturally mature and ethephon-induced abscission zones, while eliminating mesocarp background, the following formula was used:

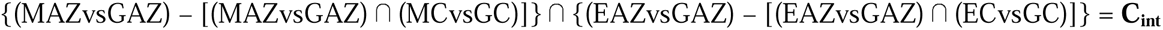

The term “vs” represents one differential gene expression (DGE) analysis (adjusted p-value < 0.05), where G, M, and E refer to green immature, horticulturally mature, and ethephon-treated samples, respectively; AZ refers to FAZ plus surrounding tissue; and C refers to mesocarp tissue.

The intersection (black region in **Figure 4C**) presents the set of genes that are significantly altered in both MAZ and EAZ relative to GAZ, after subtracting those changing in the mesocarp (MCvsGC and ECvsGC). This subtraction approach can be viewed as a simplified, two-node analogue of weighted correlation network analysis (WGCNA; (Langfelder and Horvath, 2008)), in which the pairwise correlation and soft thresholding procedures are replaced by DGE analysis filtering with a 2-fold change and an adjusted *p* < 0.05, to identify the mesocarp responsive genes. An intersection total of 733 genes (**C_int_**) was identified, annotated using Uniprot, and manually categorized for their biological roles (**Figure 4D**). We postulate that these 733 genes make up the critical players in the development of the olive FAZ during both natural maturation and after ethephon treatment (**Figure 4C, D**). Notably, we identified DEGs involved in phenylpropanoid metabolism (**Table 1**) and cell wall modification (**Table 2, 3, 4**). The remaining ones are implicated in various processes, including hormonal regulation, stress responses, amino acid metabolism and transport, listed in **Supplemental Table 1-8**, while 231 DEGs are unattributed (**Supplemental Table 9**). Real-time qPCR of selected DEGs validated the expression trends (**Supplemental Figure 1**).

**Table 1.**
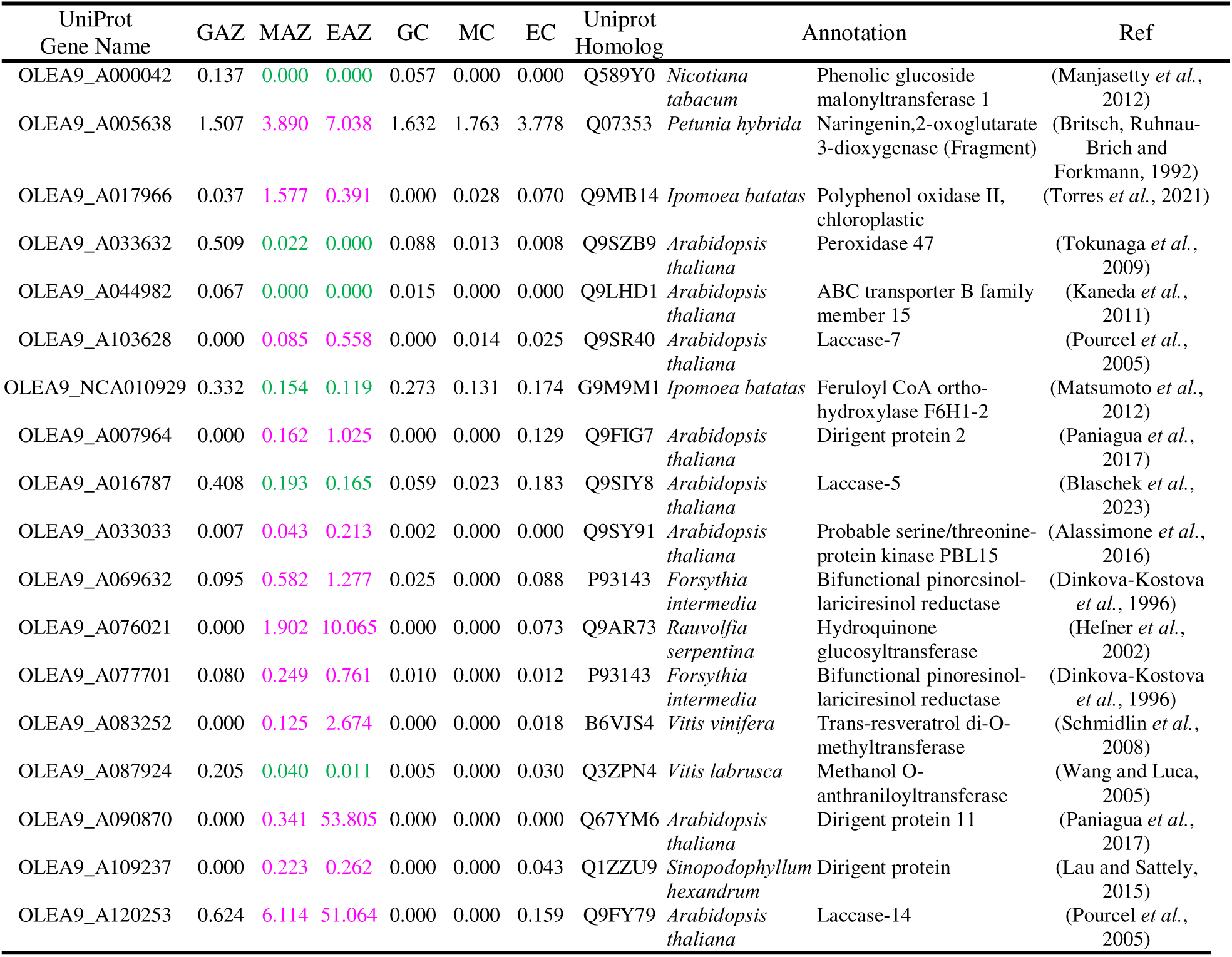
Phenylpropanoid metabolism-related DEGs in the list of critical genes (C_int_) of FAZ formation. The expression level of each group is indicated by FPKM. Magenta color denotes upregulation in MAZ and EAZ compared to GAZ, and green denotes downregulation (t-test, *p* < 0.05, n = 4). Abbreviations are listed in the legend of **Figure 4**.

| UniProt<br>Gene Name | GAZ | MAZ | EAZ | GC | MC | EC | UniProt<br>Homolog | Annotation | Ref |
| --- | --- | --- | --- | --- | --- | --- | --- | --- | --- |
| OLEA9_A000042 | 0.137 | 0.000 | 0.000 | 0.057 | 0.000 | 0.000 | Q589Y0 *Nicotiana tabacum* | Phenolic glucoside malonyltransferase 1 | (Manjasetty *et al.*, 2012) |
| OLEA9_A005638 | 1.507 | 3.890 | 7.038 | 1.632 | 1.763 | 3.778 | Q07353 *Petunia hybrida* | Naringenin,2-oxoglutarate 3-dioxygenase (Fragment) | (Britsch, Ruhnau-Brich and Forkmann, 1992) |
| OLEA9_A017966 | 0.037 | 1.577 | 0.391 | 0.000 | 0.028 | 0.070 | Q9MB14 *Ipomoea batatas* | Polyphenol oxidase II, chloroplastic | (Torres *et al.*, 2021) |
| OLEA9_A033632 | 0.509 | 0.022 | 0.000 | 0.088 | 0.013 | 0.008 | Q9SZB9 *Arabidopsis thaliana* | Peroxidase 47 | (Tokunaga *et al.*, 2009) |
| OLEA9_A044982 | 0.067 | 0.000 | 0.000 | 0.015 | 0.000 | 0.000 | Q9LHD1 *Arabidopsis thaliana* | ABC transporter B family member 15 | (Kaneda *et al.*, 2011) |
| OLEA9_A103628 | 0.000 | 0.085 | 0.558 | 0.000 | 0.014 | 0.025 | Q9SR40 *Arabidopsis thaliana* | Laccase-7 | (Pourcel *et al.*, 2005) |
| OLEA9_NCA010929 | 0.332 | 0.154 | 0.119 | 0.273 | 0.131 | 0.174 | G9M9M1 *Ipomoea batatas* | Feruloyl CoA ortho-hydroxylase F6H1-2 | (Matsumoto *et al.*, 2012) |
| OLEA9_A007964 | 0.000 | 0.162 | 1.025 | 0.000 | 0.000 | 0.129 | Q9FIG7 *Arabidopsis thaliana* | Dirigent protein 2 | (Paniagua *et al.*, 2017) |
| OLEA9_A016787 | 0.408 | 0.193 | 0.165 | 0.059 | 0.023 | 0.183 | Q9SIY8 *Arabidopsis thaliana* | Laccase-5 | (Blaschek *et al.*, 2023) |
| OLEA9_A033033 | 0.007 | 0.043 | 0.213 | 0.002 | 0.000 | 0.000 | Q9SY91 *Arabidopsis thaliana* | Probable serine/threonine-protein kinase PBL15 | (Alassimone *et al.*, 2016) |
| OLEA9_A069632 | 0.095 | 0.582 | 1.277 | 0.025 | 0.000 | 0.088 | P93143 *Forsythia intermedia* | Bifunctional pinoresinol-lariciresinol reductase | (Dinkova-Kostova *et al.*, 1996) |
| OLEA9_A076021 | 0.000 | 1.902 | 10.065 | 0.000 | 0.000 | 0.073 | Q9AR73 *Rauvolfia serpentina* | Hydroquinone glucosyltransferase | (Hefner *et al.*, 2002) |
| OLEA9_A077701 | 0.080 | 0.249 | 0.761 | 0.010 | 0.000 | 0.012 | P93143 *Forsythia intermedia* | Bifunctional pinoresinol-lariciresinol reductase | (Dinkova-Kostova *et al.*, 1996) |
| OLEA9_A083252 | 0.000 | 0.125 | 2.674 | 0.000 | 0.000 | 0.018 | B6VJS4 *Vitis vinifera* | Trans-resveratrol di-O-methyltransferase | (Schmidlin *et al.*, 2008) |
| OLEA9_A087924 | 0.205 | 0.040 | 0.011 | 0.005 | 0.000 | 0.030 | Q3ZPN4 *Vitis labrusca* | Methanol O-anthraniloyltransferase | (Wang and Luca, 2005) |
| OLEA9_A090870 | 0.000 | 0.341 | 53.805 | 0.000 | 0.000 | 0.000 | Q67YM6 *Arabidopsis thaliana* | Dirigent protein 11 | (Paniagua *et al.*, 2017) |
| OLEA9_A109237 | 0.000 | 0.223 | 0.262 | 0.000 | 0.000 | 0.043 | Q1ZZU9 *Sinopodophyllum hexandrum* | Dirigent protein | (Lau and Sattely, 2015) |
| OLEA9_A120253 | 0.624 | 6.114 | 51.064 | 0.000 | 0.000 | 0.159 | Q9FY79 *Arabidopsis thaliana* | Laccase-14 | (Pourcel *et al.*, 2005) |

**Table 2.** Glucan endo-1,3-beta-glucosidase DEGs in the list of critical genes (C_int_) of FAZ formation. The 9 DEGs are a subset of 88 genes annotated as cell wall modification and cell development. The expression level of each group is indicated by FPKM. Magenta color denotes upregulation in MAZ and EAZ compared to GAZ, and green denotes downregulation (t-test, *p* < 0.05, n = 4). Abbreviations are listed in the legend of **Figure 4**.

| Uniport<br>Gene Name | GAZ | MAZ | EAZ | GC | MC | EC | Uniprot<br>homolog | Annotation | Ref |
| --- | --- | --- | --- | --- | --- | --- | --- | --- | --- |
| OLEA9_A032538 | 0.143 | 1.003 | 0.484 | 0.082 | 0.159 | 0.256 | P36401 *Nicotiana tabacum* | Glucan endo-1,3-beta-glucosidase, acidic isoform PR-Q' | (Bulcke *et al.*, 1989) |
| OLEA9_A051307 | 0.010 | 0.118 | 0.122 | 0.000 | 0.011 | 0.026 | Q12700 *Schwanniomyces occidentalis* | Glucan 1,3-beta-glucosidase | (Esteban, Vazquez De Aldana and Del Rey, 1999) |
| OLEA9_A089990 | 0.817 | 5.846 | 3.034 | 0.967 | 1.611 | 1.118 | Q8VYE5 *Arabidopsis thaliana* | Glucan endo-1,3-beta-glucosidase 12 | (Doxey *et al.*, 2007) |
| OLEA9_A090031 | 0.257 | 0.102 | 0.085 | 0.122 | 0.043 | 0.067 | Q9C7U5 *Arabidopsis thaliana* | Glucan endo-1,3-beta-glucosidase 2 | (Doxey *et al.*, 2007) |
| OLEA9_A012299 | 0.392 | 1.058 | 1.249 | 0.124 | 0.069 | 0.409 | Q9ZQG9 *Arabidopsis thaliana* | Glucan endo-1,3-beta-glucosidase 14 | (Wang *et al.*, 2023) |
| OLEA9_A074219 | 0.113 | 0.417 | 0.491 | 0.023 | 0.019 | 0.186 | Q9ZQG9 *Arabidopsis thaliana* | Glucan endo-1,3-beta-glucosidase 14 | (Wang *et al.*, 2023) |
| OLEA9_A110794 | 0.875 | 6.072 | 2.224 | 1.324 | 1.383 | 1.042 | Q9FJU9 *Arabidopsis thaliana* | Glucan endo-1,3-beta-glucosidase 13 | (Doxey *et al.*, 2007) |
| OLEA9_A117455 | 0.112 | 1.675 | 1.131 | 0.006 | 0.000 | 0.008 | Q8L868 *Arabidopsis thaliana* | Glucan endo-1,3-beta-glucosidase 11 | (Levy *et al.*, 2007) |
| OLEA9_A121547 | 12.490 | 40.730 | 65.947 | 2.138 | 1.431 | 2.801 | Q9ZQG9 *Arabidopsis thaliana* | Glucan endo-1,3-beta-glucosidase 14 | (Wang *et al.*, 2023) |

### 3.5 Lignification of the fruit abscission zone correlates with the reduction of the fruit removal force

The development of the FAZ is often associated with tissue lignification, as observed in a variety of fruits, including citrus, olive, and pepper, via the phloroglucinol-HCl staining (Reed and Hartmann, 1976; Merelo *et al*., 2017; Hill *et al*., 2023). Hence, we evaluated the lignification level of the olive FAZ in our samples employing Basic Fuchsin staining (**Figure 5**). Basic Fuchsin staining (Ursache *et al*., 2018) provides the advantage of quantifiable fluorescent signals and enables optical sectioning using confocal microscopy, allowing more precise localization of the lignified regions. In the green immature tissue, minimal lignin staining was detectable in the forming FAZ. However, lignification was prominent in horticulturally mature and ethephon-treated samples (**Figure 5A**). Quantification of the fluorescence signal showed a three-fold increase in horticulturally mature samples and a four-fold increase in ethephon-treated samples (one-way ANOVA, Tukey test, *p* < 0.05) (**Figure 5B**). The higher degree of lignification in both horticulturally mature and ethephon-treated samples corresponds to a similar but lower FRF of ∼200 g compared to over 300 g in green immature fruits (**Figure 3, 4B**).

**Figure 5.**
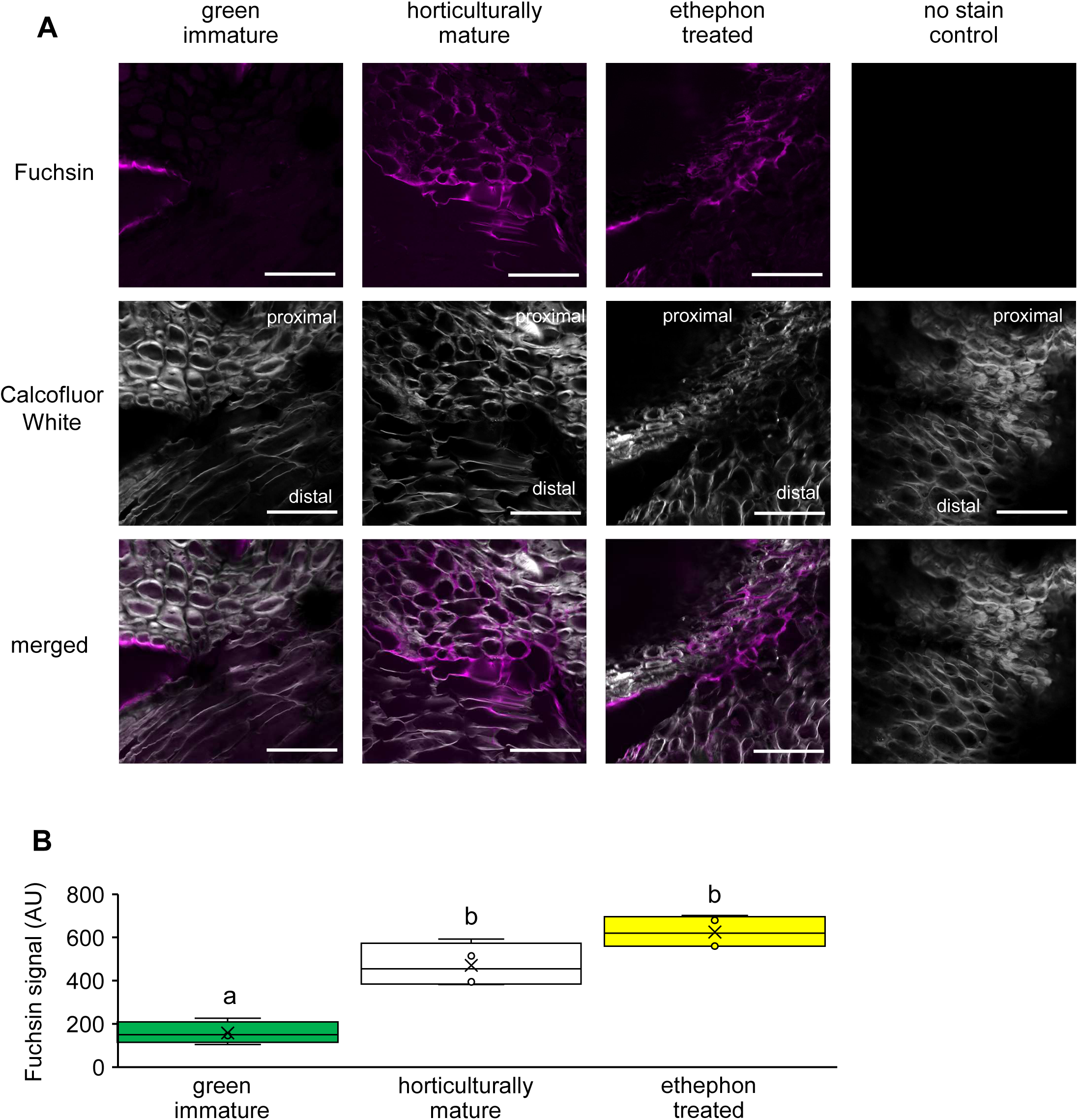
Lignification of FAZ cells stained with Basic Fuchsin. **(A)** Basic Fuchsin staining of the FAZ region as indicated in Figures 2F and **4A**. Staining of green immature, horticulturally mature, and ethephon-treated FAZ. The sections were counterstained with Calcofluor White to outline cell anatomy. No background fluorescence is detected in ethephon-treated sample without fuchsin staining. Scale bar = 100 μm. **(B)** The signal intensity in arbitrary units (AU) within the rectangle tool of 0.01 mm^2^ is measured. Different letters indicate significant difference between groups (N = 4 biological replicates for each group, one-way ANOVA, Tukey HSD, *p* < 0.05).

The increased lignification (**Figure 5**) correlates with the candidate DEGs involved in lignin biosynthesis (**Figure 4**). Specifically, the majority of the phenylpropanoid pathway-related genes were upregulated (**Table 1**). Genes of laccases and dirigent protein, typically recognized as genes directly related to lignin biosynthesis, were highly upregulated in the FAZ. Additional phenylpropanoid pathway-related genes were also upregulated, such as those implicated in the biosynthesis of lignin monomer building blocks, naringenin,2-oxoglutarate 3-dioxygenase (flavanone 3-hydroxylase, F3H) and the bifunctional pinoresinol-lariciresinol reductase (**Table 1**). In summary, both the ethylene-treated and horticulturally mature fruits were characterized by an FRF lower than 200 g, which correlated with upregulation of lignin biosynthesis genes and higher tissue lignification at the FAZ, demonstrating the similarity between the natural maturation and ethylene treatment response.

### 3.6 Ethephon treatment induces glucanase activity and reduces plasmodesmata callose at the fruit abscission zone

Beyond lignin deposition, the FAZ undergoes other significant cell wall changes. Among the 733 DEGs (**C_int_**), 88 candidate genes were identified as being involved in cell wall modification and cell development (**Figure 4D**), including nine annotated as β-1,3-glucanases (**Table 2**). Eight of the nine β- 1,3-glucanase genes were upregulated, suggesting a high callose degradation at the FAZ. These genes are homologs of Arabidopsis BG11, 12, 13, and 14 (**Table 2**), which all belong to clade α in the β-1,3- glucanase gene family (Doxey et al., 2007). OLEA9_A121547, which shows the highest expression level among the nine DEGs, is a homolog of the Arabidopsis BG14 gene. Disruption of BG14 leads to enhanced callose deposition and reduced plasmodesmata permeability in developing Arabidopsis seeds (Wang *et al*., 2023). Ethylene treatment or maturation leading to upregulation of β-1,3-glucanase transcripts has been found in many plant tissues, such as mature olive FAZ (Gil-Amado and Gomez-Jimenez, 2013), rice shoots (Simmons et al., 1992), ripening banana mesocarp (Choudhury *et al*., 2010), and dormancy-breaking grape buds (Shi *et al*., 2025). Here, we demonstrate that the olive FAZ is exhibiting the same upregulation pattern of β-1,3-glucanase genes during maturation and after ethephon treatment.

In order to better understand callose remodeling at the FAZ, we performed immunohistology studies, using the BS-400-2 β-1,3-glucan-specific antibody (Meikle *et al*., 1991; Thorne, Urbanowicz and Hahn, 2023). **Figure 6A** shows abundant plasmodesmata callose in each cell at the immature stage, but markedly reduced levels in horticulturally mature and ethephon-treated samples. Quantification of the number of fluorescent puncta revealed an over 90% decrease after ethephon treatment (**Figure 6B**). The control image with no primary antibody using a green immature sample shows no background fluorescence (**Figure 6C**). Live staining of callose at the FAZ by the aniline blue fluorochrome further corroborated the diminishing levels of callose (**Supplemental Figure 2**). Plasmodesmata permeability is regulated by callose, whose turnover is controlled by β-1,3-glucanases as has been shown in Arabidopsis and tobacco (Beffa *et al*., 1996; Levy *et al*., 2007; Zavaliev *et al*., 2011). Thus, we investigated the glucanase activity in the FAZ tissue to validate whether it correlates with the callose reduction. Our results showed an increased glucanase activity 2 days after ethephon treatment, however, not after 7 days (**Figure 6D**). The observed induction in glucanase activity upon ethephon treatment is in line with earlier studies in tobacco leaves (Felix and Meins Jr, 1987). On the basis of our expression, anatomical, and activity data, we conclude that ethylene-induced upregulation of β-1,3-glucanase genes results in higher activity, which in turn leads to reduced callose deposition at plasmodesmata, thereby promoting symplasmic flow during olive FAZ development.

**Figure 6.**
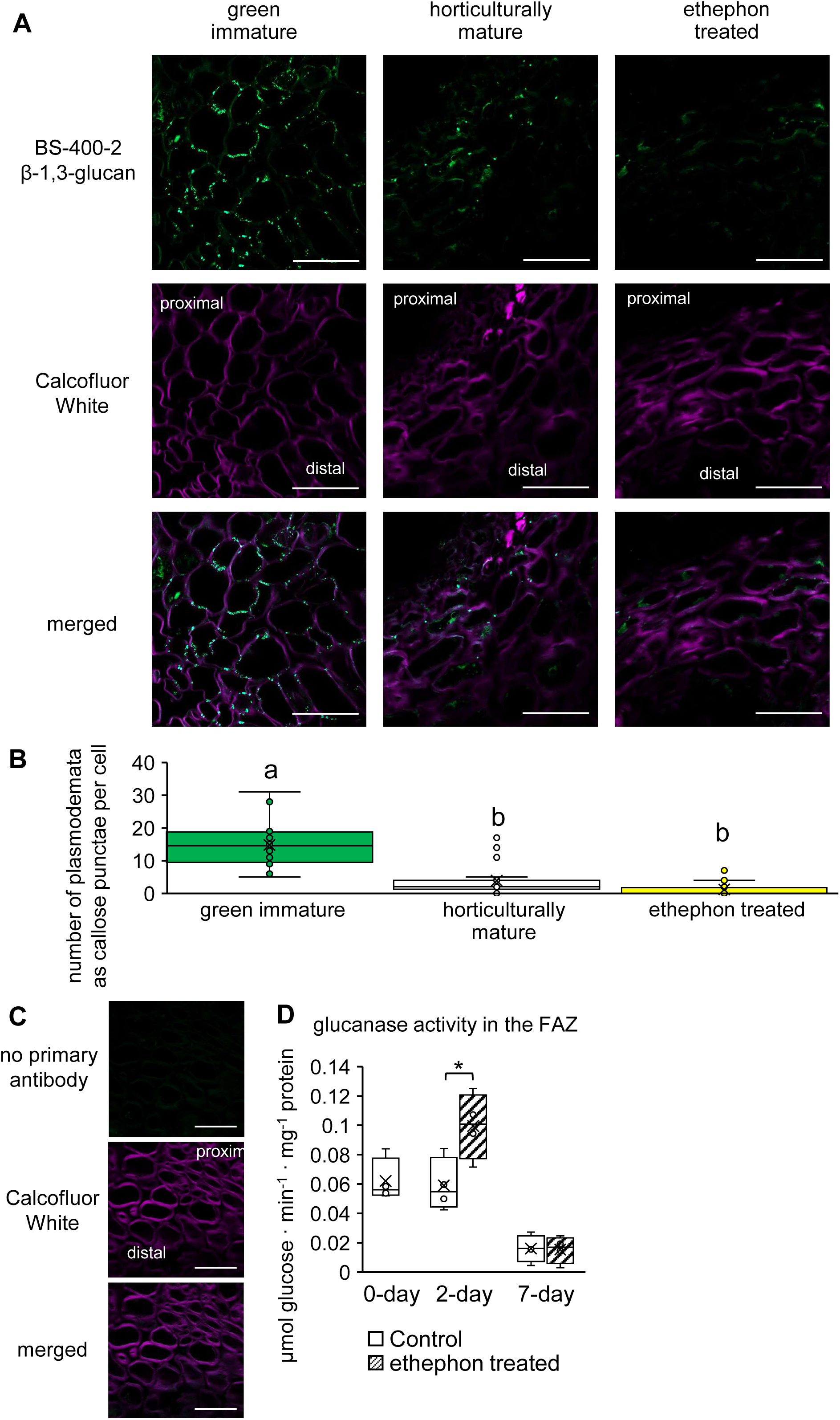
Plasmodesmata callose and glucanase activity at the FAZ. (**A**) β-1,3-glucan detection using the BS-400-2 antibody shows plasmodesmata callose at the FAZ in the three treatment groups. Note the higher immunofluorescence detection in green immature compared to horticulturally mature and ethephon-treated FAZ. Calcofluor White shows the cell profile. The Calcofluor White channel was set to display a signal intensity range of 0-7000 arbitrary units in ZEN Lite in all three groups, for contrasting the callose signal display. Scale bar = 50 μm. (**B**) Quantification of the number of plasmodesmata via callose detection. Five consecutive cells were counted in each image for the number of plasmodesmata, and the average number of plasmodesmata on each cell was calculated. Different letters indicate a significant difference between groups (N = 4 biological replicates and n = 20 cells for each group, one-way ANOVA, Tukey HSD, *p* < 0.05). (**C**) Control images with no primary antibody using a green immature sample. (**D**) Glucanase activity assay. Glucanase activity in the FAZ from total crude extract after ethephon treatment compared to the control is shown. FAZ samples were collected prior to ethephon application, 0-day, 2 days after ethephon application and 7 days after ethephon application. Asterisk indicates a significant difference (n = 4, t-test, *p* < 0.05).

### 3.7 Alkalization of fruit abscission zone prior to cell separation

Alkalization of the abscission zone was first reported in Arabidopsis flowers (Sundaresan *et al*., 2015), and has been proposed as a general mechanism for organ separation, including fruit abscission (Patharkar and Walker, 2018; Pautot, Crick and Hepworth, 2025). We used a previously established method employing the BCECF staining/chlorophyll ratio (Sundaresan *et al*., 2015; Ying *et al*., 2016; Ma *et al*., 2020) to monitor the pH of the olive FAZ. We observed an enhanced BCECF signal at the FAZ of horticulturally mature and ethephon-treated fruits, compared to the green immature ones (**Figure 7A**). Quantification of fluorescence signal showed a higher BCECF/chlorophyll ratio in the ethephon-treated samples compared to the immature samples (**Figure 7A, B**) (one-way ANOVA, *p* < 0.05, n = 4 for each group), suggesting that FAZ alkalization is induced by ethephon treatment. When comparing the BCECF signal in horticulturally mature and immature samples by t-test, the mature samples showed a significantly higher signal (t-test, *p* < 0.005, n = 4). A cytosolic pH increase in the ethephon-treated sample was indicated when we analyzed the samples under a 10X objective (**Figure 7A**, arrow). Across the samples, no chlorophyll autofluorescence level difference was detected at the FAZ region (**Figure 7C**). Hence, the observed BCECF/chlorophyll ratio change (**Figure 7B**) is dependent on the level of BCECF signal. In order to better resolve the subcellular localization of the BCECF signal, the immature and ethephon-treated samples were examined under higher magnification. Compared to the immature sample, the BCECF signal was found strongly accumulated in the cytosol in the ethephon-treated sample (**Figure 7D**, **E**, magenta arrow), and also at the cell wall and at the tricellular junction (TCJ) (**Figure 7D**, **F**, orange and white arrow), suggesting that both cytosolic and apoplastic pH increase in ethephon-treated samples. In summary, here we show the first experimental evidence of FAZ alkalization in plants.

**Figure 7.**
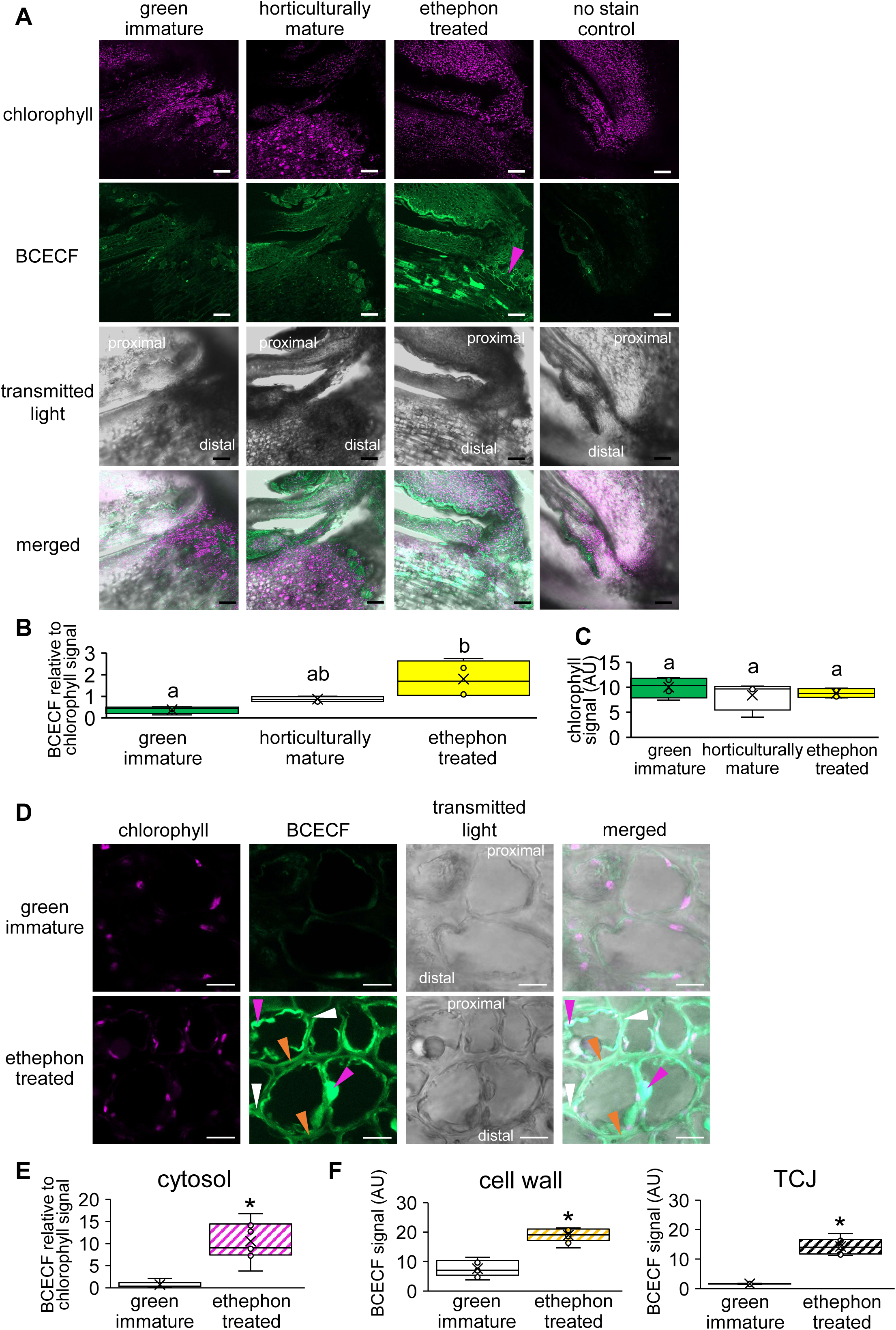
pH staining of olive FAZ using BCECF. (**A**) pH imaging of FAZ sections within the green mature, horticulturally mature, and ethephon-treated samples. BCECF, chlorophyll, and transmitted light channels are shown imaged with a 10X objective. Chlorophyll autofluorescence shows a constant emission signal across all groups. Ethephon-treated and horticulturally mature samples show enhanced signal compared to the control. The magenta arrow points to the high BCECF signal in the cytosol of FAZ cells. No stain control is imaged using an ethephon-treated sample. Each image is a maximum projection of a 12-slice z-stack. Scale bar = 100 μm. (**B**) Signal intensity of BCECF at the FAZ sections of the three groups is shown. The signal intensity was measured within the rectangle tool of 0.01 mm^2^ for BCECF and the chlorophyll channel, and their ratio was used as the proxy of pH changes. Different letters indicate significant difference between groups (N = 4 biological replicates for each group, one-way ANOVA, Tukey HSD, p < 0.05). Note the increased signal intensity of ethephon-treated and horticulturally mature samples compared to the control. (**C**) Chlorophyll signal intensity remained the same across all groups. Different letters indicate significant difference between groups (N = 4 biological replicates for each group, one-way ANOVA, Tukey HSD, *p* < 0.05). (**D**) Cytosol and apoplast alkalization at FAZ. BCECF, chlorophyll, and transmitted light images of green immature and ethephon-treated FAZ sections by a 40X objective are shown. Magenta arrows indicate a high BCECF signal at the cytosol, orange arrows point to the BCECF signal at the cell wall, and white arrows point to the BCECF signal at TCJ. Images represent single confocal planes. Scale bar = 10 μm. (**E**) Quantification BCECF/chlorophyll ratio of cytosol in immature and ethephon-treated FAZ samples. A strong BCECF signal is observed in the cytosol of FAZ cells in ethephon-treated samples. Signal intensity of BCECF and chlorophyll channels within the circle tool of 0.30 μm^2^ in cytosol was measured, and three spots of each biological replicate were measured. Asterisk indicates a significant difference in BCECF/chlorophyll ratio (N = 3 biological replicates and n = 9 spots in the cytosol for each group, t-test, *p* < 0.05). (**F**) BCECF signal at the cell wall or the TCJ in immature and ethephon-treated FAZ samples. No chlorophyll accumulates at the apoplast; hence, the chlorophyll channel was not used as a reference for the apoplast BCECF signal. BCECF levels are higher in the cell wall and TCJ in ethephon-treated compared to the green immature samples. Signal intensity within the circle tool of 0.30 μm^2^ on the cell wall was measured, and three spots of each biological replicate were measured. Asterisk indicates a significant difference (N = 3 biological replicates and n = 9 spots in the cell wall for each group, t-test, *p* < 0.05).

### 3.8 Low-methylesterified homogalacturonan decreases as the FAZ develops

In order to study the pectin changes in the FAZ during development and after ethephon treatment, we used JIM5/JIM7 antibodies targeting low/high-methylesterified homogalacturonan, respectively (Knox *et al*., 1990). The JIM5 signal was high in the TCJ of immature samples, and it decreased similarly after maturation or ethephon treatment (**Figure 8A**), while the JIM7 signal did not show a difference across the groups (**Figure 8B**). There was no background fluorescence in the absence of antibody labelling (**Figure 8C**). This suggests that the low-methylesterified homogalacturonan levels are reduced at the TCJ, while the high-methylesterified homogalacturonan levels do not change during maturation or after ethephon treatment.

**Figure 8.**
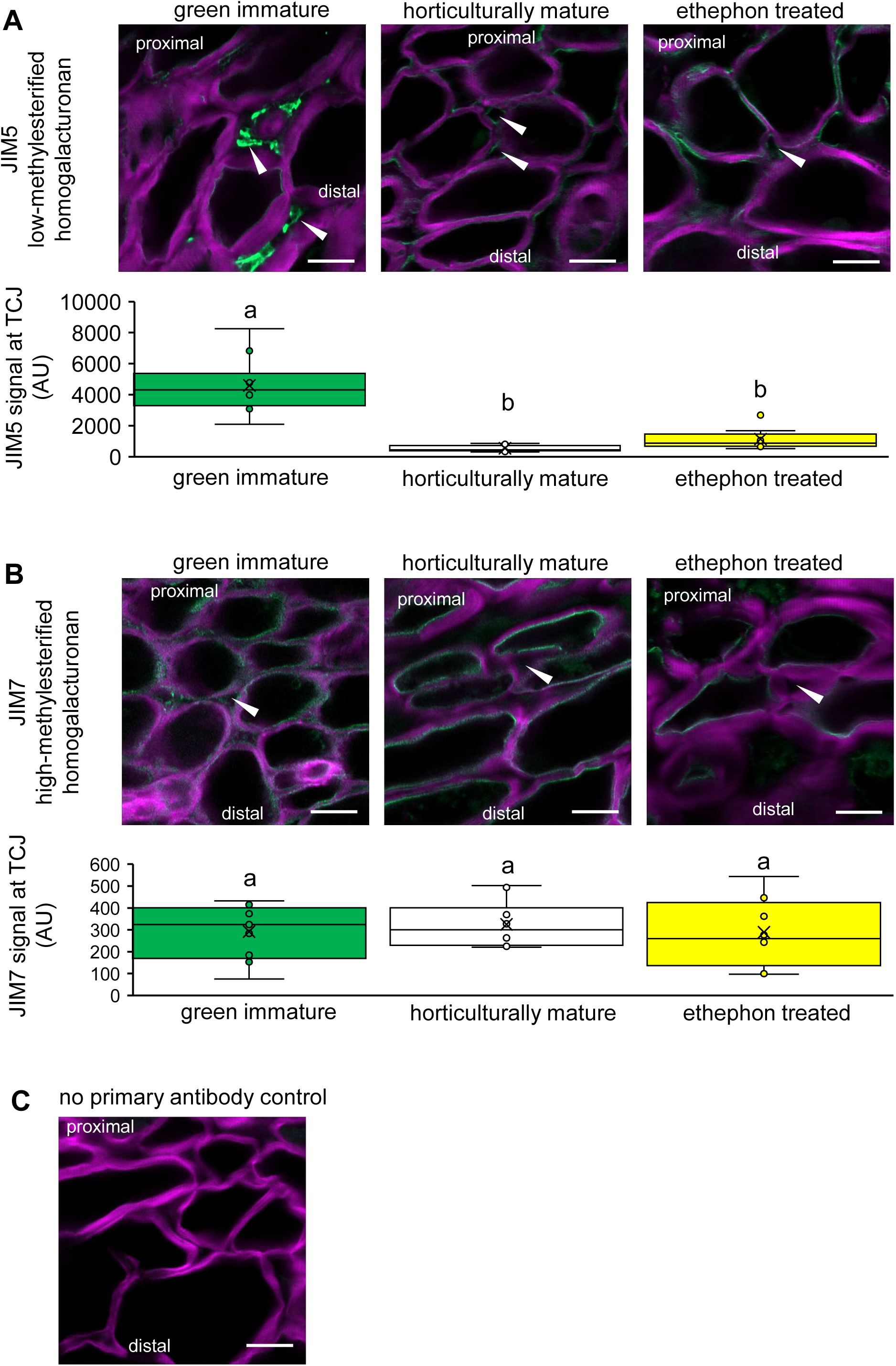
Immunolocalization of pectin modification at the FAZ. (**A**) Low-methylesterified homogalacturonan targeted by JIM5. Arrows point to TCJ. Low-methylesterified homogalacturonan accumulated at TCJ in the immature sample. Images are shown as merged format with Calcofluor white staining showing cell profiles as the counterstain. Signal intensity applying the Spline Contour tool, selecting TCJ, is measured in ZEN Lite. Different letters indicate a significant difference between groups (N = 4 biological replicates and n = 12 TCJs for each group, one-way ANOVA, Tukey HSD, *p* < 0.05). Scale bar = 10 μm. (**B**) High-methylesterified homogalacturonan targeted by JIM7. Arrows point to TCJ. No high-methylesterified homogalacturonan level change was detected in horticulturally mature or ethephon-treated samples (**C**) Control images with no primary antibody using a green immature sample.

Among our list of 733 candidate genes for FAZ development (**Figure 4**) upregulation of genes involved in pectin modification, such as PL, protein trichome birefringence-like (TBL) 39 and 40, and pectin methylesterase (PME) inhibitor, was prominent (**Table 3**). Given that PL acts on the low-esterified form of pectin (Seyedarabi *et al*., 2010), the upregulation of *PL* can account for the reduced low-methylesterified homogalacturonan at the TCJ as FAZ develops. TBL39 and TBL40 are implicated in maintaining esterification of pectin (Bischoff *et al*., 2010). Together with the PME inhibitor, TBL39/40 may be upregulated to maintain the methylesterification level of pectin, consistent with our observation that the levels of high-methylesterified homogalacturonan remain unchanged during maturation and after ethephon treatment.

**Table 3.** Pectin modification enzyme-related DEGs in the list of critical genes (C_int_) of FAZ formation. The 7 DEGs are a subset of 88 genes annotated as cell wall modification and cell development. The expression level of each group is indicated by FPKM. Magenta color denotes upregulation in MAZ and EAZ compared to GAZ, and green denotes downregulation (t-test, *p* < 0.05, n = 4). Abbreviations are listed in the legend of **Figure 4**.

| Uniport<br>Gene Name | GAZ | MAZ | EAZ | GC | MC | EC | Uniprot<br>homolog | Annotation | Ref |
| --- | --- | --- | --- | --- | --- | --- | --- | --- | --- |
| OLEA9_A020370 | 0.037 | 0.257 | 1.604 | 0.051 | 0.000 | 0.046 | O24554 *Zinnia violacea* | Pectate lyase | (Domingo *et al.*, 1998) |
| OLEA9_A011623 | 0.022 | 0.000 | 0.000 | 0.020 | 0.002 | 0.003 | P05117 *Solanum lycopersicum* | Polygalacturonase-2 | (Chun and Huber, 1998) |
| OLEA9_A036088 | 0.151 | 1.668 | 1.094 | 0.036 | 0.023 | 0.102 | A7PW81 *Vitis vinifera* | Polygalacturonase inhibitor | (De Ascensao, 2001) |
| OLEA9_A121155 | 0.414 | 5.408 | 1.230 | 0.000 | 0.000 | 0.000 | P17407 *Daucus carota* | PF04043:Plant invertase/pectin methylesterase inhibitor | (Coculo and Lionetti, 2022) |
| OLEA9_NCA022907 | 0.222 | 0.577 | 1.512 | 0.037 | 0.006 | 0.044 | Q9SIN2 *Arabidopsis thaliana* | Protein trichome birefringence-like 39 | (Bischoff *et al.*, 2010) |
| OLEA9_A034821 | 0.000 | 0.139 | 0.255 | 0.000 | 0.010 | 0.010 | Q67XC4 *Arabidopsis thaliana* | Protein trichome birefringence-like 40 | (Bischoff *et al.*, 2010) |
| OLEA9_A096439 | 0.516 | 0.028 | 0.083 | 0.039 | 0.000 | 0.000 | O48711 *Arabidopsis thaliana* | Probable pectinesterase/pectinesterase inhibitor 12 | (Coculo and Lionetti, 2022) |

### 3.9 Non-fucosylated xyloglucan is reduced at the FAZ

Previous studies suggest that hemicellulose modifications contribute to cell wall remodeling during FAZ development in olive, as indicated by a stronger xylan signal (detected with LM11) in pre-abscission samples and a weaker xyloglucan signal (detected with LM15) in post-abscission samples (Parra and Gomez-Jimenez, 2020). Therefore, beyond pectin and lignin, we examined potential changes in the state of xyloglucan. Fucosylated and non-fucosylated xyloglucan are commonly detected and distinguished by M1 and M101 of the CCRC series of monoclonal antibodies (Pattathil *et al*., 2010), while the levels of these epitopes are known to be altered in xyloglucan biosynthesis mutants (Zabotina *et al*., 2012; Zhang *et al*., 2023). Fucosylated xyloglucan levels showed no significant variations between the three groups (**Figure 9A**). This was contrasted by a decrease of non-fucosylated xyloglucan at the FAZ with fruit maturation or ethephon treatment (**Figure 9B**). No background fluorescence was detected in the absence of antibody labelling (**Figure 9C**). The M101 epitope has been detected in the SYNTAXIN OF PLANTS61 *trans*-Golgi network compartment (Wilkop *et al*., 2019), leading to the hypothesis that non-fucosylated xyloglucan can be deposited as a structural polysaccharide in the cell walls. The observed reduction in non-fucosylated xyloglucan may therefore reflect an increased degradation of this structural hemicellulose epitope during FAZ development.

**Figure 9.**
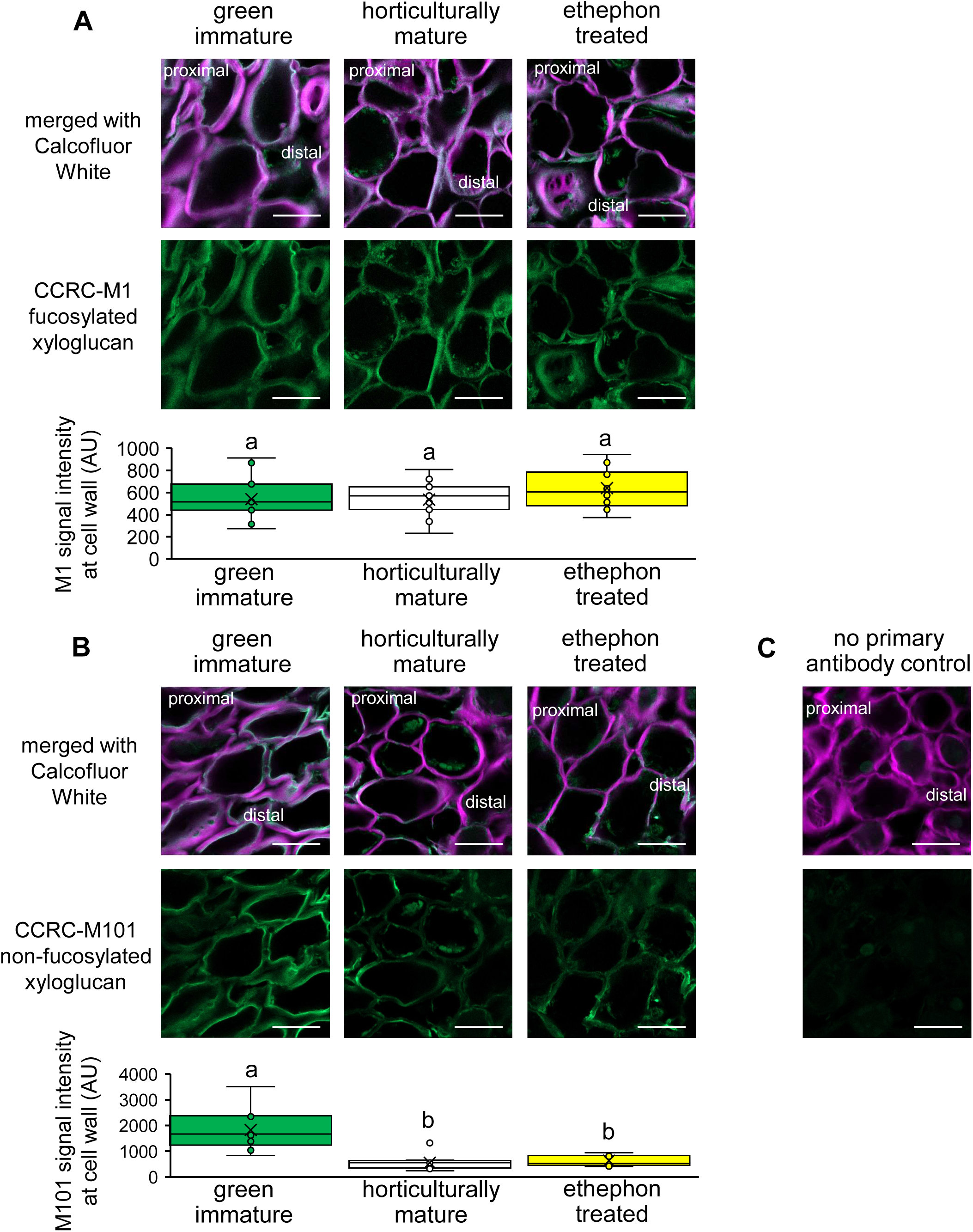
Hemicellulose modification at the FAZ revealed by immunolocalization. (**A**) Fucosylated xyloglucan targeted by CCRC-M1. No fucosylated xyloglucan change at the cell wall was detected with maturation or after ethephon treatment. Signal intensity within the circle tool of 0.30 μm^2^ on the cell wall was measured, and three spots of each biological replicate were measured. Different letters indicate a significant difference between groups (N = 4 biological replicates, n = 12 spots on the cell wall for each group, one-way ANOVA, Tukey HSD, *p* < 0.05). Scale bar = 20 μm. (**B**) Non-fucosylated xyloglucan targeted by CCRC-M101. Lower levels of non-fucosylated xyloglucan were detected at the cell walls of horticulturally mature and ethephon-treated samples compared to immature samples. (**C**) Control images with no primary antibody using a green immature sample.

Five genes encoding hemicellulose-modifying enzymes, including xylosidase, xylan glycosyltransferase, and xyloglucan endotransglucosylase, were identified among the candidates for FAZ development in the cell wall modification category (**Figure 4D**, **Table 4**), implying a complex hemicellulose remodeling during FAZ development and after ethephon treatment. The genes of two xyloglucan endotransglucosylases, *XTH2* and *XTH24*, both decreased after maturation or ethephon treatment (**Table 4**). In a previous comparison of the transcriptome of mature and ripe olive FAZ (154 vs 217 dpa), several *XTHs* were reported up- or downregulated (Gil-Amado & Gomez-Jimenez, 2013), but neither *XTH2* or *XTH24* was identified as DEGs. While our results show an increase in lignin and a reduction of low-methylesterified homogalacturonan, the detailed molecular and cellular mechanism of hemicellulose modification at the FAZ remains to be elucidated.

**Table 4.** Hemicellulose modification enzyme-related DEGs in the list of abscission zone critical players (C_int_). The 5 DEGs are a subset of 88 genes annotated as cell wall modification and cell development. The expression level of each group is indicated by FPKM. Magenta color denotes upregulation in MAZ and EAZ compared to GAZ, and green denotes downregulation (t-test, *p* < 0.05, n = 4). Abbreviations are listed in the legend of **Figure 4**.

| Uniprot<br>Gene Name | GAZ | MAZ | EAZ | GC | MC | EC | Uniprot<br>homolog | Annotation | Ref |
| --- | --- | --- | --- | --- | --- | --- | --- | --- | --- |
| OLEA9_A091862 | 0.129 | 0.004 | 0.000 | 0.022 | 0.000 | 0.000 | Q9SSE8 *Arabidopsis thaliana* | Probable glycosyltransferase At3g07620 | (Li *et al.*, 2004) |
| OLEA9_A101334 | 0.137 | 0.049 | 0.035 | 0.027 | 0.000 | 0.000 | Q9LJN4 *Arabidopsis thaliana* | Probable beta-D-xylosidase 5 | (Iglesias *et al.*, 2006) |
| OLEA9_A023165 | 0.306 | 0.726 | 1.397 | 0.134 | 0.107 | 0.214 | Q9SS43 *Arabidopsis thaliana* | Xylan glycosyltransferase MUCI21 | (Voiniciuc *et al.*, 2015) |
| OLEA9_A043949 | 0.414 | 0.129 | 0.051 | 0.092 | 0.110 | 0.094 | P35694 *Glycine max* | Xyloglucan endotransglucosylase/hydrolase 2 | (Niraula *et al.*, 2021) |
| OLEA9_A112468 | 1.206 | 0.217 | 0.247 | 0.065 | 0.026 | 0.077 | P24806 *Arabidopsis thaliana* | Xyloglucan endotransglucosylase/hydrolase protein 24 | (Rose *et al.*, 2002) |

## 4. Discussion

### 4.1 Candidate transporters for olive FAZ alkalization

This work presents a comprehensive study of the fruit abscission zone (FAZ) in a woody perennial species, analyzing in parallel two types of responses: (1) those occurring during natural FAZ maturation and (2) those induced by ethylene treatment. By comparing these two conditions, the study provides an integrated framework to identify developmental events intrinsic to natural abscission from those accelerated or enhanced by ethylene. This parallel analysis of natural and ethylene-induced responses offers new insights into the molecular and structural mechanisms underlying FAZ development in woody perennials.

Further, it provides the first evidence of FAZ alkalization prior to abscission in plants. The abscission zone alkalization in flowers is tightly associated with the ethylene signaling pathway, as shown in the Arabidopsis *ethylene-insensitive 2* and *ethylene overproducer 4* mutants, in which alkalization is delayed or absent (Sundaresan *et al*., 2015). It has been suggested that upregulation of certain transporters, such as a vacuolar H^+^-ATPase and a putative high-affinity nitrate transporter, may be associated with the cytosol alkalization of the flower abscission zone (Sundaresan *et al*., 2015). Given that membrane transporters regulate the pH of subcellular compartments (Pittman, 2012), we examined FAZ-specific transporters from our transcriptome analysis. The most prominently upregulated transporter is OLEA9_A070043, a homolog of the Arabidopsis lysine histidine transporter 1 (LHT1) (**Supplemental Table 5**). Consistent with our findings is the observation that in the flower abscission zone, the *LHT1* expression is enhanced in later stages of Arabidopsis flower development (Niederhuth, Patharkar and Walker, 2013). This could suggest a conserved function of the transporter across species. LHT1 was first identified based on its high transport rates for lysine and histidine, two alkaline amino acids, in a yeast complementation assay (Chen and Bush, 1997). An enrichment of lysine and histidine in the apoplastic wash fluids of *lht1* knockout leaves in Arabidopsis (Hirner *et al*., 2006) suggests that these amino acids serve as *in vivo* substrates of LHT1. Beyond these two amino acids, the ethylene precursor ACC has been suggested as an *in vivo* substrate of LHT1, demonstrated by the ACC-insensitive phenotype of *lht1* mutants (Shin *et al*., 2015).

The affinity of LHT1 with the substrate ACC indicates a potential intersection with ethylene signaling. Ethylene responses have been linked to amino acid metabolism, particularly histidine and lysine, in Arabidopsis and tomato. For example, application of 5 mM L-histidine induces bacterial resistance through activation of ethylene signaling (Seo *et al*., 2016). Furthermore, upregulation of lysine metabolism has been implicated in abscission zone development, as shown in the cotton boll abscission zone, in which elevated expression of the lysine-ketoglutarate reductase gene following ethephon treatment was revealed by *in situ* hybridization (Tang *et al*., 2002; Stepansky *et al*., 2006). Although amino acid uptake assays have not yet been performed in the abscission zone tissue of *lht* mutants, the above studies suggest that ACC, lysine, and histidine may be transported into the abscission zone via LHT. Given that the LHT homolog is the most prominently upregulated FAZ-specific transporter in our analysis, it represents a strong candidate for mediating lysine and histidine influx from proximal tissues (pedicel) into the abscission zone during FAZ development, potentially contributing to cytosolic alkalization. In order to validate this hypothesis, amino acid levels (including ACC) could be compared between the abscission zone and the surrounding tissues. In species with large FAZs (e.g., citrus), free-hand sectioning could be used, and laser microdissection could be used for smaller ones like the *lht1* and wild-type Arabidopsis flower abscission zones, which are distinctively composed of small packed cells (Pautot, Crick and Hepworth, 2025). In addition to cytosol alkalization, we also observed an increase in apoplastic pH at the olive FAZ following ethephon treatment, compared to the immature samples (**Figure 7D, F**). A general increase in the apoplastic pH is known to occur under stress conditions or pathogen attack. (Geilfus, 2017). In this study, we identified 137 FAZ-specific DEGs annotated as components of stress and plant immunity response pathways. It is therefore plausible that the FAZ apoplast exhibits features similar to those observed under general stress conditions, which may account for the observed pH increase (**Figure 7D, F**, **Supplemental Table 6**).

### 4.2 A trend of pectin remodeling corroborates apoplast alkalization during FAZ development

Our study shows pectin remodeling with a specific decrease in the low methyl esterified form at the FAZ, in both natural maturation and ethephon-treated samples. In earlier studies, the use of LM19 and LM20 antibodies, which recognize low- and high-methylesterified pectins, respectively, functionally analogous to JIM5 and JIM7, revealed that pre-abscission olive FAZ tissues exhibited a stronger LM19 signal than LM20 (Parra *et al*., 2020). PL activity correlates with pectin modification, as shown in tomato, in which *PL* silencing leads to higher levels of low-methylesterified homogalacturonan at the TCJ of ripe pericarp cells (Uluisik *et al*., 2016). Consistent with the observed upregulation of a FAZ-specific *PL* in our DEGs of **C_int_**, we report a reduced level of low-methylesterified homogalacturonan at the TCJ as the olive FAZ develops (**Table 3**, **Figure 8**). Thus, we consider that PL activity is a major contributor to the pectin remodeling at the FAZ.

Our transcriptome analysis revealed seven FAZ-specific genes involved in pectin modification, including genes annotated as PL, PG, PG inhibitors, PME inhibitors, and TBLs (**Table 3**). Most of these DEGs favor an alkaline pH for pectin remodeling activity (Fry, 2017). Aligned with the expectation of this pH trend is the observed pH increase at the apoplast after FAZ development, identified through our histochemical analysis via BCECF staining (**Figure 7D, F**). The pectin-modifying enzymes PL and PG both cleave pectin into smaller subunits, but with different optimal pH conditions: PL catalyzes the β- elimination reaction at an optimal pH around 8, and endo-PG catalyzes the endo-hydrolysis of pectin at an optimal pH around 5 (Niture *et al*., 2008; Fry, 2017; Uluisik and Seymour, 2020). The upregulation of *PL*, concurrent downregulation of *PG*, and increased expression of *PG INHIBITOR* (**Table 3**) are consistent with the observed apoplast alkalization and reduced levels of low-methylesterified homogalacturonan at the apoplast (**Figure 7D, F, 8A**). PME catalyzes the removal of methyl groups and exposes galacturonic acid residues (Fry, 2017), while PME inhibitors restrict this process. Presumably, the upregulation of PME inhibitors restricts de-methylesterification, and the upregulation of TBLs maintains methylesterification (**Table 3**), resulting in reduced exposure of the galacturonic acid ends, also consistent with the observed apoplast alkalization and constant levels of high-methylesterified homogalacturonan (**Figure 7D, F, 8B**). Both cellular and molecular evidence support that pectin remodeling at the apoplast follows a trend towards alkaline conditions during FAZ development. We propose that the activity of transporters such as LHT contributes to this high pH environment for pectin remodeling at FAZ (**Figure 10**).

**Figure 10.**
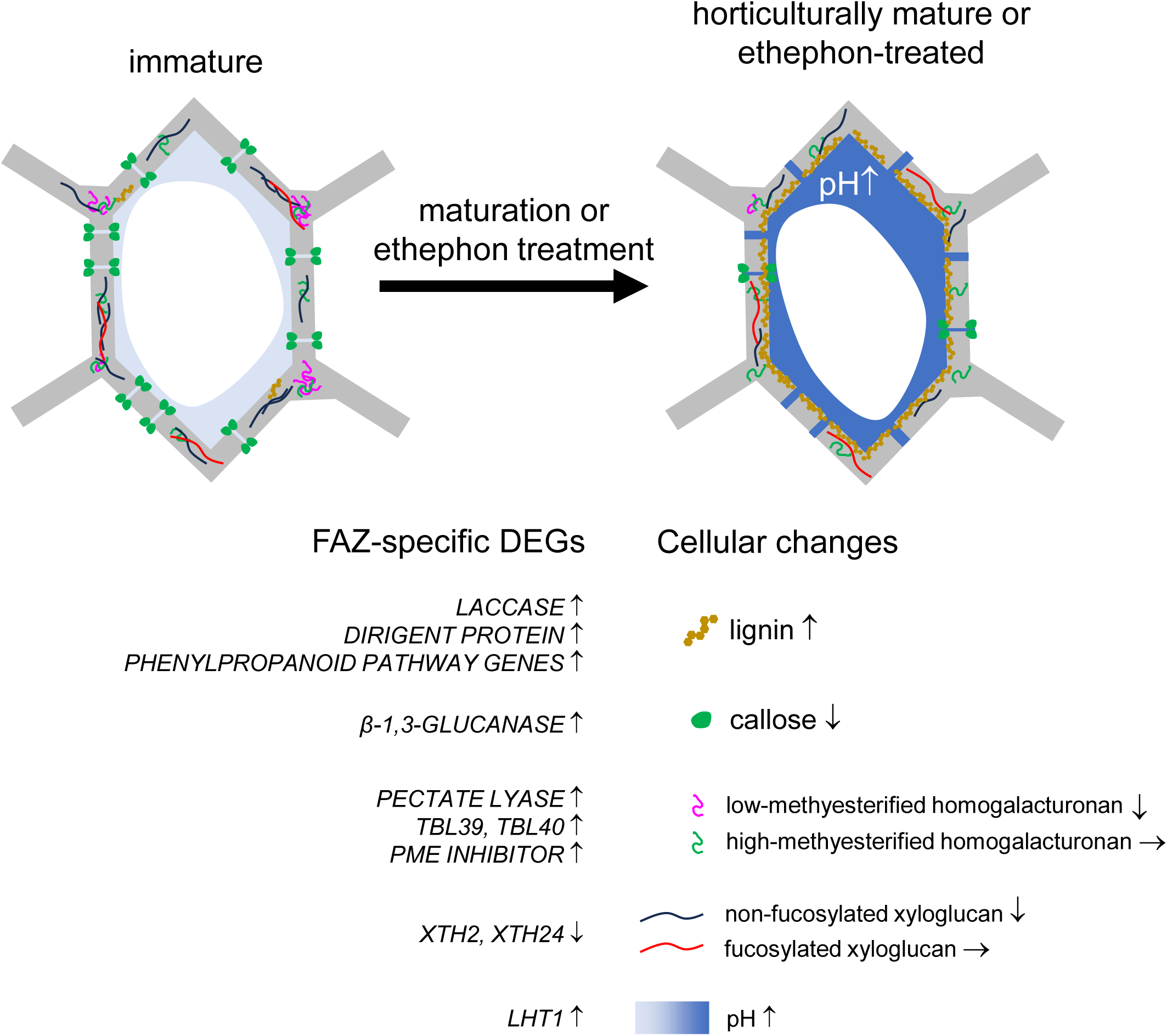
Model of olive FAZ development. With the exception of *XTH2/XTH24* and fucosylated/non-fucosylated xyloglucan, the molecular and cellular events illustrated represent inferred cause-and-effect relationships between gene expression and cellular changes during FAZ development. Upregulation of laccase, dirigent protein, and phenylpropanoid pathway genes leads to lignification of cell walls at the FAZ. Increased expression of β-1,3-glucanases results in reduced callose deposition at plasmodesmata, promoting symplasmic flow. Upregulation of *PL*, *TBL39/40*, and *PME INHIBITOR* induces a decrease in low-methylesterified homogalacturonan and maintenance of high-methylesterified homogalacturonan. Both apoplastic and cytosolic pH increase during FAZ development.

### 4.3 Identification of FAZ-specific genes

This study presents a specific set of genes (**C_int_**) critical for FAZ development, providing a foundation for both basic and applied research. The methodology ensured specificity by minimizing contamination from the mesocarp. The olive FAZ consists of only 4-5 cell layers with a width of less than 200 μm, as indicated by lignin staining (**Figure 5**). Without the assistance of histochemical staining, technologies such as laser microdissection are unable to reliably distinguish the FAZ from the surrounding tissues. However, histochemical staining is not generally considered compatible with the RNA quality requirement of transcriptome analysis (Ian, 2005; Shi and Bressan, 2006). Using a 1 mm³ tissue sample, the minimum size possible with free-hand sectioning, as the designated abscission zone for transcriptome analysis would likely cause DGE results to predominantly capture mesocarp cell wall remodeling, rather than FAZ-specific events. To address this, we employed a subset subtraction strategy, removing the mesocarp-responsive genes to identify shared regulators of FAZ development between natural maturation and ethephon treatment, thereby preventing the candidate list from being dominated by genes associated with mesocarp remodeling. **Supplemental Figure 3** illustrates the effectiveness of this approach, using *PL* and *ACC OXIDASE* as examples. In the DGE analysis of MAZvsGAZ, eight PL and nine ACC oxidase genes were identified. Many of them displayed expression patterns in naturally mature (MAZvsGAZ) and ethylene-induced (EAZvsGAZ) fruit abscission zone that are similar to mesocarp responses (MCvsGC, ECvsGC), suggesting that they are more likely to be the genes involved in the mesocarp cell wall remodeling. After applying the subtraction strategy (**C_int_**), one FAZ-specific *PL* (OLEA9_A020370, **Table 3**) and one FAZ-specific *ACC OXIDASE* (OLEA9_A097915, **Supplemental Table 2**) were identified. OLEA9_A020370, *PL*, showed distinctive upregulation in MAZ and EAZ (7- and 43-fold higher than GAZ, respectively), but did not significantly change in MC and EC, indicating its FAZ-specific role. Similarly, OLEA9_A097915, *ACC OXIDASE*, is 6-fold and 5-fold upregulated in MAZ and EAZ, but not significantly changed in MC and EC. Rather than targeting the entire PL and ACC oxidase gene families, future bioengineering and genetic screening efforts could focus on the FAZ- specific homologs. These FAZ-specific homologs could be engineered into ethylene-sensitive variants in plants, or their expression levels could be used as markers to screen for cultivars with early FAZ development. Such approaches would support the generation of ethylene-sensitive cultivars with a high fruit abscission rate at maturation.

### 4.4 Olive FAZ development model

Overall, this study offers an integrative perspective on FAZ development in a woody perennial by comparing the molecular and cellular events that occur naturally during maturation with those elicited by ethylene treatment. This framework highlights the convergence of these pathways, revealing how ethylene accentuates or reprograms processes inherent to natural abscission. By linking *in muro* cell wall modifications with transcriptional and anatomical changes, this work deepens our understanding of the mechanisms governing FAZ differentiation.

Key molecular and cellular events of olive FAZ development were identified (**Figure 10**), corresponding with the developmental reduction of FRF. We observed a reduction in plasmodesmatal callose, alkalization of the FAZ pH, decreased levels of low-methylesterified homogalacturonan at the TCJ, and reduced non-fucosylated xyloglucan in the cell wall, along with increased lignification.

FAZ alkalization is consistent with pectin modifications. Correspondingly, we identified FAZ-specific upregulated genes for these events, including phenylpropanoid pathway-related genes, PL genes, and β- 1,3-glucanase family genes. Ethylene-induced upregulation of β-1,3-glucanase genes in the olive FAZ reduced plasmodesmata callose, enhancing symplasmic connectivity and facilitating abscission. This highlights callose remodeling as a central mechanism coordinating cellular communication during FAZ development.

Core cellular and molecular features identified through this work can be used as markers in future applications, such as evaluating fruit loosening agents to reduce FRF and genetic screening for high- abscission-at-maturity cultivars to facilitate mechanical harvesting. Given the limited number of cell biology studies on plant FAZs, this work also provides a foundational reference for delineating conserved features of FAZ development.

## Acknowledgment

The authors thank Taylor Synstelien for help in field sample collection. This work is supported by the California Olive Committee and the United States Department of Agriculture Hatch (CA-D-PLS-2132-H) to GD. SZ was partially supported by the Department of Plant Sciences Graduate Student Research fellowship funded by the McDonald Endowments and facilitated by Agriculture and Natural Resources from University of California, Davis. SZ was partially supported by the Henry Jastro Graduate Research Award, the Katherine Esau Summer Graduate Fellowship, and the Agriculture and Food Research Initiative Competitive Grants Program, pre-doctoral fellowship project award no. 2023-67011-40395, from the U.S. Department of Agriculture’s National Institute of Food and Agriculture. The use of Zeiss 980 confocal microscope was made available through the National Institute of Health grant S10D026702.

