## Supplemental Figures for "Key molecular and cellular events of table olive fruit abscission zone formation during natural maturation and after ethephon treatment"

**A**

| Gene (Accession number) | Gene (Annotation) | Primer direction | 5'→3' sequence |
|---|---|---|---|
| *OLEA9_A107450* | *EIF4A* | forward | GAATCCTCCAGCAGCTCGATT |
| | | reverse | AGTGCTCGCATGACCTTCTC |
| *OLEA9_A090870* | *DIR11* | forward | AACTGTCTGGGAAGTGGCAC |
| | | reverse | GCGATCAGGTCCGGTTGTTA |
| *OLEA9_A120253* | *LAC14* | forward | GCAACTGTGTTTGTTCCCTGG |
| | | reverse | TGCACAGTTTGTCCCTCCAG |
| *OLEA9_A121547* | *BG14* | forward | CATTCCGGCCAGATCTTGCT |
| | | reverse | GTAATCGAGTGGGACCTGGC |
| *OLEA9_A110794* | *BG13* | forward | AAAGTCTGCTCCCCTGTTGC |
| | | reverse | CATGGGCTTGACTACGGGTT |
| *OLEA9_A121155* | *PMEI* | forward | GGATTGCGTGGAGGAGTTGA |
| | | reverse | GGTGTCCTCGTCTGTCAAGG |
| *OLEA9_A020370* | *PL* | forward | GATGCGCCGGAAAGTGAATG |
| | | reverse | GTATGTGGAGGACGAGGCAC |
| *OLEA9_A112468* | *XTH24* | forward | CGCGGGAAGATAGTCGAAGG |
| | | reverse | CGGTTCCAGCAGAGTTTCCA |

**B**

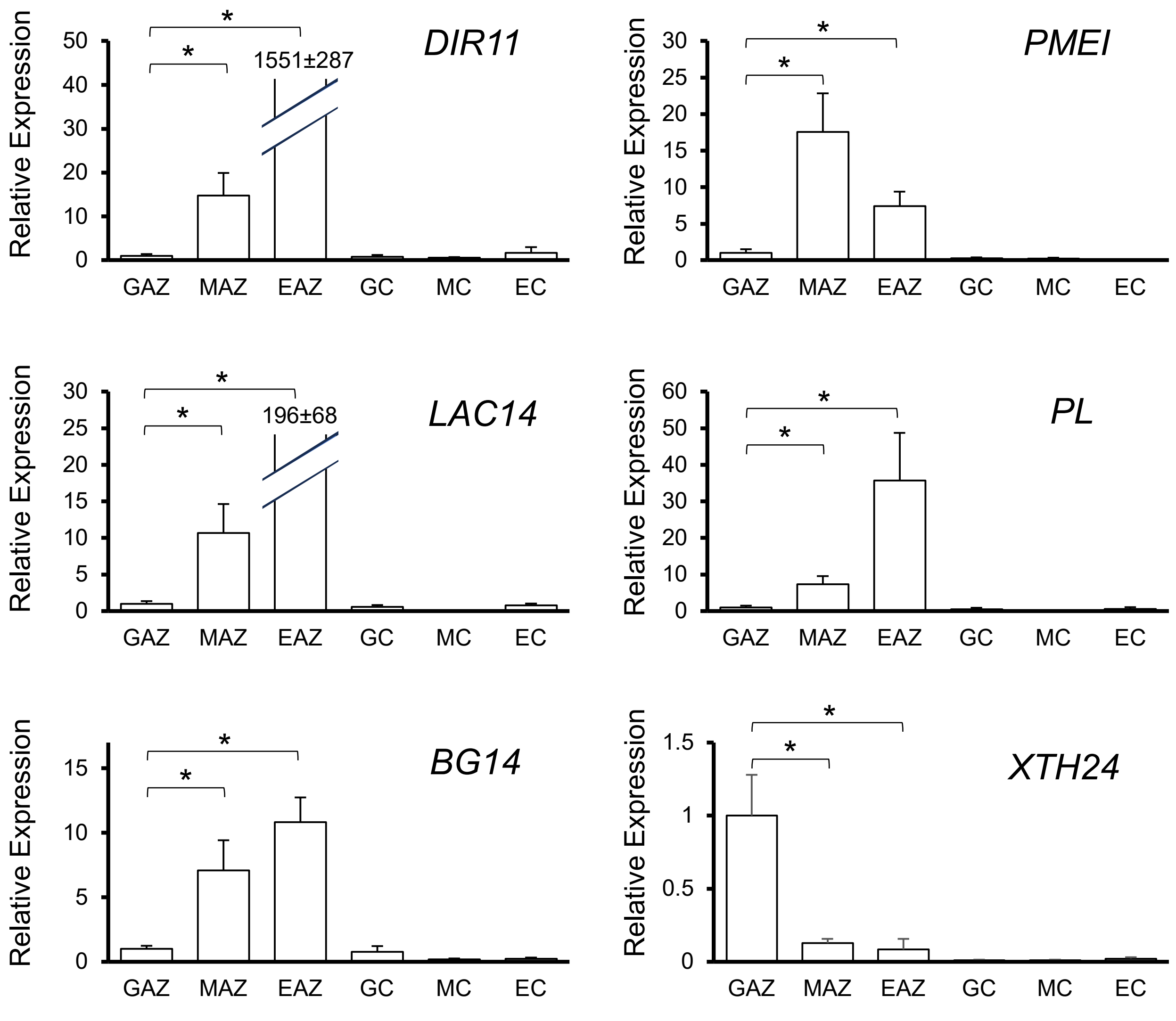


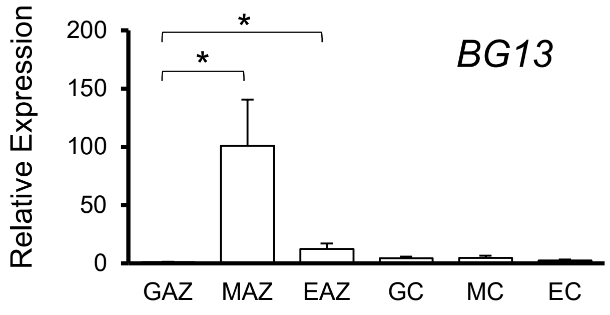


**Supplemental Figure 1**. **Real-time RT-PCR validation of selected DEG of transcriptome analysis.** (**A**) Primer sequences used in this analysis. *EIF4A* is the reference gene used in the relative quantification method. (**B**) Expression of selected genes in **Table 1-4** are validated by real-time RT-PCR.

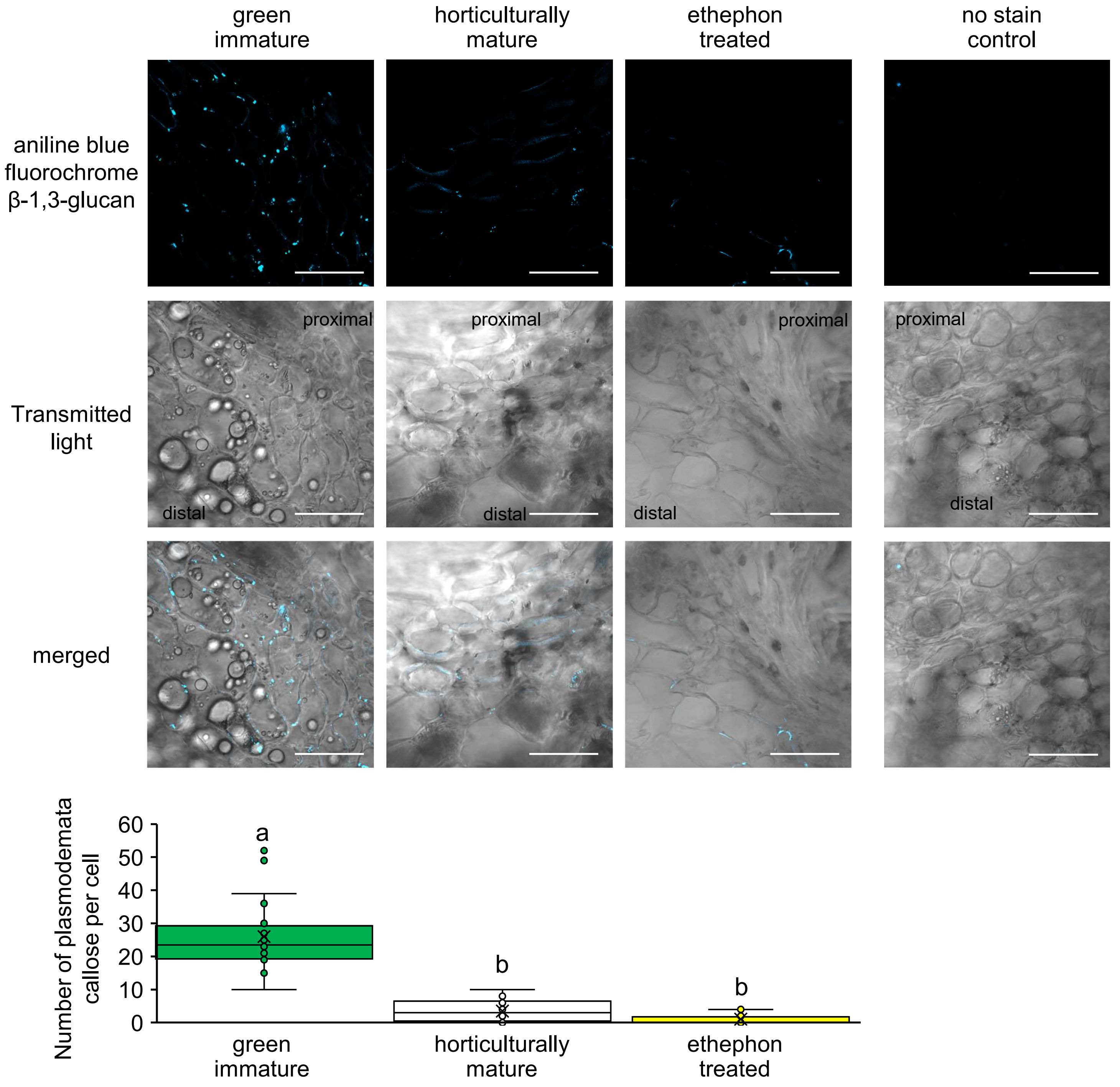


**Supplemental Figure 2. Live staining of callose by aniline blue fluorochrome.** In each image, five consecutive cells are counted for number of plasmodesmata and calculated for average number of plasmodesmata on each cell. No stain control is imaged using a green immature sample. Different letter indicates significant difference between groups ((N = 4 biological replicates and n = 20 cells for each group, one-way ANOVA, Tukey HSD, $p < 0.05$).

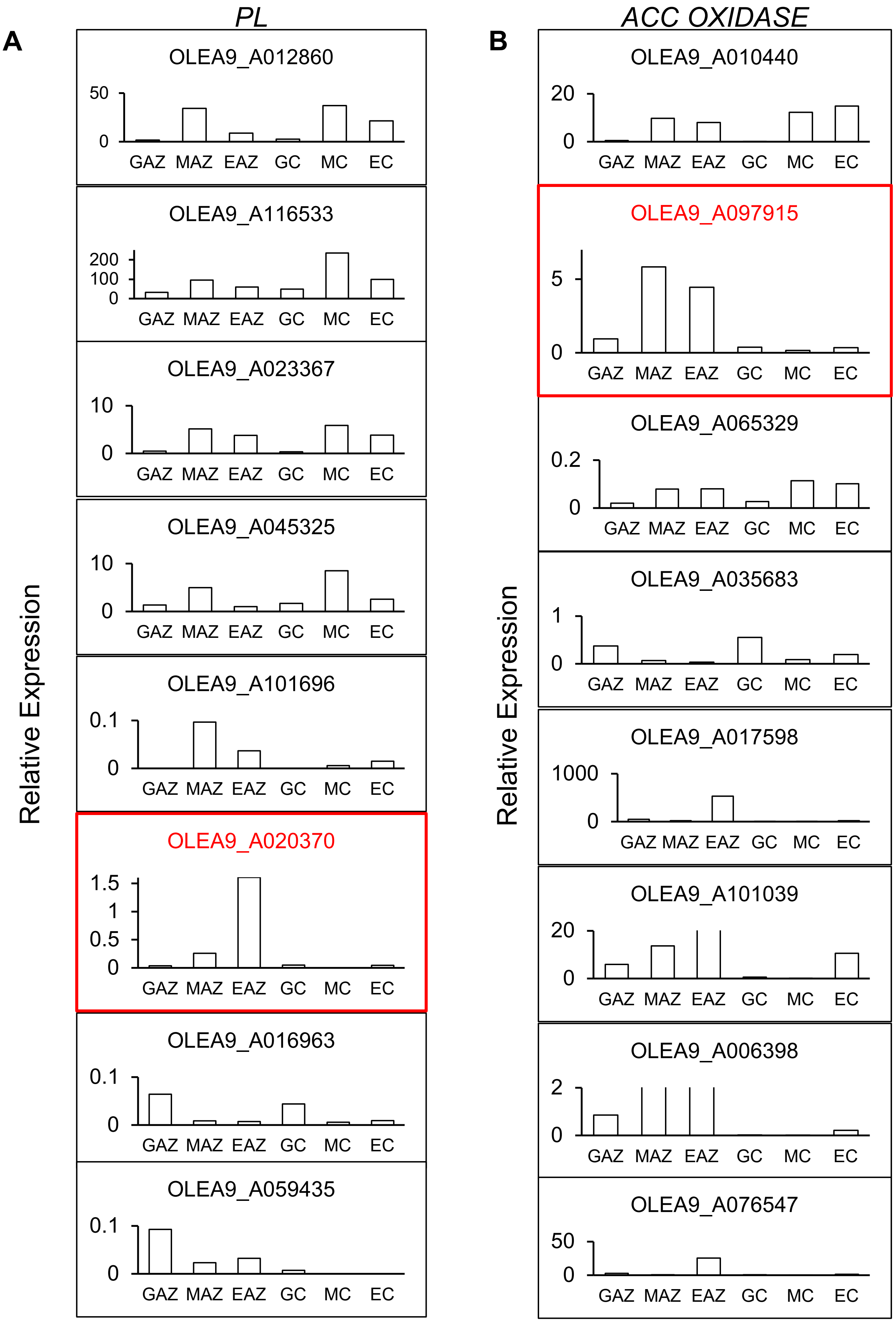


**Supplemental Figure 3. Identification of a FAZ-specific pectin lyase and ACC oxidase based on the subtraction method.** All the genes annotated as PL and ACC oxidases in DGE analysis of MAZvsGAZ are plotted by their mean FPKM value in six groups (GAZ, MAZ, EAZ, GC, MC, and EC). The genes OLEA9_A023070 and OLEA9_A097915 (highlighted in red) were identified among the 733 critical genes (**C**$_{\text{int}}$) of FAZ development by applying the subset formula as described in **Section 3.4** and in **Figure 4**, with a DGE screening threshold set to $|\log 2(\text{FoldChange})| \geq 1$ and an adjusted p-value (padj) of $\leq 0.05$. The Y axis scale was adjusted to show low expression genes.
