## Supplemental Tables for "Key molecular and cellular events of table olive fruit abscission zone formation during natural maturation and after ethephon treatment"

**Supplemental Table 1.** DEGs categorized as “cell wall modification and cell development”, shown in **Figure 4D**. The DEGs of **Table 2 – 4** and endomembrane trafficking related genes are also included in this table.

| UniProt Gene Name | GAZ | MAZ | EAZ | GC | MC | EC | Uniprot Homolog | Annotation | |
| --- | --- | --- | --- | --- | --- | --- | --- | --- | --- |
| OLEA9_A011623 | 0.022 | 0.000 | 0.000 | 0.020 | 0.002 | 0.003 | P05117 | *Solanum lycopersicum* | Polygalacturonase-2 |
| OLEA9_A019625 | 2.913 | 1.440 | 1.492 | 3.631 | 2.172 | 3.014 | Q9FVJ3 | *Arabidopsis thaliana* | ADP-ribosylation factor GTPase-activating protein AGD12 |
| OLEA9_A024326 | 0.424 | 0.166 | 0.031 | 0.035 | 0.000 | 0.000 | Q9LZR0 | *Arabidopsis thaliana* | Putative homeobox-leucine zipper protein ATHB-51 |
| OLEA9_A025777 | 0.055 | 0.191 | 0.398 | 0.045 | 0.055 | 0.138 | Q9S9M5 | *Arabidopsis thaliana* | Wall-associated receptor kinase-like 1 |
| OLEA9_A027294 | 0.174 | 0.070 | 0.083 | 0.136 | 0.047 | 0.082 | Q39817 | *Glycine max* | Calnexin homolog |
| OLEA9_A029961 | 0.064 | 0.007 | 0.000 | 0.003 | 0.000 | 0.000 | O64566 | *Arabidopsis thaliana* | Plant intracellular Ras-group-related LRR protein 6 |
| OLEA9_A030158 | 0.308 | 0.144 | 0.129 | 0.124 | 0.042 | 0.032 | Q1PEM5 | *Arabidopsis thaliana* | Proline-rich receptor-like protein kinase PERK3 |
| OLEA9_A032538 | 0.143 | 1.003 | 0.484 | 0.082 | 0.159 | 0.256 | P36401 | *Nicotiana tabacum* | Glucan endo-1,3-beta-glucosidase, acidic isoform PR-Q' |
| OLEA9_A034821 | 0.000 | 0.139 | 0.255 | 0.000 | 0.010 | 0.010 | Q67XC4 | *Arabidopsis thaliana* | Protein trichome birefringence-like 40 |
| OLEA9_A037399 | 0.541 | 0.196 | 0.200 | 0.464 | 0.303 | 0.269 | C0LGQ4 | *Arabidopsis thaliana* | Protein MALE DISCOVERER 2 |
| OLEA9_A038563 | 4.629 | 2.051 | 1.864 | 4.653 | 2.509 | 3.028 | P49967 | *Arabidopsis thaliana* | Signal recognition particle 54 kDa protein 3 |
| OLEA9_A039615 | 0.368 | 0.080 | 0.081 | 0.184 | 0.154 | 0.142 | Q8LEG3 | *Arabidopsis thaliana* | 65-kDa microtubule-associated protein 2 |
| OLEA9_A044345 | 1.350 | 0.613 | 0.608 | 1.114 | 0.722 | 0.680 | Q9LT71 | *Arabidopsis thaliana* | Reticulon-like protein B11 |
| OLEA9_A048423 | 4.452 | 9.182 | 18.213 | 2.810 | 3.736 | 4.129 | Q501H5 | *Arabidopsis thaliana* | Phosphatidylinositol/phosphatidylcholine transfer protein SFH13 |
| OLEA9_A050359 | 0.817 | 0.336 | 0.294 | 0.147 | 0.089 | 0.060 | Q9LY62 | *Arabidopsis thaliana* | Glycosylinositol phosphorylceramide mannosyl transferase 1 |
| OLEA9_A051307 | 0.010 | 0.118 | 0.122 | 0.000 | 0.011 | 0.026 | Q12700 | *Schwanniomyces occidentalis* | Glucan 1,3-beta-glucosidase |
| OLEA9_A055626 | 0.574 | 0.239 | 0.275 | 0.450 | 0.237 | 0.330 | Q39131 | *Arabidopsis thaliana* | Lamin-like protein |
| OLEA9_A056901 | 0.149 | 0.420 | 0.807 | 0.006 | 0.007 | 0.029 | Q10I20 | *Oryza sativa* subsp. *japonica* | Alpha-1,3-arabinosyltransferase XAT3 |
| OLEA9_A059281 | 2.293 | 7.547 | 61.360 | 0.278 | 0.334 | 0.440 | P13087 | *Synsepalum dulcificum* | Miraculin |
| OLEA9_A062618  (Supplemental Table 1 continues) | 0.808 | 0.202 | 0.169 | 0.947 | 0.582 | 0.390 | Q94A15 | *Arabidopsis thaliana* | Mannosyltransferase APTG1 |
| OLEA9_A064302 | 0.712 | 1.460 | 1.650 | 0.911 | 1.283 | 1.372 | P29675 | *Helianthus annuus* | Pollen-specific protein SF3 |
| OLEA9_A067026 | 9.325 | 19.100 | 24.034 | 13.217 | 24.301 | 14.518 | Q9C6S7 | *Arabidopsis thaliana* | Probable methyltransferase PMT20 |
| OLEA9_A074316 | 0.373 | 0.025 | 0.014 | 0.066 | 0.000 | 0.013 | O48716 | *Arabidopsis thaliana* | Protein JINGUBANG |
| OLEA9_A076305 | 0.174 | 0.068 | 0.060 | 0.130 | 0.058 | 0.057 | Q9LFW3 | *Arabidopsis thaliana* | COBRA-like protein 4 |
| OLEA9_A077201 | 0.069 | 0.595 | 0.548 | 0.043 | 0.052 | 0.074 | P26792 | *Daucus carota* | Beta-fructofuranosidase, insoluble isoenzyme 1 |
| OLEA9_A083606 | 4.768 | 9.662 | 12.124 | 3.366 | 3.479 | 5.994 | Q9LQN6 | *Arabidopsis thaliana* | Probable protein phosphatase 2C 4 |
| OLEA9_A084372 | 0.000 | 0.090 | 0.175 | 0.009 | 0.028 | 0.075 | P30278 | *Medicago sativa* | G2/mitotic-specific cyclin-2 (Fragment) |
| OLEA9_A084807 | 0.255 | 0.690 | 1.229 | 0.372 | 0.642 | 0.676 | Q9SGA8 | *Arabidopsis thaliana* | UDP-glycosyltransferase 83A1 |
| OLEA9_A085134 | 4.600 | 1.945 | 2.315 | 3.605 | 2.001 | 2.404 | Q52T38 | *Arabidopsis thaliana* | Protein S-acyltransferase 24 |
| OLEA9_A085704 | 1.047 | 0.248 | 0.262 | 1.020 | 0.786 | 0.641 | Q8RWB7 | *Arabidopsis thaliana* | Probable methyltransferase At1g29790 |
| OLEA9_A088370 | 0.788 | 0.138 | 0.118 | 0.021 | 0.000 | 0.012 | Q9XH43 | *Nicotiana tabacum* | CEN-like protein 2 |
| OLEA9_A089543 | 0.422 | 0.869 | 1.038 | 0.262 | 0.439 | 0.401 | Q66GR0 | *Arabidopsis thaliana* | Fasciclin-like arabinogalactan protein 17 |
| OLEA9_A089990 | 0.817 | 5.846 | 3.034 | 0.967 | 1.611 | 1.118 | Q8VYE5 | *Arabidopsis thaliana* | Glucan endo-1,3-beta-glucosidase 12 |
| OLEA9_A090031 | 0.257 | 0.102 | 0.085 | 0.122 | 0.043 | 0.067 | Q9C7U5 | *Arabidopsis thaliana* | Glucan endo-1,3-beta-glucosidase 2 |
| OLEA9_A090395 | 0.018 | 0.112 | 1.704 | 0.000 | 0.010 | 0.032 | O23349 | *Arabidopsis thaliana* | Primary amine oxidase 1 |
| OLEA9_A091862 | 0.129 | 0.004 | 0.000 | 0.022 | 0.000 | 0.000 | Q9SSE8 | *Arabidopsis thaliana* | Probable glycosyltransferase At3g07620 |
| OLEA9_A093757 | 0.078 | 0.000 | 0.012 | 0.004 | 0.000 | 0.000 | Q8GYX3 | *Arabidopsis thaliana* | Microtubule-associated protein 70-5 |
| OLEA9_A095511 | 0.300 | 2.542 | 1.022 | 0.023 | 0.041 | 0.067 | Q20BM9 | *Panax ginseng* | CASP-like protein 1 |
| OLEA9_A095734 | 0.014 | 0.107 | 0.262 | 0.024 | 0.045 | 0.037 | P46277 | *Medicago sativa* subsp. *varia* | G2/mitotic-specific cyclin-1 |
| OLEA9_A096439 | 0.516 | 0.028 | 0.083 | 0.039 | 0.000 | 0.000 | O48711 | *Arabidopsis thaliana* | Probable pectinesterase/pectinesterase inhibitor 12 |
| OLEA9_A101334 | 0.137 | 0.049 | 0.035 | 0.027 | 0.000 | 0.000 | Q9LJN4 | *Arabidopsis thaliana* | Probable beta-D-xylosidase 5 |
| OLEA9_A103252 | 0.079 | 0.017 | 0.007 | 0.016 | 0.000 | 0.000 | F4KEW8 | *Arabidopsis thaliana* | Protein NETWORKED 4A |
| OLEA9_A104351 | 6.158 | 12.734 | 15.024 | 7.279 | 13.183 | 12.536 | P29675 | *Helianthus annuus* | Pollen-specific protein SF3 |
| OLEA9_A113722 | 0.383 | 0.068 | 0.177 | 0.055 | 0.016 | 0.016 | Q9CAC1 | *Arabidopsis thaliana* | Endoglucanase 8 |
| OLEA9_A115562 | 0.132 | 0.020 | 0.008 | 0.125 | 0.042 | 0.025 | B9SA89 | *Ricinus communis* | CASP-like protein 2B1 |
| OLEA9_A115874 | 0.778 | 0.368 | 0.384 | 0.916 | 0.526 | 0.472 | P51566 | *Arabidopsis thaliana* | Serine/threonine-protein kinase AFC1 |
| OLEA9_A120547 | 0.022 | 0.151 | 0.125 | 0.027 | 0.114 | 0.039 | C0LGP9 | *Arabidopsis thaliana* | Probable leucine-rich repeat receptor-like protein kinase IMK3 |
| OLEA9_A120753 | 0.018 | 0.057 | 0.089 | 0.019 | 0.028 | 0.021 | O81905 | *Arabidopsis thaliana* | Receptor-like serine/threonine-protein kinase SD1-8 |
| OLEA9_NCA010457 | 0.139 | 0.011 | 0.013 | 0.089 | 0.000 | 0.013 | Q9LFT5 | *Arabidopsis thaliana* | EPIDERMAL PATTERNING FACTOR-like protein 1 |
| OLEA9_NCA013835  (Supplemental Table 1 continues) | 1.371 | 0.219 | 0.037 | 0.137 | 0.000 | 0.000 | Q7XXN8 | *Arabidopsis thaliana* | Small polypeptide DEVIL 16 |
| OLEA9_NCA023899 | 5.360 | 1.923 | 2.281 | 5.216 | 4.014 | 3.522 | Q5M759 | *Arabidopsis thaliana* | Vacuolar protein sorting-associated protein 22 homolog 1 |
| OLEA9_A012299 | 0.392 | 1.058 | 1.249 | 0.124 | 0.069 | 0.409 | Q9ZQG9 | *Arabidopsis thaliana* | Glucan endo-1,3-beta-glucosidase 14 |
| OLEA9_A017983 | 0.145 | 0.011 | 0.038 | 0.048 | 0.016 | 0.049 | Q8LGA1 | *Arabidopsis thaliana* | Cyclin-D4-1 |
| OLEA9_A019127 | 0.039 | 0.176 | 0.623 | 0.082 | 0.042 | 0.169 | O81906 | *Arabidopsis thaliana* | G-type lectin S-receptor-like serine/threonine-protein kinase B120 |
| OLEA9_A020370 | 0.037 | 0.257 | 1.604 | 0.051 | 0.000 | 0.046 | O24554 | *Zinnia violacea* | Pectate lyase |
| OLEA9_A021157 | 0.015 | 0.158 | 0.188 | 0.011 | 0.013 | 0.007 | P26792 | *Daucus carota* | Beta-fructofuranosidase, insoluble isoenzyme 1 |
| OLEA9_A023165 | 0.306 | 0.726 | 1.397 | 0.134 | 0.107 | 0.214 | Q9SS43 | *Arabidopsis thaliana* | Xylan glycosyltransferase MUCI21 |
| OLEA9_A028717 | 2.815 | 6.392 | 9.468 | 0.796 | 0.713 | 0.641 | Q9SHI2 | *Arabidopsis thaliana* | Leucine-rich repeat receptor-like serine/threonine-protein kinase At1g17230 |
| OLEA9_A036088 | 0.151 | 1.668 | 1.094 | 0.036 | 0.023 | 0.102 | A7PW81 | *Vitis vinifera* | Polygalacturonase inhibitor |
| OLEA9_A039241 | 0.054 | 0.232 | 0.206 | 0.101 | 0.160 | 0.086 | Q9FGQ6 | *Arabidopsis thaliana* | Microtubule-associated protein RP/EB family member 1C |
| OLEA9_A043949 | 0.414 | 0.129 | 0.051 | 0.092 | 0.110 | 0.094 | P35694 | *Glycine max* | Xyloglucan endotransglucosylase/hydrolase 2 |
| OLEA9_A045610 | 1.202 | 0.598 | 0.590 | 1.221 | 1.302 | 1.651 | Q9LDH3 | *Arabidopsis thaliana* | Probable sugar phosphate/phosphate translocator At1g12500 |
| OLEA9_A048969 | 0.379 | 0.039 | 0.000 | 0.019 | 0.046 | 0.044 | Q8GT41 | *Platanus acerifolia* | Putative invertase inhibitor |
| OLEA9_A052367 | 0.758 | 3.239 | 9.680 | 0.148 | 0.104 | 0.221 | A0A068Q721 | *Prunus mume* | Cytochrome P450 71AP13 |
| OLEA9_A060129 | 0.203 | 0.076 | 0.072 | 0.000 | 0.000 | 0.004 | Q9S7X6 | *Arabidopsis thaliana* | Thermospermine synthase ACAULIS5 |
| OLEA9_A066198 | 0.043 | 0.148 | 0.197 | 0.005 | 0.014 | 0.000 | Q41144 | *Ricinus communis* | Sugar carrier protein C |
| OLEA9_A068408 | 0.002 | 0.018 | 0.023 | 0.001 | 0.001 | 0.002 | F4JXC5 | *Arabidopsis thaliana* | Subtilisin-like protease SBT5.4 |
| OLEA9_A074219 | 0.113 | 0.417 | 0.491 | 0.023 | 0.019 | 0.186 | Q9ZQG9 | *Arabidopsis thaliana* | Glucan endo-1,3-beta-glucosidase 14 |
| OLEA9_A078401 | 0.936 | 0.383 | 0.467 | 0.637 | 0.428 | 0.800 | Q8L7M1 | *Arabidopsis thaliana* | Probable beta-1,3-galactosyltransferase 14 |
| OLEA9_A080734 | 0.167 | 0.659 | 1.679 | 0.211 | 0.038 | 0.607 | C0HJG8 | *Boswellia serrata* | Basic secretory protease (Fragments) |
| OLEA9_A095111 | 0.077 | 0.200 | 0.609 | 0.080 | 0.022 | 0.176 | Q9LMP1 | *Arabidopsis thaliana* | Wall-associated receptor kinase 2 |
| OLEA9_A095790 | 0.025 | 0.156 | 0.580 | 0.000 | 0.000 | 0.000 | Q9S9M5 | *Arabidopsis thaliana* | Wall-associated receptor kinase-like 1 |
| OLEA9_A096076 | 0.351 | 1.479 | 11.648 | 0.052 | 0.007 | 0.145 | Q38865 | *Arabidopsis thaliana* | Expansin-A6 |
| OLEA9_A100467 | 0.005 | 0.016 | 0.016 | 0.000 | 0.000 | 0.000 | Q9C9L5 | *Arabidopsis thaliana* | Wall-associated receptor kinase-like 9 |
| OLEA9_A101028 | 0.314 | 1.014 | 3.575 | 0.000 | 0.000 | 0.059 | Q9FF86 | *Arabidopsis thaliana* | BAHD acyltransferase DCR |
| OLEA9_A104376 | 0.167 | 1.212 | 0.407 | 0.015 | 0.000 | 0.019 | Q6NKQ9 | *Arabidopsis thaliana* | Plasmodesmata-located protein 8 |
| OLEA9_A106190 | 0.019 | 0.163 | 0.357 | 0.022 | 0.000 | 0.000 | Q8VZU9 | *Arabidopsis thaliana* | Derlin-1 |
| OLEA9_A108156 | 0.011 | 0.049 | 0.089 | 0.000 | 0.000 | 0.000 | Q66GQ6 | *Arabidopsis thaliana* | Type I inositol polyphosphate 5-phosphatase 5 |
| OLEA9_A110481 | 0.726 | 0.270 | 0.327 | 0.810 | 0.508 | 1.102 | Q8S8Q6 | *Arabidopsis thaliana* | Tetraspanin-8 |
| OLEA9_A110794 | 0.875 | 6.072 | 2.224 | 1.324 | 1.383 | 1.042 | Q9FJU9 | *Arabidopsis thaliana* | Glucan endo-1,3-beta-glucosidase 13 |
| OLEA9_A110954 | 0.239 | 0.608 | 1.000 | 0.047 | 0.038 | 0.067 | Q9SW48 | *Arabidopsis thaliana* | Probable alkaline/neutral invertase B |
| OLEA9_A112468 | 1.206 | 0.217 | 0.247 | 0.065 | 0.026 | 0.077 | P24806 | *Arabidopsis thaliana* | Xyloglucan endotransglucosylase/hydrolase protein 24 |
| OLEA9_A117455 | 0.112 | 1.675 | 1.131 | 0.006 | 0.000 | 0.008 | Q8L868 | *Arabidopsis thaliana* | Glucan endo-1,3-beta-glucosidase 11 |
| OLEA9_A118587 | 4.017 | 10.934 | 9.447 | 3.749 | 3.380 | 4.874 | Q8VYR7 | *Arabidopsis thaliana* | Boron transporter 1 |
| OLEA9_A119718 | 0.838 | 2.082 | 2.977 | 0.238 | 0.022 | 0.749 | none |  | PF04554:Extensin-like region |
| OLEA9_A121155 | 0.414 | 5.408 | 1.230 | 0.000 | 0.000 | 0.000 | P17407 | *Daucus carota* | 21 kDa protein |
| OLEA9_A121547 | 12.490 | 40.730 | 65.947 | 2.138 | 1.431 | 2.801 | Q9ZQG9 | *Arabidopsis thaliana* | Glucan endo-1,3-beta-glucosidase 14 |
| OLEA9_NCA022907 | 0.222 | 0.577 | 1.512 | 0.037 | 0.006 | 0.044 | Q9SIN2 | *Arabidopsis thaliana* | Protein trichome birefringence-like 39 |

(Supplemental Table 1 continues)

**Supplemental Table 2.** DEGs categorized as “hormonal regulation pathway”, the category in **Figure 4D**.

| UniProt Gene Name | GAZ | MAZ | EAZ | GC | MC | EC | Uniprot Homolog | Annotation | |
| --- | --- | --- | --- | --- | --- | --- | --- | --- | --- |
| OLEA9_A011873 | 0.225 | 0.499 | 1.166 | 0.229 | 0.247 | 0.304 | K4CI52 | *Solanum lycopersicum* | Abscisic acid 8'-hydroxylase CYP707A2 |
| OLEA9_A011891 | 0.386 | 0.053 | 0.056 | 0.065 | 0.007 | 0.017 | Q9ZUX1 | *Arabidopsis thaliana* | Cytochrome P450 94C1 |
| OLEA9_A017469 | 0.032 | 0.097 | 0.151 | 0.022 | 0.028 | 0.053 | Q9LP24 | *Arabidopsis thaliana* | Probable leucine-rich repeat receptor-like protein kinase At1g35710 |
| OLEA9_A018884 | 1.873 | 0.403 | 0.786 | 2.504 | 1.447 | 1.876 | Q9M210 | *Arabidopsis thaliana* | Ethylene-responsive transcription factor ERF035 |
| OLEA9_A089178 | 0.000 | 0.124 | 0.082 | 0.000 | 0.030 | 0.026 | Q9FH54 | *Arabidopsis thaliana* | Ethylene-responsive transcription factor ERF114 |
| OLEA9_A114398 | 0.095 | 2.796 | 0.900 | 0.030 | 0.054 | 0.139 | Q9LYU3 | *Arabidopsis thaliana* | Ethylene-responsive transcription factor ERF113 |
| OLEA9_A040878 | 0.000 | 0.787 | 0.917 | 0.000 | 0.056 | 0.142 | Q9C9I8 | *Arabidopsis thaliana* | Ethylene-responsive transcription factor ERF020 |
| OLEA9_A121142 | 0.171 | 0.043 | 0.016 | 0.000 | 0.046 | 0.000 | Q9SK03 | *Arabidopsis thaliana* | Ethylene-responsive transcription factor RAP2-7 |
| OLEA9_A027325 | 0.360 | 0.789 | 1.483 | 0.293 | 0.542 | 0.568 | Q94AH6 | *Arabidopsis thaliana* | Cullin-1 |
| OLEA9_A030374 | 0.056 | 0.000 | 0.000 | 0.015 | 0.000 | 0.000 | P32068 | *Arabidopsis thaliana* | Anthranilate synthase alpha subunit 1, chloroplastic |
| OLEA9_A031842 | 0.711 | 0.172 | 0.019 | 0.434 | 0.252 | 0.100 | P32295 | *Vigna radiata var. radiata* | Indole-3-acetic acid-induced protein ARG7 |
| OLEA9_A037900 | 0.081 | 0.315 | 0.548 | 0.020 | 0.095 | 0.116 | Q7Y0W3 | *Oryza sativa* subsp. *indica* | Two-component response regulator EHD1 |
| OLEA9_A052023 | 0.155 | 0.323 | 0.320 | 0.157 | 0.204 | 0.244 | P32746 | *Arabidopsis thaliana* | Dihydroorotate dehydrogenase (quinone), mitochondrial |
| OLEA9_A060621 | 0.157 | 0.000 | 0.000 | 0.035 | 0.000 | 0.000 | Q93Z37 | *Arabidopsis thaliana* | Protein BIG GRAIN 1-like E |
| OLEA9_A062531 | 4.310 | 1.279 | 2.203 | 6.079 | 3.199 | 3.626 | Q9FGQ7 | *Arabidopsis thaliana* | Cyclin-D3-2 |
| OLEA9_A062668 | 0.042 | 0.002 | 0.000 | 0.008 | 0.000 | 0.003 | Q9FWX7 | *Arabidopsis thaliana* | ABC transporter B family member 11 |
| OLEA9_A087005 | 0.251 | 0.017 | 0.018 | 0.016 | 0.000 | 0.000 | Q9ZPS9 | *Arabidopsis thaliana* | Serine/threonine-protein kinase BRI1-like 2 |
| OLEA9_A088141 | 0.632 | 0.189 | 0.094 | 0.062 | 0.023 | 0.018 | Q8RUN2 | *Arabidopsis thaliana* | Cytokinin riboside 5'-monophosphate phosphoribohydrolase LOG1 |
| OLEA9_A090614 | 0.489 | 0.000 | 0.000 | 0.098 | 0.000 | 0.000 | O22227 | *Arabidopsis thaliana* | Protein MIZU-KUSSEI 1 |
| OLEA9_A095940 | 1.881 | 4.597 | 12.099 | 0.675 | 0.890 | 1.046 | Q9LS19 | *Arabidopsis thaliana* | Protein DETOXIFICATION 30 |
| OLEA9_A110023 | 0.242 | 0.092 | 0.102 | 0.170 | 0.069 | 0.067 | Q9C826 | *Arabidopsis thaliana* | Xanthoxin dehydrogenase |
| OLEA9_A111536 | 0.233 | 0.045 | 0.024 | 0.029 | 0.000 | 0.016 | Q9C6S5 | *Arabidopsis thaliana* | Probable polyamine transporter At1g31830 |
| OLEA9_A113932 | 0.045 | 0.004 | 0.004 | 0.011 | 0.000 | 0.008 | C0LGF5 | *Arabidopsis thaliana* | LRR receptor-like serine/threonine-protein kinase RGI5 |
| OLEA9_NCA001329A | 0.038 | 0.004 | 0.011 | 0.005 | 0.000 | 0.001 | Q9SJ74 | *Arabidopsis thaliana* | Cytochrome b561 and DOMON domain-containing protein At2g04850 |
| OLEA9_A006108 | 0.188 | 0.052 | 0.077 | 0.000 | 0.016 | 0.010 | O22213 | *Arabidopsis thaliana* | Cytokinin dehydrogenase 1 |
| OLEA9_A029114 | 0.053 | 0.164 | 0.497 | 0.007 | 0.000 | 0.021 | Q93WV4 | *Arabidopsis thaliana* | WRKY transcription factor 71 |
| OLEA9_A032871 | 0.119 | 0.365 | 0.831 | 0.007 | 0.000 | 0.088 | O80939 | *Arabidopsis thaliana* | L-type lectin-domain containing receptor kinase IV.1 |
| OLEA9_A033778 | 1.108 | 0.190 | 0.502 | 1.113 | 0.822 | 1.211 | O49561 | *Arabidopsis thaliana* | Gibberellin 2-beta-dioxygenase 8 |
| OLEA9_A035617 | 0.139 | 0.441 | 0.325 | 0.093 | 0.092 | 0.071 | Q9LPF6 | *Arabidopsis thaliana* | Probable purine permease 11 |
| OLEA9_A035916 | 0.014 | 0.128 | 0.353 | 0.029 | 0.013 | 0.114 | Q9FG13 | *Arabidopsis thaliana* | Probable carboxylesterase 15 |
| OLEA9_A038150 | 0.116 | 0.568 | 0.475 | 0.128 | 0.057 | 0.050 | Q9SIH5 | *Arabidopsis thaliana* | Probable F-box protein At2g36090 |
| OLEA9_A052395 | 0.006 | 0.318 | 1.605 | 0.015 | 0.000 | 0.018 | O80920 | *Arabidopsis thaliana* | Abscisic acid receptor PYL4 |
| OLEA9_A055552 | 0.109 | 0.617 | 0.521 | 0.000 | 0.000 | 0.050 | O22150 | *Arabidopsis thaliana* | Auxin-responsive protein SAUR36 |
| OLEA9_A055796 | 0.012 | 0.082 | 1.275 | 0.010 | 0.005 | 0.051 | Q9C768 | *Arabidopsis thaliana* | UDP-glycosyltransferase 76B1 |
| OLEA9_A086168 | 0.005 | 0.113 | 1.083 | 0.000 | 0.000 | 0.017 | Q94AH6 | *Arabidopsis thaliana* | Cullin-1 |
| OLEA9_A086522 | 0.541 | 0.214 | 0.091 | 0.009 | 0.018 | 0.010 | Q9FFD0 | *Arabidopsis thaliana* | Auxin efflux carrier component 5 |
| OLEA9_A094455 | 0.112 | 0.971 | 1.085 | 0.006 | 0.006 | 0.019 | Q8RUN2 | *Arabidopsis thaliana* | Cytokinin riboside 5'-monophosphate phosphoribohydrolase LOG1 |
| OLEA9_A097915 | 0.954 | 5.851 | 4.461 | 0.394 | 0.177 | 0.354 | Q84MB3 | *Arabidopsis thaliana* | 1-aminocyclopropane-1-carboxylate oxidase homolog 1 |
| OLEA9_A100712 | 0.091 | 0.230 | 3.919 | 0.023 | 0.000 | 0.151 | Q8S9K8 | *Arabidopsis thaliana* | Methylesterase 10 |
| OLEA9_A107517 | 0.285 | 0.017 | 0.023 | 0.000 | 0.000 | 0.006 | Q9LSQ4 | *Arabidopsis thaliana* | Indole-3-acetic acid-amido synthetase GH3.6 |
| OLEA9_A113908 | 0.271 | 0.570 | 1.518 | 0.125 | 0.121 | 0.273 | Q94F62 | *Arabidopsis thaliana* | BRASSINOSTEROID INSENSITIVE 1-associated receptor kinase 1 |

(Supplemental Table 2 continues)

**Supplemental Table 3**. DEGs categorized as “transcription factors”, as shown in in **Figure 4D**.

| UniProt Gene Name | GAZ | MAZ | EAZ | GC | MC | EC | Uniprot Homolog | Annotation | |
| --- | --- | --- | --- | --- | --- | --- | --- | --- | --- |
| OLEA9_A003876 | 0.407 | 0.140 | 0.000 | 0.076 | 0.000 | 0.000 | Q9XGX0 | *Arabidopsis thaliana* | Protein SHORT INTERNODES |
| OLEA9_A009811 | 1.676 | 0.588 | 0.618 | 0.035 | 0.000 | 0.021 | O82268 | *Arabidopsis thaliana* | Protein LIGHT-DEPENDENT SHORT HYPOCOTYLS 3 |
| OLEA9_A011472 | 0.343 | 0.112 | 0.128 | 0.046 | 0.000 | 0.020 | Q9LQZ7 | *Arabidopsis thaliana* | B-box zinc finger protein 21 |
| OLEA9_A016457 | 1.722 | 0.617 | 0.638 | 2.072 | 1.213 | 1.298 | Q9MAH8 | *Arabidopsis thaliana* | Transcription factor TCP3 |
| OLEA9_A030413 | 1.812 | 0.584 | 0.700 | 2.731 | 2.594 | 1.552 | A2XA73 | *Oryza sativa* subsp. *indica* | Growth-regulating factor 1 |
| OLEA9_A032680 | 1.629 | 0.591 | 0.758 | 1.347 | 0.879 | 0.996 | Q9SMW7 | *Arabidopsis thaliana* | Basic transcription factor 3 |
| OLEA9_A032845 | 13.481 | 33.339 | 41.878 | 11.109 | 17.731 | 21.686 | A8MQN2 | *Arabidopsis thaliana* | Protein LNK1 |
| OLEA9_A034033 | 0.559 | 0.064 | 0.061 | 0.047 | 0.000 | 0.000 | Q93WY4 | *Arabidopsis thaliana* | Probable WRKY transcription factor 12 |
| OLEA9_A034399 | 0.411 | 0.130 | 0.036 | 0.051 | 0.000 | 0.000 | Q9FY93 | *Arabidopsis thaliana* | NAC domain-containing protein 83 |
| OLEA9_A041141 | 0.653 | 0.046 | 0.196 | 0.028 | 0.000 | 0.000 | Q9ZPW3 | *Arabidopsis thaliana* | Transcription factor HBI1 |
| OLEA9_A043201 | 1.856 | 0.853 | 0.738 | 0.363 | 0.257 | 0.342 | O22210 | *Arabidopsis thaliana* | Transcription factor MYBC1 |
| OLEA9_A053735 | 0.064 | 0.000 | 0.000 | 0.020 | 0.000 | 0.000 | Q1PEY6 | *Arabidopsis thaliana* | EPIDERMAL PATTERNING FACTOR-like protein 6 |
| OLEA9_A054613 | 17.658 | 8.807 | 8.061 | 19.780 | 14.270 | 11.583 | Q8LC03 | *Arabidopsis thaliana* | Homeobox-leucine zipper protein ATHB-13 |
| OLEA9_A066209 | 0.054 | 0.009 | 0.008 | 0.033 | 0.010 | 0.005 | Q9SVY1 | *Arabidopsis thaliana* | Zinc finger protein WIP2 |
| OLEA9_A075808 | 0.662 | 0.285 | 0.151 | 0.519 | 0.511 | 0.264 | B5X570 | *Arabidopsis thaliana* | NAC domain-containing protein 14 |
| OLEA9_A076953 | 15.447 | 7.513 | 7.786 | 20.328 | 14.585 | 11.830 | Q93V43 | *Arabidopsis thaliana* | Transcription factor TCP2 |
| OLEA9_A084376 | 0.463 | 0.034 | 0.036 | 0.108 | 0.030 | 0.019 | Q8S3M3 | *Elaeis oleifera* | NOI-like protein |
| OLEA9_A091124 | 0.061 | 0.235 | 3.400 | 0.011 | 0.038 | 0.073 | Q9FYA2 | *Arabidopsis thaliana* | Probable WRKY transcription factor 75 |
| OLEA9_A092784 | 2.332 | 0.945 | 0.916 | 1.822 | 0.946 | 1.309 | Q9LZS0 | *Arabidopsis thaliana* | Trihelix transcription factor PTL |
| OLEA9_A096644 | 0.439 | 0.171 | 0.187 | 0.737 | 0.454 | 0.368 | Q9CAA9 | *Arabidopsis thaliana* | Transcription factor bHLH49 |
| OLEA9_A098303 | 14.422 | 5.778 | 6.606 | 22.041 | 10.676 | 11.652 | O04136 | *Malus domestica* | Homeobox protein knotted-1-like 3 |
| OLEA9_A099601 | 0.145 | 0.007 | 0.000 | 0.016 | 0.000 | 0.000 | Q9C9V6 | *Arabidopsis thaliana* | BTB/POZ domain-containing protein At1g67900 |
| OLEA9_A099889 | 0.269 | 0.000 | 0.000 | 0.024 | 0.011 | 0.000 | Q38741 | *Antirrhinum majus* | Squamosa promoter-binding protein 1 |
| OLEA9_A100507 | 0.138 | 0.919 | 2.002 | 0.000 | 0.011 | 0.076 | Q10MB4 | *Oryza sativa* subsp. *japonica* | Transcription factor MYB2 |
| OLEA9_A100940  (Supplemental Table 3 continues) | 2.390 | 0.914 | 1.008 | 2.063 | 1.515 | 1.180 | Q9FKZ1 | *Arabidopsis thaliana* | Probable disease resistance protein At5g66900 |
| OLEA9_A101530 | 2.404 | 0.697 | 0.982 | 3.349 | 1.796 | 2.288 | Q9ZVP8 | *Arabidopsis thaliana* | NAC domain-containing protein 35 |
| OLEA9_A103270 | 0.805 | 0.389 | 0.329 | 0.806 | 0.441 | 0.422 | Q94A57 | *Arabidopsis thaliana* | Protein PHR1-LIKE 2 |
| OLEA9_A103310 | 0.618 | 0.201 | 0.111 | 0.722 | 0.433 | 0.348 | Q9C7U7 | *Arabidopsis thaliana* | Transcription factor MYB20 |
| OLEA9_A105792 | 0.793 | 0.392 | 0.207 | 0.589 | 0.288 | 0.313 | Q39262 | *Arabidopsis thaliana* | Zinc finger protein 3 |
| OLEA9_A105929 | 0.005 | 0.000 | 0.000 | 0.001 | 0.001 | 0.000 | Q9SLL2 | *Arabidopsis thaliana* | Protein BIG GRAIN 1-like B |
| OLEA9_A106241 | 0.516 | 0.178 | 0.216 | 0.325 | 0.150 | 0.141 | Q9LS28 | *Arabidopsis thaliana* | Transcription factor GTE12 |
| OLEA9_A106912 | 0.182 | 0.416 | 0.810 | 0.016 | 0.046 | 0.031 | Q9C519 | *Arabidopsis thaliana* | WRKY transcription factor 6 |
| OLEA9_A109666 | 0.511 | 0.066 | 0.038 | 0.075 | 0.000 | 0.027 | Q6X7J9 | *Arabidopsis thaliana* | WUSCHEL-related homeobox 4 |
| OLEA9_A110187 | 0.422 | 0.120 | 0.181 | 0.374 | 0.223 | 0.249 | B3VTV7 | *Vitis vinifera* | Transcription factor MYB60 |
| OLEA9_A112242 | 0.452 | 0.918 | 1.350 | 0.508 | 0.755 | 0.949 | Q1PEU4 | *Arabidopsis thaliana* | Pentatricopeptide repeat-containing protein At2g44880 |
| OLEA9_A114597 | 1.037 | 0.111 | 0.344 | 0.061 | 0.000 | 0.029 | Q39265 | *Arabidopsis thaliana* | Zinc finger protein 6 |
| OLEA9_A116554 | 0.000 | 0.081 | 0.253 | 0.000 | 0.008 | 0.027 | P41151 | *Arabidopsis thaliana* | Heat stress transcription factor A-1a |
| OLEA9_A117458 | 0.049 | 0.003 | 0.006 | 0.035 | 0.006 | 0.003 | Q04088 | *Arabidopsis thaliana* | Probable transcription factor PosF21 |
| OLEA9_A118460 | 0.459 | 0.219 | 0.054 | 0.051 | 0.000 | 0.000 | Q8S9N6 | *Arabidopsis thaliana* | Homeobox-leucine zipper protein ATHB-17 |
| OLEA9_A118473 | 0.075 | 0.490 | 0.479 | 0.000 | 0.073 | 0.039 | Q338B0 | *Oryza sativa* subsp. *japonica* | Heat stress transcription factor A-2c |
| OLEA9_A118522 | 16.109 | 7.138 | 7.299 | 18.201 | 9.333 | 11.222 | B4XT64 | *Oryza sativa* subsp. *japonica* | Protein NARROW LEAF 1 |
| OLEA9_NCA015710 | 0.294 | 0.104 | 0.099 | 0.206 | 0.160 | 0.144 | Q9SWG3 | *Arabidopsis thaliana* | Protein FAR-RED IMPAIRED RESPONSE 1 |
| OLEA9_NCA016551 | 0.006 | 0.038 | 0.075 | 0.002 | 0.002 | 0.004 | Q9SZL8 | *Arabidopsis thaliana* | Protein FAR1-RELATED SEQUENCE 5 |
| OLEA9_A005888 | 0.226 | 0.000 | 0.025 | 0.000 | 0.000 | 0.000 | Q6X7J9 | *Arabidopsis thaliana* | WUSCHEL-related homeobox 4 |
| OLEA9_A018761 | 19.502 | 49.270 | 55.163 | 21.050 | 33.147 | 12.675 | Q9M886 | *Arabidopsis thaliana* | LOB domain-containing protein 41 |
| OLEA9_A023080 | 0.740 | 0.315 | 0.284 | 1.148 | 1.661 | 0.818 | Q9SKY9 | *Arabidopsis thaliana* | Transcription repressor OFP16 |
| OLEA9_A041186 | 0.009 | 0.258 | 0.157 | 0.010 | 0.000 | 0.010 | O81821 | *Arabidopsis thaliana* | Heat stress transcription factor A-1b |
| OLEA9_A048746 | 0.212 | 1.441 | 1.926 | 0.032 | 0.016 | 0.037 | Q8H102 | *Arabidopsis thaliana* | Transcription factor bHLH128 |
| OLEA9_A059417 | 0.522 | 0.000 | 0.124 | 0.000 | 0.000 | 0.019 | Q9SN12 | *Arabidopsis thaliana* | Transcription factor MYB77 |
| OLEA9_A061167 | 0.070 | 0.561 | 3.466 | 0.009 | 0.000 | 0.021 | Q9FGY3 | *Arabidopsis thaliana* | Transcription factor MYB78 |
| OLEA9_A063433 | 0.446 | 1.620 | 18.150 | 0.014 | 0.000 | 0.097 | Q39264 | *Arabidopsis thaliana* | Zinc finger protein 5 |
| OLEA9_A077146 | 0.153 | 0.387 | 1.169 | 0.009 | 0.000 | 0.005 | Q93WU9 | *Arabidopsis thaliana* | Probable WRKY transcription factor 51 |
| OLEA9_A081410 | 0.388 | 1.118 | 1.033 | 0.026 | 0.000 | 0.177 | Q9LQF0 | *Arabidopsis thaliana* | Transcription factor TCP23 |
| OLEA9_A088204 | 0.005 | 0.331 | 3.336 | 0.005 | 0.000 | 0.020 | Q9LY00 | *Arabidopsis thaliana* | Probable WRKY transcription factor 70 |
| OLEA9_A088209 | 0.564 | 2.514 | 2.702 | 0.148 | 0.040 | 0.283 | D0PX88 | *Lotus japonicus* | bHLH transcription factor RHL1 |
| OLEA9_A088921 | 0.245 | 0.520 | 0.717 | 0.036 | 0.000 | 0.008 | Q84TE9 | *Arabidopsis thaliana* | Dof zinc finger protein DOF5.3 |
| OLEA9_A091564 | 0.010 | 0.083 | 1.972 | 0.000 | 0.000 | 0.008 | Q9FNI3 | *Arabidopsis thaliana* | B3 domain-containing protein At5g06250 |
| OLEA9_A092147 | 1.074 | 8.530 | 6.527 | 0.036 | 0.000 | 0.301 | Q9LZP8 | *Arabidopsis thaliana* | bZIP transcription factor 53 |
| OLEA9_A093147 | 0.026 | 0.148 | 0.416 | 0.016 | 0.000 | 0.039 | Q93WU9 | *Arabidopsis thaliana* | Probable WRKY transcription factor 51 |
| OLEA9_A096297 | 0.499 | 2.414 | 8.294 | 0.000 | 0.000 | 0.045 | Q9ZWM9 | *Arabidopsis thaliana* | AP2/ERF and B3 domain-containing transcription factor RAV1 |
| OLEA9_A107048 | 0.153 | 0.699 | 0.578 | 0.016 | 0.008 | 0.009 | O24160 | *Nicotiana tabacum* | TGACG-sequence-specific DNA-binding protein TGA-2.1 |
| OLEA9_A107773 | 0.218 | 0.598 | 0.758 | 0.053 | 0.000 | 0.032 | Q43385 | *Arabidopsis thaliana* | Dof zinc finger protein DOF3.7 |
| OLEA9_A107901 | 0.523 | 1.738 | 1.681 | 0.030 | 0.000 | 0.009 | Q9FYJ6 | *Arabidopsis thaliana* | Transcription factor bHLH111 |
| OLEA9_A117856 | 0.000 | 0.210 | 1.316 | 0.000 | 0.000 | 0.032 | Q9FYA2 | *Arabidopsis thaliana* | Probable WRKY transcription factor 75 |
| OLEA9_A119562 | 1.628 | 3.543 | 3.464 | 0.589 | 0.343 | 0.397 | Q8W4F0 | *Arabidopsis thaliana* | Protein DA1-related 1 |
| OLEA9_NCA017547 | 1.638 | 3.590 | 4.221 | 1.529 | 1.510 | 1.905 | A0A0P0X9Z7 | *Oryza sativa* subsp. *japonica* | Cysteine-tryptophan domain-containing zinc finger protein 7 |

(Supplementary Table 3 continues)

**Supplemental Table 4.** DEGs categorized as “Secondary metabolism related, other than phenylpropanoid pathway”, as shown in **Figure 4D**.

| UniProt Gene Name | GAZ | MAZ | EAZ | GC | MC | EC | Uniprot Homolog | Annotation | |
| --- | --- | --- | --- | --- | --- | --- | --- | --- | --- |
| OLEA9_A002312 | 0.812 | 0.268 | 0.346 | 1.279 | 0.685 | 0.700 | H2DH17 | *Panax ginseng* | Cytochrome P450 CYP749A22 |
| OLEA9_A018071 | 35.600 | 76.902 | 91.857 | 48.667 | 86.832 | 84.779 | P49352 | *Lupinus albus* | Farnesyl pyrophosphate synthase 2 |
| OLEA9_A018094 | 0.108 | 0.000 | 0.007 | 0.014 | 0.000 | 0.000 | A0A1W6GW32 | *Salvia miltiorrhiza* | (-)-5-epieremophilene synthase STPS1 |
| OLEA9_A066825 | 0.000 | 0.134 | 0.051 | 0.000 | 0.008 | 0.019 | A0A075FA51 | *Marrubium vulgare* | (+)-copalyl diphosphate synthase 3, chloroplastic |
| OLEA9_A072936 | 0.086 | 0.592 | 0.789 | 0.069 | 0.179 | 0.259 | Q6V4H0 | *Catharanthus roseus* | 8-hydroxygeraniol dehydrogenase |
| OLEA9_A080790 | 1.720 | 0.635 | 0.871 | 2.372 | 1.402 | 1.795 | Q9SJB4 | *Arabidopsis thaliana* | GDSL esterase/lipase At2g04570 |
| OLEA9_A080903 | 0.465 | 0.151 | 0.208 | 0.576 | 0.440 | 0.348 | Q949X0 | *Arabidopsis thaliana* | Palmitoyl-monogalactosyldiacylglycerol delta-7 desaturase, chloroplastic |
| OLEA9_A098609 | 1.241 | 0.571 | 0.593 | 1.259 | 0.721 | 0.676 | B4FHU1 | *Zea mays* | 15-cis-zeta-carotene isomerase, chloroplastic |
| OLEA9_A103529 | 2.098 | 1.039 | 1.059 | 1.632 | 0.840 | 0.965 | Q9M4W3 | *Catharanthus roseus* | 2-C-methyl-D-erythritol 2,4-cyclodiphosphate synthase, chloroplastic |
| OLEA9_A115579 | 0.349 | 0.129 | 0.144 | 0.407 | 0.308 | 0.226 | G3FIN8 | *Manihot esculenta* | Linamarin synthase 1 |
| OLEA9_A115672 | 0.128 | 0.030 | 0.032 | 0.195 | 0.188 | 0.116 | P46285 | *Triticum aestivum* | Sedoheptulose-1,7-bisphosphatase, chloroplastic |
| OLEA9_A116177 | 0.010 | 0.308 | 0.147 | 0.000 | 0.005 | 0.013 | A0A1X9IRQ7 | *Isodon rubescens* | Copalyl diphosphate synthase 1, chloroplastic |
| OLEA9_A000829 | 0.000 | 0.131 | 0.076 | 0.000 | 0.000 | 0.094 | A6YIH8 | *Hyoscyamus muticus* | Premnaspirodiene oxygenase |
| OLEA9_A001711 | 0.085 | 1.203 | 9.943 | 0.000 | 0.000 | 0.056 | Q9SVG5 | *Arabidopsis thaliana* | Berberine bridge enzyme-like 18 |
| OLEA9_A007424 | 0.368 | 0.155 | 0.018 | 0.011 | 0.000 | 0.057 | Q84ND0 | *Antirrhinum majus* | Tricyclene synthase Oc15, chloroplastic |
| OLEA9_A013817 | 0.042 | 0.423 | 0.336 | 0.000 | 0.000 | 0.020 | A0A1W6GW32 | *Salvia miltiorrhiza* | (-)-5-epieremophilene synthase STPS1 |
| OLEA9_A025606 | 0.033 | 0.428 | 10.479 | 0.011 | 0.000 | 0.025 | Q8VYH2 | *Arabidopsis thaliana* | Squalene epoxidase 3 |
| OLEA9_A037830 | 0.371 | 0.891 | 1.249 | 0.153 | 0.105 | 0.172 | Q9LMI7 | *Arabidopsis thaliana* | Putative acyl-coenzyme A oxidase 3.2, peroxisomal |
| OLEA9_A046907 | 0.027 | 0.068 | 0.255 | 0.022 | 0.013 | 0.048 | Q6V4H0 | *Catharanthus roseus* | 8-hydroxygeraniol dehydrogenase |
| OLEA9_A051613 | 0.374 | 2.792 | 12.000 | 0.796 | 0.387 | 0.451 | K4CEE8 | *Solanum lycopersicum* | Beta-amyrin 28-monooxygenase |
| OLEA9_A058687 | 0.000 | 0.493 | 0.982 | 0.000 | 0.000 | 0.000 | A0A1W6GW32 | *Salvia miltiorrhiza* | (-)-5-epieremophilene synthase STPS1 |
| OLEA9_A058720 | 0.174 | 0.815 | 1.123 | 0.030 | 0.000 | 0.029 | A0A2K9RFZ8 | *Vitex agnus-castus* | Syn-copalyl diphosphate synthase TPS3, chloroplastic |
| OLEA9_A087759 | 0.142 | 0.372 | 2.440 | 0.007 | 0.000 | 0.000 | Q9FF86 | *Arabidopsis thaliana* | BAHD acyltransferase DCR |
| OLEA9_A089679 | 0. 009 | 0.026 | 0.039 | 0.017 | 0.019 | 0.011 | Q6BE25 | *Cucurbita pepo* | Cycloartenol synthase |
| OLEA9_A093824 | 0.000 | 0.307 | 0.831 | 0.000 | 0.000 | 0.064 | A0A068Q6L2 | *Prunus mume* | Cytochrome P450 736A117 |
| OLEA9_A098480 | 0.000 | 0.047 | 0.209 | 0.000 | 0.000 | 0.006 | A0A1D6HSP4 | *Zea mays* | Dimethylnonatriene synthase |
| OLEA9_A099177 | 1.396 | 3.807 | 3.565 | 0.418 | 0.177 | 0.697 | A0A068Q6L2 | *Prunus mume* | Cytochrome P450 736A117 |
| OLEA9_A101358 | 0.065 | 0.004 | 0.005 | 0.004 | 0.000 | 0.011 | O81191 | *Salvia officinalis* | 1,8-cineole synthase, chloroplastic |
| OLEA9_A108518 | 0.048 | 0.897 | 4.047 | 0.043 | 0.015 | 0.063 | O82146 | *Panax ginseng* | Beta-amyrin synthase 2 |
| OLEA9_A114699 | 0.000 | 0.075 | 0.047 | 0.002 | 0.000 | 0.014 | B2HIN2 | *Mycobacterium marinum* | Long-chain-fatty-acid--AMP ligase FadD26 |

(Supplemental Table 4 continues)

**Supplemental Table 5.** DEGs categorized as “amino acid metabolism and transport”, as shown in **Figure 4D**.

| UniProt Gene Name | GAZ | MAZ | EAZ | GC | MC | EC | Uniprot Homolog | Annotation | |
| --- | --- | --- | --- | --- | --- | --- | --- | --- | --- |
| OLEA9_A004664 | 0.724 | 0.183 | 0.074 | 0.105 | 0.000 | 0.000 | Q9FHH5 | *Arabidopsis thaliana* | Protein GLUTAMINE DUMPER 3 |
| OLEA9_A062454 | 7.229 | 14.898 | 20.755 | 9.450 | 18.041 | 14.466 | Q9LV03 | *Arabidopsis thaliana* | Glutamate synthase 1 [NADH], chloroplastic |
| OLEA9_A105593 | 0.550 | 0.121 | 0.004 | 0.030 | 0.000 | 0.000 | P92934 | *Arabidopsis thaliana* | Amino acid permease 6 |
| OLEA9_A108127 | 0.580 | 1.393 | 3.876 | 0.789 | 1.419 | 1.483 | Q93XM7 | *Arabidopsis thaliana* | Mitochondrial carnitine/acylcarnitine carrier-like protein |
| OLEA9_A110035 | 1.014 | 0.373 | 0.521 | 0.901 | 0.577 | 0.848 | Q0WMZ5 | *Arabidopsis thaliana* | Outer envelope pore protein 16-2, chloroplastic |
| OLEA9_A110240 | 0.635 | 0.235 | 0.142 | 0.124 | 0.095 | 0.057 | Q39134 | *Arabidopsis thaliana* | Amino acid permease 3 |
| OLEA9_A001729 | 0.103 | 0.029 | 0.000 | 0.000 | 0.000 | 0.000 | P92934 | *Arabidopsis thaliana* | Amino acid permease 6 |
| OLEA9_A005349 | 0.777 | 1.892 | 3.324 | 0.337 | 0.324 | 0.557 | Q9FKS8 | *Arabidopsis thaliana* | Lysine histidine transporter 1 |
| OLEA9_A009187 | 0.060 | 0.202 | 0.311 | 0.049 | 0.013 | 0.040 | Q9FKS8 | *Arabidopsis thaliana* | Lysine histidine transporter 1 |
| OLEA9_A025415 | 0.796 | 2.751 | 9.598 | 0.000 | 0.000 | 0.051 | Q9FL41 | *Arabidopsis thaliana* | WAT1-related protein At5g07050 |
| OLEA9_A036181 | 2.026 | 4.388 | 7.937 | 1.929 | 1.008 | 1.763 | Q43314 | *Arabidopsis thaliana* | Glutamate dehydrogenase 1 |
| OLEA9_A059616 | 0.126 | 0.360 | 0.480 | 0.026 | 0.025 | 0.087 | Q680I5 | *Arabidopsis thaliana* | Glutathione hydrolase 2 |
| OLEA9_A070043 | 7.508 | 21.722 | 30.218 | 4.466 | 4.108 | 5.324 | Q9FKS8 | *Arabidopsis thaliana* | Lysine histidine transporter 1 |
| OLEA9_A076335 | 2.000 | 7.838 | 8.256 | 0.407 | 0.335 | 0.782 | Q9SW07 | *Arabidopsis thaliana* | Protein GLUTAMINE DUMPER 2 |
| OLEA9_A103595 | 0.353 | 0.995 | 1.976 | 0.085 | 0.045 | 0.114 | Q84MA5 | *Arabidopsis thaliana* | Cationic amino acid transporter 1 |
| OLEA9_A110766 | 0.078 | 0.384 | 0.726 | 0.029 | 0.009 | 0.068 | Q9ZPR7 | *Arabidopsis thaliana* | Ureide permease 1 |

**Supplemental Table 6.** DEGs categorized as “stress response and plant immunity pathway” related, as shown in **Figure 4D**.

| UniProt Gene Name | GAZ | MAZ | EAZ | GC | MC | EC | Uniprot Homolog | Annotation | |
| --- | --- | --- | --- | --- | --- | --- | --- | --- | --- |
| OLEA9_A001961 | 0.155 | 0.017 | 0.037 | 0.113 | 0.021 | 0.051 | Q9LUS3 | *Arabidopsis thaliana* | Pentatricopeptide repeat-containing protein At3g16610 |
| OLEA9_A003521 | 0.269 | 0.097 | 0.097 | 0.046 | 0.013 | 0.009 | P93400 | *Nicotiana tabacum* | Phospholipase D alpha 1 |
| OLEA9_A008324 | 6.144 | 1.808 | 2.261 | 6.872 | 4.534 | 4.518 | Q9LYD3 | *Arabidopsis thaliana* | Dehydration-responsive element-binding protein 3 |
| OLEA9_A013437 | 1.922 | 0.847 | 0.906 | 2.278 | 1.379 | 1.235 | P17598 | *Gossypium hirsutum* | Catalase isozyme 1 |
| OLEA9_A014307 | 0.129 | 0.008 | 0.014 | 0.159 | 0.045 | 0.053 | Q8LDP4 | *Arabidopsis thaliana* | Peptidyl-prolyl cis-trans isomerase CYP19-4 |
| OLEA9_A016218 | 0.127 | 0.379 | 1.732 | 0.108 | 0.135 | 0.202 | Q08480 | *Oryza sativa* subsp. *japonica* | Adenylate kinase 4 |
| OLEA9_A016987 | 2.720 | 0.791 | 1.198 | 3.843 | 2.250 | 2.522 | Q8RXN0 | *Arabidopsis thaliana* | ABC transporter G family member 11 |
| OLEA9_A020387 | 0.353 | 0.042 | 0.000 | 0.308 | 0.065 | 0.068 | O04450 | *Arabidopsis thaliana* | T-complex protein 1 subunit epsilon |
| OLEA9_A026559 | 45.576 | 20.539 | 18.185 | 53.939 | 30.847 | 34.621 | Q9MAH3 | *Arabidopsis thaliana* | Protein DJ-1 homolog B |
| OLEA9_A034100 | 0.204 | 0.038 | 0.000 | 0.022 | 0.000 | 0.009 | Q9S7T8 | *Arabidopsis thaliana* | Serpin-ZX |
| OLEA9_A035536 | 1.351 | 0.559 | 0.209 | 0.329 | 0.187 | 0.093 | Q9FI03 | *Arabidopsis thaliana* | NDR1/HIN1-like protein 26 |
| OLEA9_A037877 | 2.182 | 5.197 | 5.899 | 1.909 | 2.984 | 3.784 | O81832 | *Arabidopsis thaliana* | G-type lectin S-receptor-like serine/threonine-protein kinase At4g27290 |
| OLEA9_A040165 | 0.200 | 0.087 | 0.026 | 0.019 | 0.000 | 0.006 | Q40588 | *Nicotiana tabacum* | L-ascorbate oxidase |
| OLEA9_A042247 | 0.023 | 0.156 | 0.283 | 0.043 | 0.058 | 0.085 | Q9LZX1 | *Arabidopsis thaliana* | Protein LURP-one-related 15 |
| OLEA9_A046831 | 0.154 | 0.000 | 0.000 | 0.023 | 0.000 | 0.000 | O49696 | *Arabidopsis thaliana* | Aluminum-activated malate transporter 12 |
| OLEA9_A049330 | 0.000 | 0.142 | 0.190 | 0.000 | 0.022 | 0.034 | Q9LV60 | *Arabidopsis thaliana* | Cysteine-rich repeat secretory protein 55 |
| OLEA9_A050365 | 0.151 | 0.024 | 0.000 | 0.035 | 0.000 | 0.000 | none |  | PF00582:Universal stress protein family |
| OLEA9_A051239 | 15.952 | 7.018 | 6.706 | 18.979 | 10.253 | 9.411 | O49931 | *Pisum sativum* | Protein TIC 55, chloroplastic |
| OLEA9_A052865 | 0.601 | 1.243 | 1.246 | 0.763 | 1.182 | 1.076 | Q940Q2 | *Arabidopsis thaliana* | Pentatricopeptide repeat-containing protein At1g07590, mitochondrial |
| OLEA9_A056412 | 0.279 | 0.039 | 0.000 | 0.299 | 0.115 | 0.134 | Q9SVL6 | *Arabidopsis thaliana* | Cold-regulated 413 plasma membrane protein 2 |
| OLEA9_A057751 | 0.093 | 0.639 | 0.712 | 0.325 | 0.338 | 0.591 | Q852K5 | *Oryza sativa* subsp. *japonica* | Zinc finger A20 and AN1 domain-containing stress-associated protein 6 |
| OLEA9_A058719 | 3.457 | 9.739 | 8.840 | 2.466 | 2.854 | 3.774 | Q9ZV43 | *Arabidopsis thaliana* | Protein CHROMATIN REMODELING 8 |
| OLEA9_A059775 | 0.959 | 0.374 | 0.371 | 1.154 | 0.806 | 0.686 | Q9SSG3 | *Arabidopsis thaliana* | HIPL1 protein |
| OLEA9_A069896 | 0.659 | 0.227 | 0.237 | 1.019 | 0.751 | 0.532 | F4KGN5 | *Arabidopsis thaliana* | Solute carrier family 40 member 2 |
| OLEA9_A073725 | 0.468 | 3.877 | 4.911 | 0.686 | 1.061 | 0.776 | Q8S9L6 | *Arabidopsis thaliana* | Cysteine-rich receptor-like protein kinase 29 |
| OLEA9_A077372  (Supplemental Table 6 continues) | 0.033 | 0.004 | 0.000 | 0.018 | 0.002 | 0.002 | Q69SP5 | *Oryza sativa* subsp. *japonica* | LRR receptor-like serine/threonine-protein kinase ER1 |
| OLEA9_A083916 | 0.000 | 0.612 | 21.765 | 0.000 | 0.019 | 0.053 | Q9ZP41 | *Citrus jambhiri* | EG45-like domain containing protein |
| OLEA9_A089316 | 0.661 | 0.254 | 0.175 | 0.314 | 0.192 | 0.203 | Q9LI89 | *Arabidopsis thaliana* | F-box/kelch-repeat protein At3g27150 |
| OLEA9_A090932 | 0.039 | 0.284 | 0.224 | 0.116 | 0.221 | 0.147 | Q9M9A5 | *Arabidopsis thaliana* | Protein PLANT CADMIUM RESISTANCE 6 |
| OLEA9_A097119 | 0.175 | 0.648 | 0.501 | 0.257 | 0.394 | 0.312 | Q9FLS9 | *Arabidopsis thaliana* | Pentatricopeptide repeat-containing protein At5g61800 |
| OLEA9_A098106 | 1.794 | 0.885 | 0.620 | 0.240 | 0.090 | 0.227 | Q67YC0 | *Arabidopsis thaliana* | Inorganic pyrophosphatase 1 |
| OLEA9_A111095 | 0.462 | 1.861 | 1.214 | 0.305 | 0.495 | 0.332 | Q9LHQ6 | *Arabidopsis thaliana* | Organic cation/carnitine transporter 4 |
| OLEA9_A112344 | 0.218 | 0.000 | 0.046 | 0.196 | 0.079 | 0.120 | Q9NWL6 | *Homo sapiens* | Asparagine synthetase domain-containing protein 1 |
| OLEA9_A118635 | 0.299 | 0.026 | 0.014 | 0.133 | 0.011 | 0.028 | O82176 | *Arabidopsis thaliana* | Protein EXORDIUM-like 7 |
| OLEA9_NCA004599 | 0.670 | 0.332 | 0.277 | 0.460 | 0.253 | 0.249 | Q8W4I6 | *Arabidopsis thaliana* | GTP-binding protein BRASSINAZOLE INSENSITIVE PALE GREEN 2, chloroplastic |
| OLEA9_NCA006497 | 1.750 | 0.796 | 0.734 | 2.165 | 1.332 | 1.557 | P17801 | *Zea mays* | Putative receptor protein kinase ZmPK1 |
| OLEA9_NCA009450 | 0.767 | 0.159 | 0.232 | 0.565 | 0.470 | 0.254 | Q93ZH2 | *Arabidopsis thaliana* | Nuclear transcription factor Y subunit A-3 |
| OLEA9_NCA017212 | 0.734 | 0.355 | 0.275 | 0.495 | 0.376 | 0.275 | P48534 | *Pisum sativum* | L-ascorbate peroxidase, cytosolic |
| OLEA9_A008246 | 0.009 | 0.725 | 1.474 | 0.000 | 0.000 | 0.024 | Q9SX25 | *Arabidopsis thaliana* | Probable carboxylesterase 6 |
| OLEA9_A011769 | 0.330 | 1.089 | 10.442 | 0.047 | 0.000 | 0.089 | Q93WF6 | *Arabidopsis thaliana* | Protein SENESCENCE-ASSOCIATED GENE 21, mitochondrial |
| OLEA9_A015861 | 0.642 | 0.193 | 0.139 | 0.270 | 0.345 | 0.114 | Q9SXA1 | *Arabidopsis thaliana* | Phosphatidylinositol 4-kinase alpha 1 |
| OLEA9_A024386 | 0.079 | 4.914 | 12.494 | 0.013 | 0.000 | 0.014 | P22195 | *Arachis hypogaea* | Cationic peroxidase 1 |
| OLEA9_A024871 | 0.032 | 0.907 | 3.090 | 0.000 | 0.000 | 0.019 | Q8L8Z8 | *Arabidopsis thaliana* | Monothiol glutaredoxin-S2 |
| OLEA9_A026743 | 0.000 | 0.193 | 0.683 | 0.000 | 0.016 | 0.000 | Q9CAS1 | *Arabidopsis thaliana* | Thioredoxin H8 |
| OLEA9_A027157 | 0.425 | 11.173 | 1.792 | 0.045 | 0.000 | 0.046 | Q9LS46 | *Arabidopsis thaliana* | Aluminum-activated malate transporter 9 |
| OLEA9_A030695 | 0.000 | 0.026 | 0.070 | 0.000 | 0.000 | 0.010 | Q9SUQ7 | *Arabidopsis thaliana* | Cation/H(+) antiporter 17 |
| OLEA9_A030765 | 0.062 | 0.286 | 0.162 | 0.044 | 0.032 | 0.045 | C0LGP4 | *Arabidopsis thaliana* | Probable LRR receptor-like serine/threonine-protein kinase At3g47570 |
| OLEA9_A036711 | 0.000 | 0.080 | 0.726 | 0.000 | 0.000 | 0.030 | P87027 | *Schizosaccharomyces pombe* | Septum-promoting GTP-binding protein 1 |
| OLEA9_A042762 | 0.422 | 0.068 | 0.145 | 0.053 | 0.006 | 0.069 | Q8LGM7 | *Solanum lycopersicum* | Molybdenum cofactor sulfurase |
| OLEA9_A043586 | 0.376 | 1.211 | 7.889 | 0.075 | 0.014 | 0.061 | Q9SZG1 | *Arabidopsis thaliana* | Glycosyltransferase 6 |
| OLEA9_A047398 | 0.446 | 1.429 | 1.682 | 0.449 | 0.389 | 0.659 | Q9T050 | *Arabidopsis thaliana* | SPX domain-containing membrane protein At4g11810 |
| OLEA9_A048278 | 0.069 | 0.851 | 1.252 | 0.000 | 0.000 | 0.025 | Q9FN75 | *Arabidopsis thaliana* | AAA-ATPase At5g17760 |
| OLEA9_A048798 | 0.000 | 0.207 | 0.237 | 0.000 | 0.000 | 0.046 | A0A068Q6L2 | *Prunus mume* | Cytochrome P450 736A117 |
| OLEA9_A048843  (Supplemental Table 6 continues) | 0.006 | 0. 141 | 4.243 | 0.000 | 0.000 | 0.000 | P22195 | *Arachis hypogaea* | Cationic peroxidase 1 |
| OLEA9_A056751 | 0.000 | 0.193 | 0.243 | 0.022 | 0.010 | 0.063 | Q9LIK7 | *Arabidopsis thaliana* | Putative calcium-transporting ATPase 13, plasma membrane-type |
| OLEA9_A057532 | 0.060 | 0.200 | 0.315 | 0.081 | 0.037 | 0.078 | Q9SYD6 | *Arabidopsis thaliana* | Protein DETOXIFICATION 42 |
| OLEA9_A058819 | 1.758 | 3.716 | 3.783 | 1.029 | 0.499 | 1.896 | Q9LJJ7 | *Arabidopsis thaliana* | AAA-ATPase At3g28580 |
| OLEA9_A061082 | 0.000 | 0.905 | 2.057 | 0.000 | 0.000 | 0.048 | Q9LV60 | *Arabidopsis thaliana* | Cysteine-rich repeat secretory protein 55 |
| OLEA9_A063487 | 0.050 | 0.172 | 0.310 | 0.066 | 0.063 | 0.072 | O81832 | *Arabidopsis thaliana* | G-type lectin S-receptor-like serine/threonine-protein kinase At4g27290 |
| OLEA9_A064208 | 10.601 | 4.626 | 3.885 | 10.539 | 10.880 | 6.071 | F4I9E1 | *Arabidopsis thaliana* | Protein NUCLEAR FUSION DEFECTIVE 4 |
| OLEA9_A067613 | 0.112 | 0.282 | 0.245 | 0.023 | 0.000 | 0.010 | Q9SJL0 | *Arabidopsis thaliana* | UDP-glycosyltransferase 86A1 |
| OLEA9_A073443 | 0.012 | 0.202 | 0.111 | 0.014 | 0.000 | 0.016 | Q9LIK7 | *Arabidopsis thaliana* | Putative calcium-transporting ATPase 13, plasma membrane-type |
| OLEA9_A074682 | 0.023 | 0.091 | 1.283 | 0.009 | 0.003 | 0.000 | Q9SV13 | *Arabidopsis thaliana* | Sulfate transporter 3.1 |
| OLEA9_A077420 | 1.330 | 4.860 | 14.405 | 0.074 | 0.000 | 0.080 | O50001 | *Prunus armeniaca* | Major allergen Pru ar 1 |
| OLEA9_A077498 | 0.031 | 0.189 | 0.653 | 0.013 | 0.012 | 0.068 | Q9SXB4 | *Arabidopsis thaliana* | G-type lectin S-receptor-like serine/threonine-protein kinase At1g11300 |
| OLEA9_A080650 | 0.047 | 0.252 | 0.296 | 0.011 | 0.008 | 0.003 | Q8VZG8 | *Arabidopsis thaliana* | MDIS1-interacting receptor like kinase 2 |
| OLEA9_A096723 | 0.090 | 0.201 | 0.292 | 0.094 | 0.084 | 0.107 | Q9LMX4 | *Arabidopsis thaliana* | Probable inactive purple acid phosphatase 1 |
| OLEA9_A096826 | 0.111 | 0.636 | 0.882 | 0.000 | 0.000 | 0.000 | Q93ZF5 | *Arabidopsis thaliana* | Phosphate transporter PHO1 homolog 1 |
| OLEA9_A100270 | 0.091 | 0.526 | 0.597 | 0.024 | 0.015 | 0.010 | Q9M9Q6 | *Arabidopsis thaliana* | Serine carboxypeptidase-like 50 |
| OLEA9_A106152 | 0.136 | 0.572 | 2.137 | 0.095 | 0.000 | 0.039 | Q9ZVX4 | *Arabidopsis thaliana* | UDP-glycosyltransferase 90A1 |
| OLEA9_A120394 | 0.702 | 1.433 | 2.098 | 0.769 | 0.778 | 0.708 | O64477 | *Arabidopsis thaliana* | G-type lectin S-receptor-like serine/threonine-protein kinase At2g19130 |
| OLEA9_NCA012876 | 0.043 | 0.492 | 0.478 | 0.048 | 0.051 | 0.026 | P30221 | *Solanum lycopersicum* | 17.8 kDa class I heat shock protein |
| OLEA9_A035270 | 15.811 | 33.762 | 35.968 | 17.053 | 23.789 | 22.702 | Q76CU2 | *Nicotiana tabacum* | Pleiotropic drug resistance protein 1 |
| OLEA9_A002846 | 0.148 | 2.849 | 1.738 | 0.004 | 0.000 | 0.000 | Q9FX85 | *Arabidopsis thaliana* | Peroxidase 10 |
| OLEA9_A051698 | 0.000 | 0.206 | 0.145 | 0.000 | 0.000 | 0.021 | A9ZPJ7 | *Hordeum vulgare* | Agmatine coumaroyltransferase-2 |
| OLEA9_A074071 | 0.102 | 1.220 | 0.977 | 0.000 | 0.000 | 0.000 | Q9FX85 | *Arabidopsis thaliana* | Peroxidase 10 |
| OLEA9_A002739 | 0.653 | 0.280 | 0.245 | 0.652 | 0.394 | 0.371 | Q9FKZ1 | *Arabidopsis thaliana* | Probable disease resistance protein At5g66900 |
| OLEA9_A015218 | 0.939 | 2.365 | 3.447 | 0.836 | 1.258 | 1.636 | Q9LYW9 | *Arabidopsis thaliana* | DnaJ protein P58IPK homolog |
| OLEA9_A020942 | 0.383 | 0.013 | 0.044 | 0.004 | 0.000 | 0.000 | Q3E9B5 | *Arabidopsis thaliana* | Protein NRT1/ PTR FAMILY 7.1 |
| OLEA9_A029093 | 2.898 | 0.219 | 1.103 | 0.174 | 0.060 | 0.034 | Q39088 | *Arabidopsis thaliana* | Dof zinc finger protein DOF3.4 |
| OLEA9_A031883 | 0.000 | 0.056 | 0.053 | 0.000 | 0.004 | 0.021 | Q9SD62 | *Arabidopsis thaliana* | Putative receptor-like protein kinase At3g47110 |
| OLEA9_A044091 | 0.000 | 0.295 | 0.464 | 0.000 | 0.021 | 0.035 | P0C035 | *Arabidopsis thaliana* | RING-H2 finger protein ATL60 |
| OLEA9_A048901  (Supplemental Table 6 continues) | 0.219 | 0.532 | 3.179 | 0.040 | 0.107 | 0.084 | Q9LTC0 | *Arabidopsis thaliana* | Probable serine/threonine-protein kinase PBL19 |
| OLEA9_A049063 | 1.966 | 0.765 | 0.839 | 2.087 | 1.228 | 1.923 | Q9LHT0 | *Arabidopsis thaliana* | Tropinone reductase homolog At5g06060 |
| OLEA9_A052956 | 0.057 | 0.016 | 0.005 | 0.030 | 0.029 | 0.022 | O82777 | *Solanum lycopersicum* | Subtilisin-like protease SBT3 |
| OLEA9_A053133 | 0.608 | 0.072 | 0.249 | 0.432 | 0.198 | 0.212 | F4I082 | *Arabidopsis thaliana* | Non-specific lipid transfer protein GPI-anchored 6 |
| OLEA9_A057478 | 10.249 | 28.240 | 28.955 | 14.392 | 17.432 | 27.422 | Q9ZQ96 | *Arabidopsis thaliana* | UDP-glycosyltransferase 73C3 |
| OLEA9_A057750 | 3.727 | 1.731 | 1.895 | 4.570 | 3.076 | 3.271 | Q7XA39 | *Solanum bulbocastanum* | Putative disease resistance protein RGA4 |
| OLEA9_A089177 | 0.000 | 1.538 | 1.556 | 0.000 | 0.028 | 0.097 | P80211 | *Amaranthus caudatus* | Trypsin/subtilisin inhibitor |
| OLEA9_A091359 | 0.221 | 0.484 | 0.619 | 0.177 | 0.348 | 0.274 | Q9LVI6 | *Arabidopsis thaliana* | Probable inactive receptor kinase RLK902 |
| OLEA9_A091665 | 0.174 | 0.060 | 0.016 | 0.011 | 0.002 | 0.004 | Q9SHI3 | *Arabidopsis thaliana* | Receptor-like protein 2 |
| OLEA9_A097040 | 0.651 | 0.310 | 0.276 | 0.587 | 0.355 | 0.392 | Q9FFI2 | *Arabidopsis thaliana* | Autophagy protein 5 |
| OLEA9_A100005 | 0.283 | 0.033 | 0.066 | 0.500 | 0.365 | 0.310 | Q1MX30 | *Oryza sativa* subsp. *indica* | Receptor kinase-like protein Xa21 |
| OLEA9_A109478 | 0.301 | 0.705 | 0.658 | 0.308 | 0.411 | 0.541 | Q9SX38 | *Arabidopsis thaliana* | Putative disease resistance protein At1g50180 |
| OLEA9_A114936 | 0.153 | 0.072 | 0.068 | 0.064 | 0.016 | 0.033 | Q9LNV9 | *Arabidopsis thaliana* | Receptor-like protein 1 |
| OLEA9_A115666 | 216.644 | 48.886 | 95.760 | 314.486 | 164.124 | 193.925 | P85524 | *Actinidia deliciosa* | Kirola |
| OLEA9_NCA002642 | 0.170 | 1.538 | 1.478 | 0.000 | 0.021 | 0.249 | P37118 | *Solanum melongena* | Cytochrome P450 71A2 |
| OLEA9_NCA004336 | 1.024 | 2.614 | 2.298 | 1.259 | 1.474 | 1.876 | C0LGN2 | *Arabidopsis thaliana* | Probable leucine-rich repeat receptor-like serine/threonine-protein kinase At3g14840 |
| OLEA9_NCA004337 | 3.368 | 6.895 | 7.483 | 4.347 | 4.952 | 7.591 | C0LGN2 | *Arabidopsis thaliana* | Probable leucine-rich repeat receptor-like serine/threonine-protein kinase At3g14840 |
| OLEA9_A011276 | 0.000 | 0.175 | 0.665 | 0.000 | 0.000 | 0.000 | P86971 | *Fagopyrum tataricum* | Trypsin inhibitor |
| OLEA9_A015286 | 0.130 | 0.265 | 0.363 | 0.042 | 0.006 | 0.098 | O64793 | *Arabidopsis thaliana* | G-type lectin S-receptor-like serine/threonine-protein kinase At1g67520 |
| OLEA9_A015659 | 0.000 | 0.477 | 2.074 | 0.007 | 0.000 | 0.021 | P85524 | *Actinidia deliciosa* | Kirola |
| OLEA9_A021856 | 0.275 | 0.957 | 1.543 | 0.353 | 0.153 | 0.311 | Q9FYB5 | *Arabidopsis thaliana* | Chaperone protein dnaJ 11, chloroplastic |
| OLEA9_A022057 | 0.000 | 0.151 | 8.052 | 0.000 | 0.000 | 0.049 | Q9FXA4 | *Arabidopsis thaliana* | U-box domain-containing protein 26 |
| OLEA9_A022151 | 0.283 | 2.148 | 7.225 | 0.046 | 0.028 | 0.114 | Q8H1D6 | *Arabidopsis thaliana* | Receptor-like cytosolic serine/threonine-protein kinase RBK1 |
| OLEA9_A022485 | 0.335 | 0.821 | 1.130 | 0.127 | 0.118 | 0.155 | Q9C9Z2 | *Arabidopsis thaliana* | Protein RETICULATA-RELATED 3, chloroplastic |
| OLEA9_A023468 | 0.258 | 0.865 | 1.021 | 0.182 | 0.055 | 0.295 | Q9FF29 | *Arabidopsis thaliana* | PR5-like receptor kinase |
| OLEA9_A031387 | 0.137 | 0.901 | 1.234 | 0.136 | 0.024 | 0.093 | Q9SD62 | *Arabidopsis thaliana* | Putative receptor-like protein kinase At3g47110 |
| OLEA9_A033141 | 0.183 | 1.557 | 1.297 | 0.066 | 0.111 | 0.025 | F4HTV4 | *Arabidopsis thaliana* | Receptor-like protein 14 |
| OLEA9_A042127 | 0.006 | 0.101 | 0.113 | 0.000 | 0.000 | 0.063 | Q0JEU6 | *Oryza sativa* subsp. *japonica* | G-type lectin S-receptor-like serine/threonine-protein kinase LECRK3 |
| OLEA9_A046627 | 0.285 | 0.122 | 0.118 | 0.068 | 0.083 | 0.029 | Q8GRU6 | *Lotus japonicus* | Leucine-rich repeat receptor-like kinase protein HAR1 |
| OLEA9_A052325 | 0.000 | 0.103 | 3.906 | 0.000 | 0.000 | 0.030 | Q8VWQ5 | *Arabidopsis thaliana* | Probable WRKY transcription factor 50 |
| OLEA9_A057231 | 0.035 | 1.231 | 0.294 | 0.000 | 0.000 | 0.000 | P85524 | *Actinidia deliciosa* | Kirola |
| OLEA9_A069169 | 0.007 | 0.190 | 0.178 | 0.000 | 0.000 | 0.010 | Q9ASS4 | *Arabidopsis thaliana* | Probably inactive leucine-rich repeat receptor-like protein kinase At5g48380 |
| OLEA9_A074684 | 0.067 | 0.479 | 0.395 | 0.020 | 0.000 | 0.000 | F4I9S3 | *Arabidopsis thaliana* | Receptor-like protein 9a |
| OLEA9_A080470 | 0.035 | 0.159 | 0.269 | 0.004 | 0.000 | 0.009 | Q6JN47 | *Solanum lycopersicum* | Receptor-like protein EIX1 |
| OLEA9_A080644 | 0.051 | 0.284 | 1.215 | 0.140 | 0.026 | 0.058 | P0DO21 | *Nicotiana benthamiana* | Probable aspartic proteinase GIP2 |
| OLEA9_A082808 | 0.001 | 0.074 | 0.084 | 0.004 | 0.003 | 0.010 | Q9FK76 | *Arabidopsis thaliana* | Subtilisin-like protease SBT5.6 |
| OLEA9_A082849 | 0.027 | 1.998 | 0.474 | 0.000 | 0.000 | 0.007 | Q9M0A5 | *Arabidopsis thaliana* | Gamma-glutamyl peptidase 3 |
| OLEA9_A087022 | 0.098 | 0.302 | 1.562 | 0.062 | 0.041 | 0.135 | Q8VYP5 | *Arabidopsis thaliana* | Probable protein S-acyltransferase 14 |
| OLEA9_A092051 | 0.144 | 0.345 | 0.743 | 0.100 | 0.108 | 0.088 | Q94KB7 | *Arabidopsis thaliana* | MLO-like protein 6 |
| OLEA9_A093660 | 0.176 | 0.045 | 0.027 | 0.000 | 0.000 | 0.000 | Q1MX30 | *Oryza sativa* subsp. *indica* | Receptor kinase-like protein Xa21 |
| OLEA9_A101742 | 0.095 | 0.347 | 0.348 | 0.135 | 0.340 | 0.061 | Q8VZF4 | *Arabidopsis thaliana* | Serine/threonine-protein kinase ZRK3 |
| OLEA9_A103366 | 0.376 | 0.788 | 1.003 | 0.089 | 0.072 | 0.085 | Q9M667 | *Arabidopsis thaliana* | Disease resistance protein RPP13 |
| OLEA9_A103542 | 0.082 | 0.269 | 0.360 | 0.044 | 0.008 | 0.122 | A2XQD3 | *Oryza sativa* subsp. *indica* | G-type lectin S-receptor-like serine/threonine-protein kinase LECRK2 |
| OLEA9_A104662 | 0.122 | 0.905 | 0.340 | 0.079 | 0.068 | 0.088 | Q6JN47 | *Solanum lycopersicum* | Receptor-like protein EIX1 |
| OLEA9_A105521 | 0.013 | 0.163 | 0.983 | 0.000 | 0.000 | 0.000 | O82752 | *Arabidopsis thaliana* | Protein DETOXIFICATION 49 |
| OLEA9_A106137 | 0.031 | 2.727 | 14.528 | 0.192 | 0.163 | 0.421 | P16273 | *Hordeum vulgare* | Pathogen-related protein |
| OLEA9_A107068 | 0.167 | 0.672 | 0.797 | 0.007 | 0.007 | 0.015 | Q9SRL2 | *Arabidopsis thaliana* | Receptor-like protein 34 |
| OLEA9_A109671 | 0.042 | 0.505 | 0.550 | 0.018 | 0.000 | 0.005 | F4HTV4 | *Arabidopsis thaliana* | Receptor-like protein 14 |
| OLEA9_A110616 | 0.042 | 0.000 | 0.000 | 0.000 | 0.000 | 0.000 | Q93Y09 | *Arabidopsis thaliana* | Serine carboxypeptidase-like 45 |
| OLEA9_A112443 | 0.790 | 1.825 | 9.147 | 0.190 | 0.142 | 0.230 | Q9SF86 | *Arabidopsis thaliana* | Serine/threonine-protein kinase PCRK1 |
| OLEA9_A113877 | 0.231 | 0.749 | 0.903 | 0.097 | 0.000 | 0.029 | O64973 | *Arabidopsis thaliana* | Disease resistance protein RPS5 |
| OLEA9_A120073 | 0.059 | 0.219 | 0.244 | 0.223 | 0.078 | 0.260 | Q8VZF4 | *Arabidopsis thaliana* | Serine/threonine-protein kinase ZRK3 |
| OLEA9_A122006 | 0.336 | 1.251 | 2.269 | 0.086 | 0.036 | 0.100 | Q9C6A6 | *Arabidopsis thaliana* | Receptor-like protein 13 |
| OLEA9_NCA002199 | 0.654 | 1.598 | 1.530 | 0.619 | 0.591 | 1.019 | Q9LRR4 | *Arabidopsis thaliana* | Putative disease resistance RPP13-like protein 1 |
| OLEA9_NCA025329 | 0.000 | 0.298 | 1.653 | 0.000 | 0.000 | 0.035 | P85524 | *Actinidia deliciosa* | Kirola |

(Supplemental Table 6 continues)

**Supplemental Table 7.** DEGs categorized as “DNA packaging and RNA processing”, as shown in **Figure 4D**.

| UniProt Gene Name | GAZ | MAZ | EAZ | GC | MC | EC | Uniprot Homolog | Annotation | |
| --- | --- | --- | --- | --- | --- | --- | --- | --- | --- |
| OLEA9_A001910 | 0.401 | 0.871 | 0.928 | 0.475 | 0.810 | 0.509 | F4HXV6 | *Arabidopsis thaliana* | Nuclear pore complex protein NUP155 |
| OLEA9_A004176 | 0.067 | 0.024 | 0.027 | 0.098 | 0.053 | 0.060 | P20825 | *Drosophila melanogaster* | Retrovirus-related Pol polyprotein from transposon 297 |
| OLEA9_A013192 | 0.000 | 0.036 | 0.018 | 0.001 | 0.004 | 0.013 | Q8I7P9 | *Drosophila melanogaster* | Retrovirus-related Pol polyprotein from transposon opus |
| OLEA9_A019395 | 1.129 | 0.568 | 0.457 | 1.192 | 0.724 | 0.675 | Q9FI49 | *Arabidopsis thaliana* | Pentatricopeptide repeat-containing protein At5g50990 |
| OLEA9_A022817 | 0.038 | 0.010 | 0.007 | 0.017 | 0.007 | 0.010 | P10978 | *Nicotiana tabacum* | Retrovirus-related Pol polyprotein from transposon TNT 1-94 |
| OLEA9_A026159 | 9.037 | 4.375 | 4.461 | 9.086 | 5.430 | 5.216 | Q94CJ5 | *Arabidopsis thaliana* | Protein RETICULATA-RELATED 4, chloroplastic |
| OLEA9_A042606 | 0.441 | 0.217 | 0.192 | 0.484 | 0.316 | 0.270 | P10978 | *Nicotiana tabacum* | Retrovirus-related Pol polyprotein from transposon TNT 1-94 |
| OLEA9_A055700 | 0.030 | 0.073 | 0.062 | 0.041 | 0.072 | 0.046 | Q94HW2 | *Arabidopsis thaliana* | Retrovirus-related Pol polyprotein from transposon RE1 |
| OLEA9_A057931 | 0.792 | 0.394 | 0.388 | 0.912 | 0.478 | 0.513 | Q3SZ85 | *Bos taurus* | Putative peptidyl-tRNA hydrolase PTRHD1 |
| OLEA9_A061746 | 0.167 | 0.464 | 0.701 | 0.175 | 0.276 | 0.221 | P10978 | *Nicotiana tabacum* | Retrovirus-related Pol polyprotein from transposon TNT 1-94 |
| OLEA9_A083271 | 0.034 | 0.105 | 0.185 | 0.040 | 0.042 | 0.082 | Q9LYW9 | *Arabidopsis thaliana* | DnaJ protein P58IPK homolog |
| OLEA9_A085197 | 0.036 | 0.100 | 0.180 | 0.044 | 0.129 | 0.084 | Q94HW2 | *Arabidopsis thaliana* | Retrovirus-related Pol polyprotein from transposon RE1 |
| OLEA9_A095151 | 0.085 | 0.188 | 0.260 | 0.065 | 0.096 | 0.113 | Q6E7H0 | *Arabidopsis thaliana* | Origin of replication complex subunit 3 |
| OLEA9_A095603 | 0.370 | 0.081 | 0.056 | 0.330 | 0.325 | 0.138 | Q9FZD4 | *Arabidopsis thaliana* | Putative pentatricopeptide repeat-containing protein At1g26500 |
| OLEA9_A098792 | 0.099 | 0.035 | 0.036 | 0.082 | 0.030 | 0.038 | Q94HW2 | *Arabidopsis thaliana* | Retrovirus-related Pol polyprotein from transposon RE1 |
| OLEA9_A107585 | 0.058 | 0.019 | 0.015 | 0.069 | 0.036 | 0.043 | Q99315 | *Saccharomyces cerevisiae (strain ATCC 204508 / S288c)* | Transposon Ty3-G Gag-Pol polyprotein |
| OLEA9_A112982 | 0.109 | 0.039 | 0.030 | 0.025 | 0.006 | 0.019 | P10978 | *Nicotiana tabacum* | Retrovirus-related Pol polyprotein from transposon TNT 1-94 |
| OLEA9_NCA006515 | 0.279 | 1.271 | 0.666 | 0.257 | 0.445 | 0.436 | P10978 | *Nicotiana tabacum* | Retrovirus-related Pol polyprotein from transposon TNT 1-94 |
| OLEA9_NCA006890 | 0.104 | 0.045 | 0.034 | 0.045 | 0.026 | 0.020 | Q94HW2 | *Arabidopsis thaliana* | Retrovirus-related Pol polyprotein from transposon RE1 |
| OLEA9_NCA009252 | 4.889 | 2.399 | 2.360 | 4.690 | 2.748 | 2.648 | Q94HW2 | *Arabidopsis thaliana* | Retrovirus-related Pol polyprotein from transposon RE1 |
| OLEA9_NCA015480 | 0.081 | 0.000 | 0.000 | 0.024 | 0.000 | 0.000 | F4IXE7 | *Arabidopsis thaliana* | Increased DNA methylation 1 |
| OLEA9_NCA018557 | 0.000 | 0.431 | 0.698 | 0.000 | 0.040 | 0.025 | P04323 | *Drosophila melanogaster* | Retrovirus-related Pol polyprotein from transposon 17.6 |
| OLEA9_A007653 | 0.612 | 2.138 | 5.776 | 0.041 | 0.019 | 0.234 | Q8L8Z8 | *Arabidopsis thaliana* | Monothiol glutaredoxin-S2 |
| OLEA9_A010838 | 0.338 | 0.126 | 0.121 | 0.138 | 0.144 | 0.188 | O57683 | *Xenopus laevis* | Splicing factor 3B subunit 1 |
| OLEA9_A013063 | 0.034 | 0.074 | 0.125 | 0.017 | 0.005 | 0.042 | P10978 | *Nicotiana tabacum* | Retrovirus-related Pol polyprotein from transposon TNT 1-94 |
| OLEA9_A020606 | 0.145 | 0.067 | 0.062 | 0.105 | 0.109 | 0.065 | P10978 | *Nicotiana tabacum* | Retrovirus-related Pol polyprotein from transposon TNT 1-94 |
| OLEA9_A024481 | 0.010 | 0.021 | 0.034 | 0.002 | 0.001 | 0.005 | P10978 | *Nicotiana tabacum* | Retrovirus-related Pol polyprotein from transposon TNT 1-94 |
| OLEA9_A033073 | 0.006 | 0.019 | 0.097 | 0.003 | 0.000 | 0.000 | P10978 | *Nicotiana tabacum* | Retrovirus-related Pol polyprotein from transposon TNT 1-94 |
| OLEA9_A051984 | 0.113 | 0.474 | 0.268 | 0.002 | 0.000 | 0.004 | Q94HW2 | *Arabidopsis thaliana* | Retrovirus-related Pol polyprotein from transposon RE1 |
| OLEA9_A052832 | 0.097 | 0.025 | 0.024 | 0.034 | 0.053 | 0.058 | Q9SY09 | *Arabidopsis thaliana* | Small nuclear ribonucleoprotein SmD1b |
| OLEA9_A065485 | 0.002 | 0.054 | 0.108 | 0.000 | 0.000 | 0.000 | Q94HW2 | *Arabidopsis thaliana* | Retrovirus-related Pol polyprotein from transposon RE1 |
| OLEA9_A071071 | 0.000 | 0.018 | 0.017 | 0.000 | 0.000 | 0.000 | P10978 | *Nicotiana tabacum* | Retrovirus-related Pol polyprotein from transposon TNT 1-94 |
| OLEA9_A073692 | 0.030 | 0.065 | 0.285 | 0.042 | 0.027 | 0.051 | P10978 | *Nicotiana tabacum* | Retrovirus-related Pol polyprotein from transposon TNT 1-94 |
| OLEA9_A081133 | 0.146 | 0.398 | 0.331 | 0.315 | 0.416 | 0.262 | Q9SF35 | *Arabidopsis thaliana* | 40S ribosomal protein S23-1 |
| OLEA9_A083104 | 0.000 | 0.053 | 0.032 | 0.000 | 0.000 | 0.015 | P10978 | *Nicotiana tabacum* | Retrovirus-related Pol polyprotein from transposon TNT 1-94 |
| OLEA9_A089421 | 0.037 | 0.082 | 0.154 | 0.128 | 0.101 | 0.055 | P10978 | *Nicotiana tabacum* | Retrovirus-related Pol polyprotein from transposon TNT 1-94 |
| OLEA9_NCA010236 | 0.008 | 0.038 | 0.028 | 0.009 | 0.017 | 0.009 | P20825 | *Drosophila melanogaster* | Retrovirus-related Pol polyprotein from transposon 297 |
| OLEA9_NCA012877 | 0.022 | 0.064 | 0.088 | 0.002 | 0.001 | 0.007 | Q8I7P9 | *Drosophila melanogaster* | Retrovirus-related Pol polyprotein from transposon opus |
| OLEA9_NCA014735 | 0.669 | 0.320 | 0.341 | 0.457 | 0.615 | 0.268 | Q94HW2 | *Arabidopsis thaliana* | Retrovirus-related Pol polyprotein from transposon RE1 |
| OLEA9_NCA015135 | 0.000 | 0.046 | 0.057 | 0.000 | 0.000 | 0.000 | P10978 | *Nicotiana tabacum* | Retrovirus-related Pol polyprotein from transposon TNT 1-94 |
| OLEA9_PT000044 | 6.639 | 2.001 | 3.379 | 2.501 | 2.701 | 1.757 | Q7FNS5 | *Atropa belladonna* | 50S ribosomal protein L20, chloroplastic |

(Supplemental Table 7 continues)

**Supplemental Table 8.** DEGs categorized as “housekeeping genes (including genes of chloroplast and mitochondria located enzymes, iron metabolism, and ubiquitin-proteasome pathway)”, as shown in **Figure 4D**.

| UniProt Gene Name | GAZ | MAZ | EAZ | GC | MC | EC | Uniprot Homolog | Annotation | |
| --- | --- | --- | --- | --- | --- | --- | --- | --- | --- |
| OLEA9_A021336 | 2.673 | 1.265 | 1.237 | 2.686 | 1.499 | 1.697 | Q9FJW4 | *Arabidopsis thaliana* | NADH dehydrogenase [ubiquinone] iron-sulfur protein 4, mitochondrial |
| OLEA9_A023811 | 2.422 | 5.035 | 5.843 | 1.896 | 2.720 | 3.398 | Q94BU8 | *Arabidopsis thaliana* | Guanosine deaminase |
| OLEA9_A043283 | 0.263 | 0.000 | 0.000 | 0.022 | 0.000 | 0.000 | Q9XGI7 | *Solanum lycopersicum* | Nicotianamine synthase |
| OLEA9_A053802 | 0.259 | 0.033 | 0.049 | 0.031 | 0.000 | 0.012 | Q84TF5 | *Arabidopsis thaliana* | Probable E3 ubiquitin-protein ligase RHA4A |
| OLEA9_A056253 | 6.769 | 17.278 | 14.753 | 7.666 | 14.487 | 8.602 | Q5JL96 | *Oryza sativa* subsp. *japonica* | Probable E3 ubiquitin-protein ligase RZFP34 |
| OLEA9_A064234 | 0.145 | 0.072 | 0.039 | 0.035 | 0.025 | 0.010 | O24457 | *Arabidopsis thaliana* | Pyruvate dehydrogenase E1 component subunit alpha-3, chloroplastic |
| OLEA9_A068765 | 0.217 | 0.030 | 0.046 | 0.127 | 0.117 | 0.043 | Q42961 | *Nicotiana tabacum* | Phosphoglycerate kinase, chloroplastic |
| OLEA9_A068798 | 2.034 | 0.523 | 0.831 | 1.856 | 0.895 | 1.294 | O81208 | *Arabidopsis thaliana* | Light-harvesting complex-like protein OHP1, chloroplastic |
| OLEA9_A070255 | 2.715 | 1.354 | 1.270 | 2.965 | 1.856 | 1.564 | O48721 | *Arabidopsis thaliana* | Uroporphyrinogen-III synthase, chloroplastic |
| OLEA9_A071923 | 2.048 | 0.945 | 0.852 | 2.197 | 1.555 | 1.251 | Q9LYR5 | *Arabidopsis thaliana* | Peptidyl-prolyl cis-trans isomerase FKBP19, chloroplastic |
| OLEA9_A072703 | 1.440 | 2.969 | 3.190 | 1.765 | 3.006 | 3.488 | Q9C5M0 | *Arabidopsis thaliana* | Mitochondrial dicarboxylate/tricarboxylate transporter DTC |
| OLEA9_A073535 | 0.153 | 0.006 | 0.029 | 0.188 | 0.091 | 0.052 | P29677 | *Solanum tuberosum* | Mitochondrial-processing peptidase subunit alpha |
| OLEA9_A075311 | 0.006 | 0.101 | 0.059 | 0.000 | 0.035 | 0.010 | P27643 | *Bacillus subtilis (strain 168)* | Stage V sporulation protein K |
| OLEA9_A076093 | 0.000 | 0.241 | 0.825 | 0.000 | 0.008 | 0.062 | Q84TG3 | *Arabidopsis thaliana* | E3 ubiquitin-protein ligase PUB23 |
| OLEA9_A079185 | 0.207 | 0.839 | 1.222 | 0.031 | 0.035 | 0.033 | Q9SGP6 | *Arabidopsis thaliana* | Glutaredoxin-C9 |
| OLEA9_A082869 | 0.132 | 0.026 | 0.000 | 0.024 | 0.000 | 0.000 | Q0WPS2 | *Arabidopsis thaliana* | Cytochrome b561 domain-containing protein At4g18260 |
| OLEA9_A083536 | 1.485 | 0.663 | 0.257 | 1.128 | 0.631 | 0.789 | Q8RWM7 | *Arabidopsis thaliana* | Protein COFACTOR ASSEMBLY OF COMPLEX C SUBUNIT B CCB3, chloroplastic |
| OLEA9_A083667 | 0.773 | 0.352 | 0.234 | 0.791 | 0.700 | 0.420 | P93033 | *Arabidopsis thaliana* | Fumarate hydratase 1, mitochondrial |
| OLEA9_A085403 | 0.731 | 0.337 | 0.365 | 0.451 | 0.333 | 0.313 | P0CD42 | *Saccharum officinarum* | NAD(P)H-quinone oxidoreductase subunit 2 A, chloroplastic |
| OLEA9_A085781 | 0.660 | 0.280 | 0.301 | 0.807 | 0.630 | 0.535 | Q9SM57 | *Pisum sativum* | Outer envelope pore protein 21, chloroplastic |
| OLEA9_A085839 | 0.535 | 1.215 | 1.156 | 0.182 | 0.214 | 0.344 | Q8LFY8 | *Arabidopsis thaliana* | RING-H2 finger protein ATL54 |
| OLEA9_A087831  (Supplemental Table 8 continues) | 0.537 | 0.103 | 0.020 | 0.075 | 0.000 | 0.011 | Q8VY10 | *Arabidopsis thaliana* | Probable ubiquitin-conjugating enzyme E2 24 |
| OLEA9_A088525 | 0.002 | 0.030 | 0.021 | 0.002 | 0.018 | 0.012 | Q9SEI3 | *Arabidopsis thaliana* | 26S proteasome regulatory subunit 10B homolog A |
| OLEA9_A089264 | 21.333 | 9.683 | 10.829 | 29.460 | 14.963 | 17.822 | O86013 | *Novosphingobium aromaticivorans* | 4-hydroxy-2-oxovalerate aldolase |
| OLEA9_A090423 | 5.333 | 2.508 | 2.460 | 6.118 | 3.032 | 3.823 | P80680 | *Zea mays* | Ferredoxin-thioredoxin reductase, variable chain |
| OLEA9_A092797 | 2.991 | 1.439 | 1.263 | 2.374 | 2.185 | 2.005 | Q9SR43 | *Arabidopsis thaliana* | Phytochromobilin:ferredoxin oxidoreductase, chloroplastic |
| OLEA9_A094254 | 0.705 | 0.184 | 0.232 | 0.231 | 0.096 | 0.126 | P93768 | *Nicotiana tabacum* | Probable 26S proteasome non-ATPase regulatory subunit 3 |
| OLEA9_A097874 | 0.709 | 0.337 | 0.241 | 0.593 | 0.384 | 0.410 | Q9FJW4 | *Arabidopsis thaliana* | NADH dehydrogenase [ubiquinone] iron-sulfur protein 4, mitochondrial |
| OLEA9_A100523 | 14.978 | 32.455 | 45.222 | 21.304 | 31.933 | 36.780 | Q7FAH2 | *Oryza sativa* subsp. *japonica* | Glyceraldehyde-3-phosphate dehydrogenase 2, cytosolic |
| OLEA9_A104392 | 5.716 | 2.741 | 2.729 | 4.225 | 2.320 | 2.354 | O82627 | *Antirrhinum majus* | Granule-bound starch synthase 1, chloroplastic/amyloplastic |
| OLEA9_A104768 | 0.587 | 0.286 | 0.295 | 0.813 | 0.469 | 0.418 | P0CG85 | *Nicotiana sylvestris* | Polyubiquitin |
| OLEA9_A106289 | 0.099 | 0.036 | 0.018 | 0.061 | 0.028 | 0.037 | Q9STT5 | *Arabidopsis thaliana* | ABC transporter A family member 7 |
| OLEA9_A108390 | 0.508 | 0.082 | 0.068 | 0.058 | 0.000 | 0.023 | Q9XGI7 | *Solanum lycopersicum* | Nicotianamine synthase |
| OLEA9_A114414 | 1.416 | 0.597 | 0.663 | 1.616 | 0.995 | 0.951 | none |  | PF01485:IBR domain, a half RING-finger domain |
| OLEA9_A115763 | 0.436 | 0.036 | 0.037 | 0.134 | 0.000 | 0.042 | O23344 | *Arabidopsis thaliana* | Ferredoxin C 1, chloroplastic |
| OLEA9_A119337 | 0.175 | 0.447 | 0.577 | 0.046 | 0.083 | 0.165 | Q9SP35 | *Arabidopsis thaliana* | Mitochondrial import inner membrane translocase subunit TIM17-2 |
| OLEA9_A121464 | 0.616 | 0.256 | 0.221 | 0.486 | 0.298 | 0.250 | Q8L649 | *Arabidopsis thaliana* | E3 ubiquitin-protein ligase BIG BROTHER |
| OLEA9_NCA006082 | 2.419 | 0.970 | 0.222 | 0.477 | 0.000 | 0.119 | O64763 | *Arabidopsis thaliana* | E3 ubiquitin-protein ligase ATL9 |
| OLEA9_NCA019175 | 0.209 | 0.055 | 0.072 | 0.163 | 0.092 | 0.095 | Q9AV81 | *Oryza sativa* subsp. *japonica* | Pre-mRNA-processing factor 19 |
| OLEA9_NCA019284 | 0.253 | 0.056 | 0.036 | 0.218 | 0.084 | 0.055 | Q06FL7 | *Pelargonium hortorum* | NAD(P)H-quinone oxidoreductase subunit 5, chloroplastic |
| OLEA9_A000445 | 0.015 | 0.140 | 0.568 | 0.000 | 0.000 | 0.000 | Q9FXE3 | *Arabidopsis thaliana* | SufE-like protein 2, chloroplastic |
| OLEA9_A015132 | 0.014 | 0.347 | 8.011 | 0.000 | 0.000 | 0.059 | Q9SVG5 | *Arabidopsis thaliana* | Berberine bridge enzyme-like 18 |
| OLEA9_A036189 | 0.008 | 0.158 | 0.610 | 0.000 | 0.000 | 0.000 | Q9LIG0 | *Arabidopsis thaliana* | Clavaminate synthase-like protein At3g21360 |
| OLEA9_A057840 | 0.287 | 2.332 | 0.946 | 0.021 | 0.016 | 0.079 | Q949S5 | *Arabidopsis thaliana* | F-box protein PP2-B11 |
| OLEA9_A060255 | 0.016 | 0.254 | 0.505 | 0.056 | 0.018 | 0.089 | Q9T075 | *Arabidopsis thaliana* | Protein RMD5 homolog |
| OLEA9_A064303 | 0.373 | 0.800 | 1.448 | 0.076 | 0.000 | 0.228 | none |  | PF13639:Ring finger domain |
| OLEA9_A076805 | 0.478 | 2.043 | 8.674 | 0.017 | 0.000 | 0.000 | Q8L8Z8 | *Arabidopsis thaliana* | Monothiol glutaredoxin-S2 |
| OLEA9_A078564 | 0.222 | 0.877 | 1.738 | 0.041 | 0.000 | 0.070 | Q8L8Z8 | *Arabidopsis thaliana* | Monothiol glutaredoxin-S2 |
| OLEA9_A088017 | 3.802 | 14.571 | 12.656 | 0.097 | 0.022 | 0.192 | Q8L8Z8 | *Arabidopsis thaliana* | Monothiol glutaredoxin-S2 |
| OLEA9_A088768 | 0.086 | 0.005 | 0.000 | 0.048 | 0.054 | 0.051 | Q9SU56 | *Arabidopsis thaliana* | L-galactono-1,4-lactone dehydrogenase, mitochondrial |
| OLEA9_A092537 | 0.015 | 0.420 | 0.316 | 0.000 | 0.000 | 0.000 | Q6K609 | *Oryza sativa* subsp. *japonica* | Glutaredoxin-C3 |
| OLEA9_A093386 | 0.251 | 0.980 | 2.904 | 0.265 | 0.114 | 0.421 | Q42342 | *Arabidopsis thaliana* | Cytochrome b5 isoform E |
| OLEA9_A093601 | 0.115 | 0.366 | 2.744 | 0.060 | 0.015 | 0.137 | Q6R567 | *Capsicum annuum* | E3 ubiquitin-protein ligase RMA1H1 |
| OLEA9_A095712 | 0.574 | 7.342 | 7.299 | 0.121 | 0.000 | 0.114 | Q9LPU9 | *Arabidopsis thaliana* | Vacuolar iron transporter homolog 1 |
| OLEA9_A097817 | 0.000 | 0.130 | 0.313 | 0.025 | 0.000 | 0.085 | Q8GWS3 | *Arabidopsis thaliana* | Heavy metal-associated isoprenylated plant protein 2 |
| OLEA9_A097884 | 0.538 | 1.380 | 3.970 | 0.041 | 0.000 | 0.085 | Q6NQE2 | *Arabidopsis thaliana* | Probable NAD(P)H dehydrogenase (quinone) FQR1-like 1 |
| OLEA9_A100371 | 0.018 | 0.906 | 0.355 | 0.000 | 0.000 | 0.000 | Q9SVG5 | *Arabidopsis thaliana* | Berberine bridge enzyme-like 18 |
| OLEA9_A106781 | 0.012 | 0.217 | 0.235 | 0.000 | 0.000 | 0.000 | Q9LIF1 | *Arabidopsis thaliana* | Monothiol glutaredoxin-S10 |
| OLEA9_A107461 | 0.000 | 0.201 | 0.128 | 0.000 | 0.000 | 0.007 | Q9SVG5 | *Arabidopsis thaliana* | Berberine bridge enzyme-like 18 |
| OLEA9_A117840 | 0.158 | 0.845 | 0.444 | 0.019 | 0.005 | 0.016 | Q94A73 | *Arabidopsis thaliana* | BTB/POZ domain-containing protein At5g66560 |
| OLEA9_A120980 | 0.194 | 2.112 | 1.149 | 2.807 | 1.426 | 2.440 | Q9FIU6 | *Arabidopsis thaliana* | Organelle RRM domain-containing protein 2, mitochondrial |
| OLEA9_NCA004656 | 0.000 | 0.350 | 0.210 | 0.000 | 0.000 | 0.000 | Q9SI09 | *Arabidopsis thaliana* | Probable E3 ubiquitin-protein ligase XERICO |
| OLEA9_NCA008829 | 0.864 | 2.125 | 2.137 | 0.033 | 0.000 | 0.000 | Q9LYC6 | *Arabidopsis thaliana* | Glutaredoxin-C11 |
| OLEA9_A037082 | 0.154 | 0.041 | 0.017 | 0.098 | 0.019 | 0.017 | Q9S713 | *Arabidopsis thaliana* | Serine/threonine-protein kinase STN7, chloroplastic |
| OLEA9_A049858 | 2.709 | 1.286 | 1.358 | 2.445 | 1.412 | 1.430 | Q9M2D2 | *Arabidopsis thaliana* | UPF0187 protein At3g61320, chloroplastic |

(Supplemental Table 8 continues)

**Supplemental Table 9.** Unknown DEGs (category in Figure 4D) that belong to the 733 DEGs specific to FAZ.

| UniProt Gene Name | GAZ | MAZ | EAZ | GC | MC | EC |
| --- | --- | --- | --- | --- | --- | --- |
| OLEA9_A003539 | 0.261 | 0.081 | 0.076 | 0.041 | 0.000 | 0.000 |
| OLEA9_A004863 | 0.377 | 0.077 | 0.069 | 0.296 | 0.159 | 0.069 |
| OLEA9_A006492 | 1.592 | 0.501 | 0.510 | 1.789 | 0.964 | 0.899 |
| OLEA9_A007365 | 2.694 | 0.863 | 0.544 | 3.459 | 2.503 | 1.740 |
| OLEA9_A008215 | 0.493 | 0.239 | 0.232 | 0.579 | 0.524 | 0.343 |
| OLEA9_A008988 | 3.071 | 1.409 | 1.378 | 2.994 | 1.741 | 1.584 |
| OLEA9_A010854 | 1.352 | 0.654 | 0.634 | 1.378 | 0.678 | 0.712 |
| OLEA9_A011311 | 2.735 | 1.329 | 1.270 | 3.284 | 2.546 | 2.823 |
| OLEA9_A011607 | 0.435 | 0.904 | 1.037 | 0.624 | 1.111 | 0.857 |
| OLEA9_A011864 | 1.467 | 0.619 | 0.736 | 1.854 | 0.906 | 1.272 |
| OLEA9_A012571 | 0.290 | 1.369 | 7.049 | 0.046 | 0.355 | 0.379 |
| OLEA9_A012656 | 0.760 | 0.246 | 0.251 | 0.061 | 0.000 | 0.000 |
| OLEA9_A013357 | 0.140 | 0.354 | 0.457 | 0.237 | 0.329 | 0.370 |
| OLEA9_A015781 | 1.272 | 0.492 | 0.607 | 0.976 | 0.519 | 0.694 |
| OLEA9_A017477 | 0.463 | 0.176 | 0.167 | 0.464 | 0.344 | 0.233 |
| OLEA9_A018019 | 2.288 | 5.986 | 5.392 | 2.625 | 5.058 | 4.662 |
| OLEA9_A020953 | 1.651 | 0.487 | 0.460 | 2.108 | 1.204 | 1.832 |
| OLEA9_A022054 | 0.441 | 0.098 | 0.107 | 0.148 | 0.000 | 0.000 |
| OLEA9_A023381 | 9.287 | 4.199 | 3.669 | 9.448 | 5.796 | 4.950 |
| OLEA9_A024780 | 0.472 | 1.781 | 4.279 | 0.279 | 0.502 | 0.496 |
| OLEA9_A028204 | 0.297 | 0.047 | 0.040 | 0.217 | 0.108 | 0.180 |
| OLEA9_A028377 | 0.363 | 0.122 | 0.126 | 0.162 | 0.087 | 0.132 |
| OLEA9_A028993 | 0.386 | 0.026 | 0.028 | 0.512 | 0.171 | 0.210 |
| OLEA9_A029275 | 19.271 | 9.288 | 9.659 | 25.847 | 17.073 | 19.179 |
| OLEA9_A031265 | 0.959 | 0.464 | 0.442 | 0.845 | 0.572 | 0.451 |
| OLEA9_A031576 | 0.106 | 0.007 | 0.014 | 0.032 | 0.000 | 0.000 |
| OLEA9_A032420 | 0.651 | 0.135 | 0.021 | 0.130 | 0.000 | 0.037 |
| OLEA9_A033838 | 3.564 | 1.564 | 1.613 | 3.373 | 2.591 | 1.950 |
| OLEA9_A034556 | 0.269 | 0.890 | 0.790 | 0.293 | 0.400 | 0.453 |
| OLEA9_A036562 | 0.070 | 2.030 | 9.577 | 0.030 | 0.030 | 0.169 |
| OLEA9_A040312  (Supplemental Table 9 continues) | 0.502 | 0.132 | 0.125 | 0.522 | 0.421 | 0.277 |
| OLEA9_A041467 | 2.338 | 0.925 | 1.061 | 3.025 | 1.547 | 1.548 |
| OLEA9_A044516 | 0.038 | 0.651 | 37.074 | 0.000 | 0.211 | 0.054 |
| OLEA9_A044678 | 0.610 | 0.041 | 0.046 | 0.531 | 0.275 | 0.096 |
| OLEA9_A045188 | 0.583 | 1.240 | 2.850 | 0.686 | 0.988 | 1.214 |
| OLEA9_A046894 | 2.051 | 0.908 | 0.665 | 2.023 | 1.517 | 1.026 |
| OLEA9_A047082 | 1.135 | 0.450 | 0.464 | 0.849 | 0.806 | 0.589 |
| OLEA9_A047662 | 0.231 | 1.062 | 0.993 | 0.000 | 0.080 | 0.056 |
| OLEA9_A048103 | 2.046 | 1.019 | 0.752 | 1.800 | 0.988 | 1.019 |
| OLEA9_A049536 | 15.402 | 6.036 | 4.089 | 20.497 | 16.615 | 10.244 |
| OLEA9_A050494 | 0.051 | 0.000 | 0.000 | 0.010 | 0.000 | 0.000 |
| OLEA9_A052441 | 2.026 | 0.969 | 0.977 | 2.155 | 1.791 | 1.316 |
| OLEA9_A055852 | 2.774 | 0.373 | 0.983 | 2.224 | 2.156 | 1.820 |
| OLEA9_A055877 | 0.254 | 0.599 | 1.029 | 0.143 | 0.270 | 0.176 |
| OLEA9_A057360 | 2.148 | 0.933 | 0.830 | 2.800 | 2.170 | 1.573 |
| OLEA9_A057409 | 0.104 | 0.296 | 0.479 | 0.008 | 0.016 | 0.047 |
| OLEA9_A059920 | 0.117 | 0.000 | 0.000 | 0.066 | 0.000 | 0.000 |
| OLEA9_A060825 | 0.120 | 0.034 | 0.021 | 0.110 | 0.069 | 0.090 |
| OLEA9_A063439 | 0.303 | 0.075 | 0.035 | 0.030 | 0.000 | 0.013 |
| OLEA9_A066751 | 0.283 | 0.056 | 0.048 | 0.374 | 0.187 | 0.146 |
| OLEA9_A066966 | 0.768 | 0.119 | 0.100 | 0.869 | 0.387 | 0.280 |
| OLEA9_A068077 | 2.185 | 0.922 | 0.956 | 1.871 | 0.911 | 0.975 |
| OLEA9_A068600 | 0.193 | 0.042 | 0.036 | 0.118 | 0.031 | 0.032 |
| OLEA9_A068749 | 0.003 | 0.050 | 0.324 | 0.021 | 0.021 | 0.038 |
| OLEA9_A068788 | 2.812 | 1.112 | 1.430 | 2.570 | 1.605 | 1.716 |
| OLEA9_A071992 | 0.743 | 0.206 | 0.075 | 0.077 | 0.000 | 0.000 |
| OLEA9_A072749 | 2.277 | 0.878 | 1.124 | 1.698 | 1.073 | 1.053 |
| OLEA9_A073454 | 0.515 | 0.204 | 0.241 | 0.451 | 0.430 | 0.277 |
| OLEA9_A076342 | 1.433 | 0.642 | 0.430 | 1.583 | 1.105 | 0.889 |
| OLEA9_A077358 | 0.277 | 0.130 | 0.119 | 0.186 | 0.115 | 0.140 |
| OLEA9_A077713 | 0.020 | 0.442 | 0.397 | 0.209 | 0.508 | 0.638 |
| OLEA9_A080683 | 0.120 | 0.016 | 0.011 | 0.081 | 0.030 | 0.047 |
| OLEA9_A080887  (Supplemental Table 9 continues) | 0.200 | 0.089 | 0.099 | 0.150 | 0.101 | 0.085 |
| OLEA9_A082432 | 1.570 | 3.609 | 4.442 | 0.534 | 0.572 | 0.794 |
| OLEA9_A082961 | 4.814 | 2.167 | 2.402 | 7.836 | 5.075 | 4.633 |
| OLEA9_A083284 | 2.654 | 1.285 | 1.120 | 2.554 | 1.417 | 1.367 |
| OLEA9_A084105 | 0.296 | 0.611 | 0.612 | 0.276 | 0.286 | 0.306 |
| OLEA9_A086365 | 0.558 | 0.131 | 0.081 | 0.438 | 0.125 | 0.171 |
| OLEA9_A086473 | 0.245 | 0.017 | 0.000 | 0.083 | 0.079 | 0.019 |
| OLEA9_A087270 | 0.050 | 0.200 | 0.168 | 0.061 | 0.126 | 0.122 |
| OLEA9_A087500 | 0.461 | 0.098 | 0.029 | 0.072 | 0.000 | 0.000 |
| OLEA9_A090616 | 0.333 | 0.025 | 0.006 | 0.039 | 0.000 | 0.000 |
| OLEA9_A092095 | 0.672 | 0.067 | 0.104 | 0.644 | 0.535 | 0.256 |
| OLEA9_A092251 | 0.839 | 0.274 | 0.088 | 0.110 | 0.000 | 0.000 |
| OLEA9_A092938 | 0.601 | 0.121 | 0.107 | 0.354 | 0.087 | 0.119 |
| OLEA9_A101906 | 0.104 | 0.031 | 0.021 | 0.102 | 0.033 | 0.045 |
| OLEA9_A102012 | 2.986 | 0.620 | 0.765 | 0.168 | 0.102 | 0.000 |
| OLEA9_A102171 | 2.221 | 0.844 | 1.098 | 2.141 | 1.060 | 1.340 |
| OLEA9_A104665 | 0.116 | 0.025 | 0.006 | 0.018 | 0.000 | 0.000 |
| OLEA9_A105250 | 8.332 | 4.055 | 4.148 | 11.509 | 6.598 | 6.797 |
| OLEA9_A110987 | 2.960 | 1.488 | 1.382 | 3.405 | 2.236 | 1.713 |
| OLEA9_A114180 | 0.188 | 0.062 | 0.052 | 0.028 | 0.000 | 0.018 |
| OLEA9_A115169 | 0.649 | 1.430 | 1.730 | 0.501 | 0.563 | 0.946 |
| OLEA9_A115733 | 2.055 | 0.911 | 0.589 | 1.058 | 0.757 | 0.995 |
| OLEA9_A116565 | 0.141 | 0.045 | 0.037 | 0.152 | 0.074 | 0.095 |
| OLEA9_A117159 | 0.127 | 0.007 | 0.000 | 0.056 | 0.026 | 0.046 |
| OLEA9_A119023 | 0.286 | 0.103 | 0.089 | 0.133 | 0.068 | 0.038 |
| OLEA9_A120534 | 0.130 | 0.038 | 0.029 | 0.150 | 0.070 | 0.091 |
| OLEA9_NCA000523 | 0.598 | 0.199 | 0.269 | 0.478 | 0.339 | 0.233 |
| OLEA9_NCA001584 | 0.518 | 2.165 | 1.873 | 0.407 | 0.508 | 0.871 |
| OLEA9_NCA001965 | 0.641 | 0.152 | 0.054 | 0.505 | 0.443 | 0.112 |
| OLEA9_NCA002073 | 2.622 | 1.257 | 1.139 | 1.975 | 1.735 | 1.259 |
| OLEA9_NCA002074 | 2.075 | 0.328 | 0.642 | 1.274 | 0.522 | 1.028 |
| OLEA9_NCA002240 | 0.472 | 0.163 | 0.214 | 0.649 | 0.398 | 0.447 |
| OLEA9_NCA002326  (Supplemental Table 9 continues) | 0.009 | 0.116 | 0.745 | 0.010 | 0.072 | 0.097 |
| OLEA9_NCA002467 | 0.178 | 0.409 | 0.461 | 0.075 | 0.123 | 0.109 |
| OLEA9_NCA002753 | 0.390 | 0.023 | 0.000 | 0.107 | 0.000 | 0.000 |
| OLEA9_NCA003495 | 0.774 | 0.268 | 0.253 | 0.581 | 0.549 | 0.347 |
| OLEA9_NCA003633 | 0.455 | 0.127 | 0.151 | 0.466 | 0.386 | 0.188 |
| OLEA9_NCA003752 | 0.044 | 0.368 | 0.495 | 0.021 | 0.040 | 0.065 |
| OLEA9_NCA005866 | 0.092 | 0.017 | 0.018 | 0.099 | 0.031 | 0.066 |
| OLEA9_NCA005935 | 0.000 | 0.315 | 0.854 | 0.000 | 0.030 | 0.069 |
| OLEA9_NCA005975 | 2.312 | 0.839 | 1.020 | 2.394 | 1.320 | 1.663 |
| OLEA9_NCA006682 | 3.109 | 1.517 | 1.313 | 2.632 | 2.108 | 1.440 |
| OLEA9_NCA006973 | 1.185 | 0.573 | 0.441 | 0.850 | 0.447 | 0.504 |
| OLEA9_NCA007589 | 1.106 | 0.369 | 0.318 | 0.842 | 0.552 | 0.340 |
| OLEA9_NCA007645 | 0.581 | 0.239 | 0.227 | 0.520 | 0.338 | 0.319 |
| OLEA9_NCA009050 | 0.179 | 0.000 | 0.017 | 0.058 | 0.000 | 0.017 |
| OLEA9_NCA010807A | 0.010 | 0.141 | 0.110 | 0.006 | 0.054 | 0.007 |
| OLEA9_NCA010870 | 0.329 | 0.017 | 0.073 | 0.465 | 0.142 | 0.179 |
| OLEA9_NCA011321 | 0.133 | 0.027 | 0.010 | 0.084 | 0.023 | 0.031 |
| OLEA9_NCA011366 | 0.363 | 0.071 | 0.063 | 0.285 | 0.155 | 0.140 |
| OLEA9_NCA012152 | 0.081 | 0.301 | 0.439 | 0.154 | 0.213 | 0.278 |
| OLEA9_NCA013742 | 0.087 | 0.420 | 0.615 | 0.078 | 0.190 | 0.190 |
| OLEA9_NCA013951 | 2.179 | 0.746 | 1.019 | 2.032 | 1.479 | 1.987 |
| OLEA9_NCA014377 | 0.016 | 0.837 | 0.766 | 0.019 | 0.101 | 0.080 |
| OLEA9_NCA014509 | 0.242 | 0.102 | 0.080 | 0.227 | 0.190 | 0.216 |
| OLEA9_NCA014529 | 0.326 | 0.826 | 1.140 | 0.511 | 0.943 | 0.610 |
| OLEA9_NCA014892 | 2.230 | 1.049 | 1.103 | 2.216 | 1.291 | 1.104 |
| OLEA9_NCA017653 | 0.607 | 1.342 | 1.350 | 0.559 | 0.951 | 0.693 |
| OLEA9_NCA017931 | 1.447 | 0.286 | 0.128 | 1.290 | 0.561 | 0.677 |
| OLEA9_NCA018173 | 0.202 | 0.014 | 0.000 | 0.014 | 0.000 | 0.000 |
| OLEA9_NCA018176 | 0.379 | 1.057 | 0.785 | 0.313 | 0.578 | 0.329 |
| OLEA9_NCA018310 | 0.000 | 1.937 | 0.272 | 0.000 | 0.051 | 0.053 |
| OLEA9_NCA018659 | 0.127 | 0.923 | 1.297 | 0.000 | 0.027 | 0.139 |
| OLEA9_NCA019553 | 2.302 | 1.122 | 0.909 | 2.782 | 1.508 | 1.762 |
| OLEA9_NCA019650  (Supplemental Table 9 continues) | 0.419 | 0.062 | 0.037 | 0.030 | 0.000 | 0.000 |
| OLEA9_NCA019788 | 0.033 | 0.401 | 0.229 | 0.127 | 0.276 | 0.131 |
| OLEA9_NCA020091 | 0.147 | 0.397 | 0.354 | 0.120 | 0.234 | 0.191 |
| OLEA9_NCA020323 | 0.451 | 0.176 | 0.121 | 0.406 | 0.243 | 0.172 |
| OLEA9_NCA021213 | 1.011 | 0.243 | 0.348 | 1.055 | 0.589 | 0.539 |
| OLEA9_NCA021586 | 0.277 | 0.020 | 0.000 | 0.035 | 0.000 | 0.000 |
| OLEA9_NCA021790 | 0.420 | 0.029 | 0.000 | 0.338 | 0.059 | 0.000 |
| OLEA9_NCA022578 | 0.161 | 0.720 | 1.708 | 0.027 | 0.041 | 0.181 |
| OLEA9_NCA022704 | 0.504 | 0.077 | 0.057 | 0.165 | 0.000 | 0.029 |
| OLEA9_NCA022708 | 0.443 | 0.132 | 0.151 | 0.283 | 0.235 | 0.120 |
| OLEA9_NCA023268 | 0.346 | 0.000 | 0.000 | 0.161 | 0.040 | 0.039 |
| OLEA9_NCA023917 | 1.491 | 0.689 | 0.570 | 1.054 | 0.504 | 0.429 |
| OLEA9_NCA025639 | 0.664 | 0.266 | 0.191 | 0.690 | 0.354 | 0.350 |
| OLEA9_A000080 | 0.785 | 2.746 | 16.143 | 0.263 | 0.000 | 0.731 |
| OLEA9_A002684 | 0.287 | 0.129 | 0.114 | 0.028 | 0.000 | 0.041 |
| OLEA9_A003248 | 1.426 | 0.628 | 0.662 | 1.008 | 0.699 | 1.090 |
| OLEA9_A007587 | 0.178 | 1.172 | 1.692 | 0.034 | 0.000 | 0.083 |
| OLEA9_A009959 | 0.760 | 2.404 | 2.276 | 0.423 | 0.359 | 0.654 |
| OLEA9_A013077 | 0.107 | 0.431 | 0.723 | 0.013 | 0.003 | 0.023 |
| OLEA9_A013564 | 0.407 | 1.324 | 0.961 | 0.659 | 0.645 | 0.959 |
| OLEA9_A014162 | 0.152 | 1.383 | 3.287 | 0.120 | 0.000 | 1.553 |
| OLEA9_A020026 | 0.000 | 0.102 | 0.739 | 0.000 | 0.000 | 0.111 |
| OLEA9_A020056 | 0.007 | 0.164 | 1.508 | 0.010 | 0.000 | 0.003 |
| OLEA9_A020152 | 0.564 | 1.837 | 2.589 | 0.033 | 0.007 | 0.089 |
| OLEA9_A023951 | 0.066 | 0.778 | 1.370 | 0.268 | 0.431 | 0.069 |
| OLEA9_A025639 | 0.165 | 0.392 | 1.243 | 0.012 | 0.000 | 0.021 |
| OLEA9_A028297 | 0.442 | 2.136 | 1.630 | 0.084 | 0.013 | 0.058 |
| OLEA9_A038471 | 0.011 | 0.035 | 0.073 | 0.008 | 0.006 | 0.004 |
| OLEA9_A039530 | 0.000 | 0.188 | 0.229 | 0.111 | 0.022 | 0.431 |
| OLEA9_A042782 | 0.076 | 1.288 | 1.576 | 0.011 | 0.000 | 0.037 |
| OLEA9_A043033 | 0.132 | 0.047 | 0.045 | 0.003 | 0.000 | 0.018 |
| OLEA9_A046405 | 0.000 | 0.106 | 0.101 | 0.000 | 0.007 | 0.000 |
| OLEA9_A048394  (Supplemental Table 9 continue) | 0.172 | 0.049 | 0.012 | 0.063 | 0.099 | 0.081 |
| OLEA9_A048562 | 0.735 | 2.430 | 5.683 | 0.240 | 0.085 | 0.462 |
| OLEA9_A052144 | 0.287 | 0.670 | 1.107 | 0.559 | 0.394 | 0.306 |
| OLEA9_A068815 | 0.011 | 0.496 | 0.935 | 0.000 | 0.000 | 0.000 |
| OLEA9_A071740 | 0.077 | 0.653 | 0.827 | 0.038 | 0.052 | 0.012 |
| OLEA9_A072952 | 0.045 | 0.688 | 0.373 | 0.000 | 0.000 | 0.138 |
| OLEA9_A073928 | 0.002 | 0.045 | 0.099 | 0.000 | 0.000 | 0.000 |
| OLEA9_A074104 | 0.253 | 0.033 | 0.000 | 0.000 | 0.000 | 0.000 |
| OLEA9_A074259 | 0.196 | 2.009 | 0.872 | 0.040 | 0.028 | 0.108 |
| OLEA9_A079423 | 0.000 | 0.107 | 4.588 | 0.000 | 0.000 | 0.000 |
| OLEA9_A080523 | 0.085 | 1.120 | 4.087 | 0.094 | 0.037 | 0.042 |
| OLEA9_A081930 | 0.000 | 0.148 | 0.393 | 0.033 | 0.021 | 0.089 |
| OLEA9_A082853 | 0.108 | 0.677 | 0.439 | 0.076 | 0.051 | 0.076 |
| OLEA9_A083969 | 0.507 | 6.880 | 2.434 | 0.018 | 0.000 | 0.169 |
| OLEA9_A085227 | 0.177 | 0.500 | 1.058 | 0.093 | 0.010 | 0.281 |
| OLEA9_A085319 | 0.255 | 0.856 | 6.699 | 0.093 | 0.030 | 0.029 |
| OLEA9_A085936 | 0.058 | 0.216 | 0.246 | 0.031 | 0.000 | 0.040 |
| OLEA9_A087505 | 0.682 | 2.610 | 2.587 | 0.491 | 0.266 | 0.688 |
| OLEA9_A088965 | 0.167 | 0.583 | 0.784 | 0.090 | 0.023 | 0.272 |
| OLEA9_A091674 | 0.079 | 0.465 | 5.847 | 0.000 | 0.000 | 0.000 |
| OLEA9_A097256 | 0.097 | 1.625 | 15.448 | 0.027 | 0.000 | 0.000 |
| OLEA9_A097730 | 2.030 | 4.607 | 4.951 | 0.024 | 0.000 | 0.074 |
| OLEA9_A099319 | 0.269 | 0.039 | 0.007 | 0.000 | 0.000 | 0.000 |
| OLEA9_A099834 | 0.101 | 0.279 | 0.236 | 0.045 | 0.028 | 0.024 |
| OLEA9_A101531 | 0.048 | 0.178 | 0.403 | 0.044 | 0.023 | 0.006 |
| OLEA9_A104005 | 0.021 | 0.364 | 0.560 | 0.023 | 0.004 | 0.095 |
| OLEA9_A106592 | 0.234 | 0.736 | 5.100 | 0.055 | 0.000 | 0.185 |
| OLEA9_A110392 | 2.458 | 1.087 | 0.953 | 2.131 | 1.457 | 2.573 |
| OLEA9_A112591 | 0.176 | 2.853 | 4.701 | 0.053 | 0.000 | 0.000 |
| OLEA9_A113158 | 1.190 | 0.549 | 0.607 | 0.393 | 0.225 | 0.407 |
| OLEA9_A113473 | 0.011 | 0.385 | 0.280 | 0.000 | 0.000 | 0.000 |
| OLEA9_A115112 | 0.127 | 0.531 | 31.344 | 0.115 | 0.014 | 0.050 |
| OLEA9_A115829  (Supplemental Table 9 continue) | 0.204 | 0.921 | 1.811 | 0.117 | 0.000 | 0.314 |
| OLEA9_A116831 | 0.663 | 2.762 | 7.209 | 0.114 | 0.024 | 0.059 |
| OLEA9_A118366 | 0.012 | 1.040 | 2.234 | 0.079 | 0.066 | 0.340 |
| OLEA9_A119601 | 2.093 | 7.335 | 5.277 | 3.556 | 3.383 | 4.854 |
| OLEA9_NCA000044 | 0.891 | 0.370 | 0.217 | 0.272 | 0.280 | 0.217 |
| OLEA9_NCA000291 | 0.224 | 1.084 | 0.732 | 0.072 | 0.000 | 0.000 |
| OLEA9_NCA000999 | 0.074 | 0.564 | 0.322 | 0.013 | 0.000 | 0.089 |
| OLEA9_NCA001860 | 0.000 | 0.217 | 2.660 | 0.000 | 0.000 | 0.000 |
| OLEA9_NCA002393 | 0.316 | 0.000 | 0.035 | 0.000 | 0.000 | 0.000 |
| OLEA9_NCA004360 | 0.000 | 1.898 | 1.568 | 0.000 | 0.000 | 0.025 |
| OLEA9_NCA004361 | 0.000 | 4.698 | 6.795 | 0.000 | 0.000 | 0.000 |
| OLEA9_NCA005116 | 0.059 | 0.403 | 1.314 | 0.039 | 0.024 | 0.116 |
| OLEA9_NCA005144 | 0.183 | 1.471 | 1.323 | 0.000 | 0.000 | 0.062 |
| OLEA9_NCA005145 | 0.595 | 1.785 | 1.678 | 0.040 | 0.025 | 0.239 |
| OLEA9_NCA006019 | 0.000 | 0.274 | 0.756 | 0.000 | 0.000 | 0.124 |
| OLEA9_NCA006272 | 0.000 | 0.160 | 3.578 | 0.000 | 0.000 | 0.034 |
| OLEA9_NCA006675 | 0.000 | 0.223 | 0.423 | 0.000 | 0.000 | 0.000 |
| OLEA9_NCA006677 | 0.006 | 0.170 | 0.370 | 0.000 | 0.000 | 0.014 |
| OLEA9_NCA008613 | 0.316 | 0.044 | 0.016 | 0.000 | 0.000 | 0.000 |
| OLEA9_NCA009314 | 0.146 | 0.870 | 1.534 | 0.063 | 0.020 | 0.274 |
| OLEA9_NCA009315 | 0.043 | 0.443 | 1.199 | 0.000 | 0.000 | 0.055 |
| OLEA9_NCA010015 | 0.034 | 0.423 | 0.316 | 0.160 | 0.265 | 0.124 |
| OLEA9_NCA010815 | 0.353 | 2.125 | 5.757 | 0.218 | 0.091 | 0.814 |
| OLEA9_NCA011778 | 0.011 | 0.090 | 0.075 | 0.000 | 0.000 | 0.027 |
| OLEA9_NCA014404 | 0.071 | 0.640 | 0.393 | 0.040 | 0.020 | 0.060 |
| OLEA9_NCA015140 | 0.230 | 1.301 | 5.194 | 0.081 | 0.025 | 0.060 |
| OLEA9_NCA017690 | 0.000 | 0.143 | 0.468 | 0.030 | 0.000 | 0.143 |
| OLEA9_NCA017909 | 0.052 | 0.279 | 0.415 | 0.213 | 0.149 | 0.140 |
| OLEA9_NCA018366 | 0.073 | 0.370 | 0.534 | 0.106 | 0.052 | 0.064 |
| OLEA9_NCA018508 | 0.018 | 0.328 | 0.388 | 0.063 | 0.000 | 0.294 |
| OLEA9_NCA018680 | 1.475 | 0.581 | 0.491 | 0.694 | 0.646 | 0.775 |
| OLEA9_NCA019468 | 0.166 | 0.761 | 0.607 | 0.000 | 0.000 | 0.130 |
| OLEA9_NCA019967 | 0.025 | 0.416 | 0.295 | 0.000 | 0.000 | 0.011 |
| OLEA9_NCA019990 | 0.025 | 0.430 | 3.070 | 0.000 | 0.026 | 0.000 |
| OLEA9_NCA020102 | 2.166 | 0.648 | 1.039 | 2.482 | 2.751 | 1.656 |
| OLEA9_NCA020585 | 0.010 | 0.175 | 0.141 | 0.033 | 0.176 | 0.024 |
| OLEA9_NCA021801 | 0.230 | 1.006 | 1.185 | 0.223 | 0.220 | 0.334 |
| OLEA9_NCA023228 | 0.019 | 0.527 | 0.979 | 0.061 | 0.000 | 0.048 |
| OLEA9_NCA023457 | 0.009 | 0.118 | 0.112 | 0.028 | 0.000 | 0.000 |
| OLEA9_NCA024954 | 0.117 | 0.479 | 0.880 | 0.050 | 0.000 | 0.119 |
| OLEA9_NCA025431 | 0.125 | 0.627 | 1.307 | 0.077 | 0.000 | 0.025 |

(Supplemental Table 9 continues)
